# *De novo* assembly, annotation, and comparative analysis of 26 diverse maize genomes

**DOI:** 10.1101/2021.01.14.426684

**Authors:** Matthew B. Hufford, Arun S. Seetharam, Margaret R. Woodhouse, Kapeel M. Chougule, Shujun Ou, Jianing Liu, William A. Ricci, Tingting Guo, Andrew Olson, Yinjie Qiu, Rafael Della Coletta, Silas Tittes, Asher I. Hudson, Alexandre P. Marand, Sharon Wei, Zhenyuan Lu, Bo Wang, Marcela K. Tello-Ruiz, Rebecca D. Piri, Na Wang, Dong won Kim, Yibing Zeng, Christine H. O’Connor, Xianran Li, Amanda M. Gilbert, Erin Baggs, Ksenia V. Krasileva, John L. Portwood, Ethalinda K.S. Cannon, Carson M. Andorf, Nancy Manchanda, Samantha J. Snodgrass, David E. Hufnagel, Qiuhan Jiang, Sarah Pedersen, Michael L. Syring, David A. Kudrna, Victor Llaca, Kevin Fengler, Robert J. Schmitz, Jeffrey Ross-Ibarra, Jianming Yu, Jonathan I. Gent, Candice N. Hirsch, Doreen Ware, R. Kelly Dawe

## Abstract

We report *de novo* genome assemblies, transcriptomes, annotations, and methylomes for the 26 inbreds that serve as the founders for the maize nested association mapping population. The data indicate that the number of pan-genes exceeds 103,000 and that the ancient tetraploid character of maize continues to degrade by fractionation to the present day. Excellent contiguity over repeat arrays and complete annotation of centromeres further reveal the locations and internal structures of major cytological landmarks. We show that combining structural variation with SNPs can improve the power of quantitative mapping studies. Finally, we document variation at the level of DNA methylation, and demonstrate that unmethylated regions are enriched for cis-regulatory elements that overlap QTL and contribute to changes in gene expression.

**One sentence summary:** A multi-genome analysis of maize reveals previously unknown variation in gene content, genome structure, and methylation.

## Main text

Maize is the most widely planted crop in the world and an important model system for the study of gene function. The species is known for its extreme genetic diversity, which has allowed for broad adaptation throughout the tropics and intensive use in temperate regions. Much of its success can be attributed to a remarkable degree of heterosis when divergent inbred lines are crossed to make F1 hybrids. Nevertheless, most current genomic resources are referenced to a single inbred, B73. Yet prior data suggest the B73 genome contains only 63-74% of the genes and/or low-copy sequences in the full maize pan-genome (*1–4*). Moreover, there is extensive structural polymorphism in non-coding and regulatory genomic regions that has been shown to contribute to variation in numerous traits (*5*). In recent years, additional maize genomes have been assembled, allowing limited characterization of the species pan-genome and the extent of structural variation (*2, 6–10*). However, comparisons across genome projects are often confounded by differences in assembly and annotation methods.

The maize Nested Association Mapping (NAM) population was developed as a means to study the genetic architecture of quantitative traits (*11*). Twenty-five founder inbred lines were strategically selected to represent the breadth of maize diversity including lines from temperate, tropical, sweet corn, and popcorn germplasm (*12*). Each NAM parental inbred was crossed to B73 and selfed to generate 25 distinct populations of 200 recombinant inbred lines that combine the advantages of linkage and association mapping for important agronomic traits (*13*). Important biological infrastructure continues to be developed around these lines (e.g. (*14–16*)) but comprehensive genomic resources are needed to fully realize the power of the NAM population.

Here we describe the 25 assembled and annotated genomes for the NAM founder inbreds and an improved reference assembly of B73 (**Table S1**). In our comprehensive characterization of maize genomic diversity, we evaluate the maize pan-genome and its fractionation from a tetraploid ancestor, visualize the diversity of transposons and tandem repeat arrays, deploy enzymatic methyl-seq and ATAC-seq to characterize the pan-epigenome, and identify structural and epigenetic variation that impact phenotype.

### Consistency and quality of genome assemblies

The 26 genomes were sequenced to high depth (63-85X) using PacBio long-read technology, assembled into contigs using a hybrid approach (see Methods), corrected with long-read and Illumina short-read data, scaffolded using Bionano optical maps, and ordered into pseudomolecules using linkage data from the NAM recombinant inbred lines and maize pan-genome anchor markers (*4*). Assembly and annotation statistics far exceed nearly all available maize assemblies, with the total length of placed scaffolds (2.102-2.162Gb) at the estimated genome size of maize, a mean scaffold N50 of 119.2Mb (contig N50 of 25.7Mb), complete gene space (mean of 96% complete BUSCOs; (*17*)), and, based on the LTR Assembly Index (LAI, mean of 28; (*18*)), full assembly of the transposable-element-laden portions of the genome (**Table 1; Table S2**).

**Table 1:**
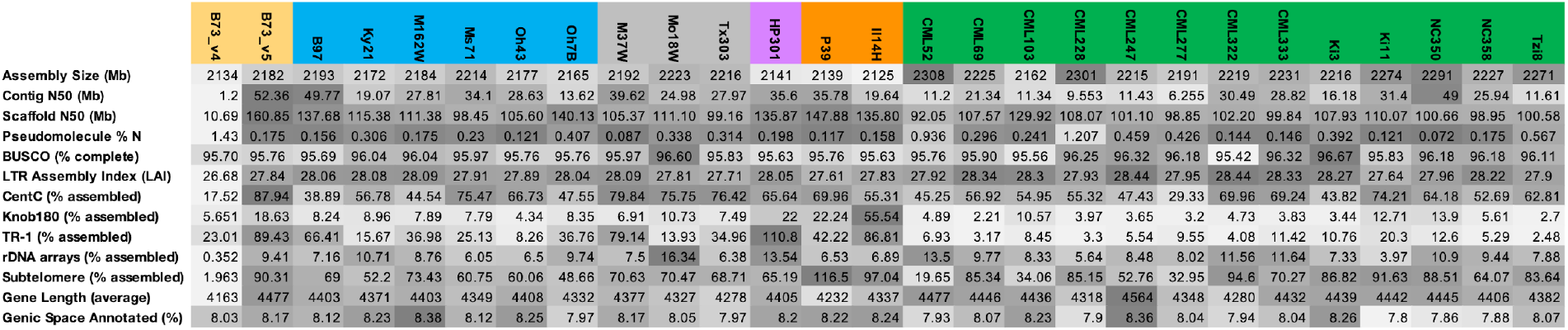
Quality metrics for genome assemblies and gene model annotations. Darker shading indicates higher quality.

### Gene identification and diversity in gene content

We sequenced mRNA from ten tissues in replicate for each inbred. These data were used as the basis for evidence-based gene annotation of each line, which was then improved using public B73 full-length cDNA and expressed sequence tags (ESTs). The evidence set was augmented with *ab initio* gene models and the gene structures uniformly refined for all accessions using phylogeny-based methods. This pipeline revealed an average of 40,621 (SE = 117) protein-coding and 4,998 (SE = 100) non-coding gene models per genome, with well over a million independent gene models generated across the 26 lines. Phylostrata analysis revealed that the great majority of genes share orthologs with species in the *Andropogoneae* tribe and grass family (**Fig. 1A**). The accuracy of the annotations, measured by the congruence between annotations and supporting evidence (Annotation Edit Distance, AED) (*19*), is substantially higher than previous reference maize and sorghum annotations (**Fig. S1**) (*2, 6, 10, 20–22*).

**Figure 1.**
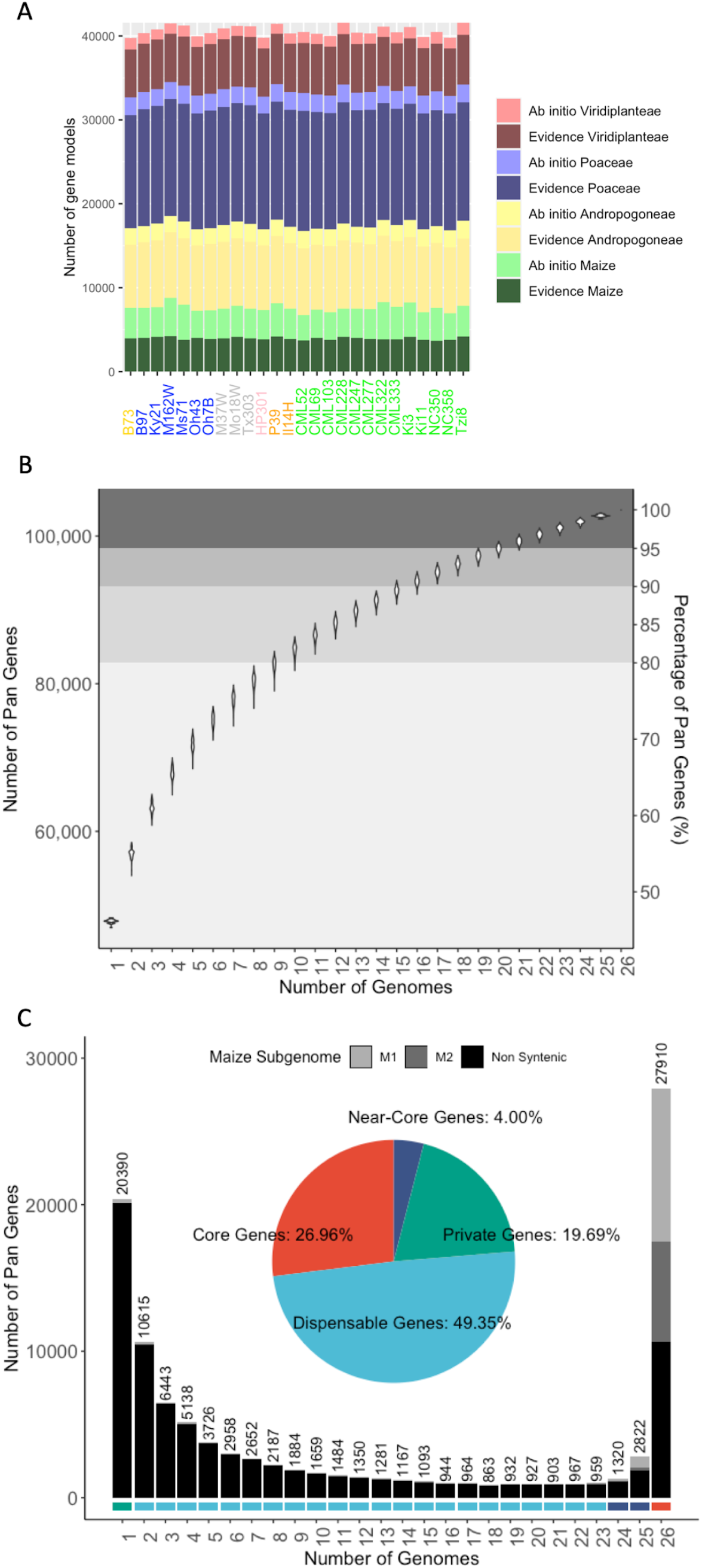
The gene space in the NAM genomes. **A**) Pan-genes categorized by annotation method and phylostrata. Genes annotated with evidence have mRNA support whereas *ab initio* genes are predicted based on DNA sequence alone. Genes within progressing phylostrata - species *Zea mays* (maize), tribe *Andropogoneae*, family *Poaceae*, kingdom *Viridiplantae* - are more conserved. **B**) The number of pan-genes added with each additional genome assembly. The error bars reflect different outcomes when the order of genomes was changed (the data were bootstrapped 1000 times). **C**) Frequency of pan-genes in the NAM genomes. The lower graph shows the number of genes present in only one genome (private), present in 2-23 genomes (dispensable), present in 24 or 25 genomes (near-core), and present in all 26 genomes (core). Grey shades indicate the proportion present in syntenic (M1 and M2 genomes) and non-syntenic positions. For B and C, tandem duplicates were considered as a single gene.

Based on the canonical transcripts from this complete set of annotations, we assessed the gene catalog of the pan-genome. Genes with high sequence similarity, located within blocks of homologous sequence in pairwise comparisons, were grouped together as one pan-gene. In many cases, a gene was not annotated by our computational pipeline in a particular inbred line, yet at least 90% of the gene was present in the correct homologous location; when this occurred, the pan-gene was considered present (**Fig. S2 A-B**; see Methods), even though in some cases the absence of annotation may be associated with fractionation and/or pseudogenization.

Across the 26 genomes, a total of 103,538 pan-genes were identified. Previous analysis of the maize pan-genome reported ∼63,000 pan-genes based on transcriptome assemblies of seedling RNA-seq reads from 500 individuals (*1*). The superior contiguity of our assemblies, as well as the application of both *ab initio* and evidence-based annotation using RNA-seq from a diverse set of ten tissues, likely accounts for the increased sensitivity here. Over 80% of pan-genes were identified within just ten inbred lines based on a bootstrap resampling of genomes; the rate of pan-gene increase as new genomes were added diminished beyond this point (**Fig. 1B**).

Pan-genes, excluding tandem duplicates, were classified as core (present in all 26 lines), near-core (present in 24-25 lines), dispensable (present in 2-23 lines), and private (present in only one line) (**Fig. 1C**). For each genotype, the portion of genes classified into each of these groups was consistent, with an average of 58.39% (SE = 0.07%) belonging to the core genome, 8.22% (SE = 0.05%) to the near-core genome, 31.75% (SE = 0.09%) to the dispensable genome, and 1.64% (SE = 0.08%) private genes (**Fig. 1C**; **Fig. S2 C-D; Table S3**). In total, there are 32,052 genes in the core/near-core portion of the pan-genome and 71,486 genes in the dispensable/private portion. The majority of core/near-core genes are syntenic to sorghum (57.8%) whereas this is rarely the case for dispensable/private genes (1.8% syntenic). Similarly, the core genes are generally from higher phylostrata levels (i.e. *Viridiplanteae* and *Poaceae*), while those in the near-core and dispensable sets either share orthologs only with closely related species or are maize-specific (**Fig. S2 F**). A total of 16,267 pan-genes had a putative tandem duplicate in at least one genome, of which 6,556 were found in a single genome. On a per gene basis in genomes with at least one tandem duplicate the average copy number is 2.20 (SE = 0.01) (**Fig. S2 E**).

### Partial tetraploidy and tempo of fractionation

The maize ancestor underwent a whole-genome duplication (WGD) allopolyploidy event 5-20 MYA ((*23, 24*), **Fig. 2A**). Evidence for WGD is found in the existence of two separate genomes that are broken and rearranged, yet still show clear synteny to sorghum (*23, 25*). Many duplicated genes have since undergone loss, or fractionation, reducing maize to its current diploid state (*25, 26*). Further, fractionation is biased towards one homoeologous genome (M2, more fractionated) over the other (M1, less fractionated) (*25*). The M1 and M2 subgenomes are composed almost exclusively of core (87.23%) and near-core (6.19%) pan-genes (**Figs. 1C, 2A**).

**Figure 2.**
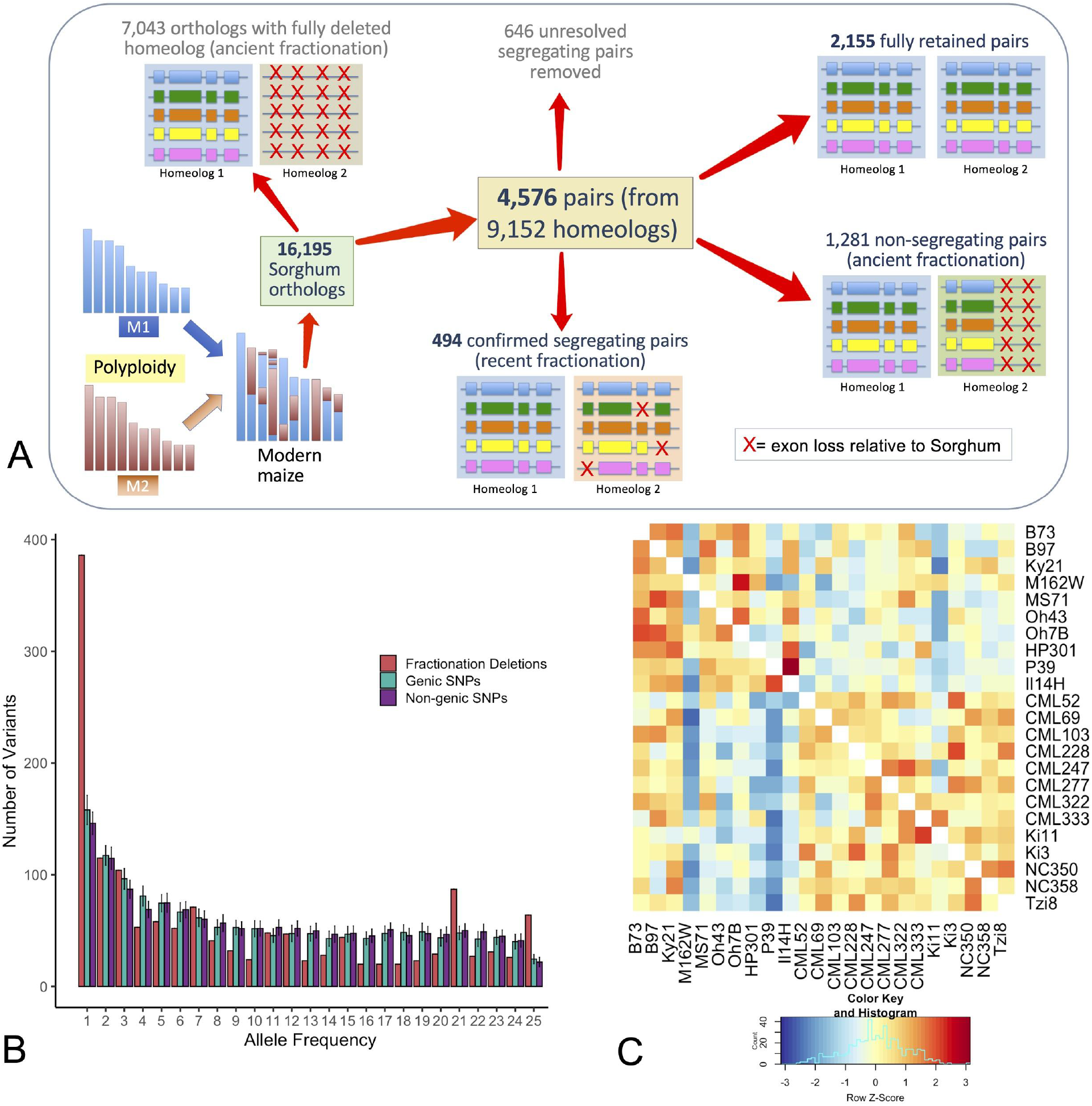
The tempo of fractionation in maize. **A**) Schematic showing how genes were categorized. 16,195 conservatively chosen orthologs were subdivided into classes representing retained pairs, ancient fractionation, and recent fractionation. **B**) Unfolded site frequency spectrum (SFS) of segregating exon loss and non-coding SNPs (genic and non-genic) using sorghum to define the ancestral state. **C**) Heatmap of the number of co-retained exons between any two NAM lines. Lines with mixed ancestry (M37W, Mo18W, Tx303) are excluded. Colors indicate the Z-score (the difference measured in standard deviations between a single pairwise comparison and all others in the row).

Given the ancient timeframe of the WGD in maize and the rapid tempo of fractionation observed in other species (*27, 28*), little variation in homoeolog retention is expected at the species level. In fact, prior work in temperate maize has suggested that most fractionation occurred long before maize was domesticated (*6, 29*). However, this diverse set of genomes allows for a more complete characterization of fractionation within the coalescence of the species. Since fractionation can occur at the level of small deletions (*26, 30*), we evaluated both partial and complete homoeolog loss beginning with a conservative set of 16,195 maize pan-orthologs. We determined that 7,043 were single-copy orthologs, where the homoeologous gene was likely deleted prior to maize speciation (**Fig. 2A**). Fractionation bias was substantial in this set, with 70% of single-copy orthologs retained in M1 and 30% retained in M2. In addition, we identified 4,576 homoeologous pairs (**Fig. 2A**) of which 2,155 had the same exon structure of the sorghum ortholog in both homoeologs. In 1,281 pairs, at least one copy of the gene differed from its sorghum ortholog, but did not vary among NAM lines, likely representing fractionation that pre-dated *Zea mays*. These ancient deletions were also biased toward M2, but much less substantially (9.4% deletion excess in M2), potentially reflecting different exon structure in the paleopolyploid progenitors. Another 1,140 pairs varied across the genomes in their pattern of exon retention, segregating for deletions or structural differences in at least one copy of the gene. This segregating set was manually curated (**Dataset S1**) to remove loci where exons or flanking sequence could not be confidently identified (**Fig. 2A**), resulting in a curated set of 494 homoeolog pairs segregating for fractionation, which represents more than 10% of the homoeologous pairs present in the pan-genome. Of these, 281 M2 homoeologs had exon loss compared to 236 M1 homoeologs, a 19% difference (p < 0.05, χ^2^ test), suggesting ongoing biased fractionation.

Coalescent theory predicts that segregating mutations, like the fractionation deletions identified, should have arisen within the last 4*N_e_* generations. If the effective population size in the maize progenitor teosinte is a reasonable upward bound for maize (*N_e_* = 150,000; (*31*)), we can infer that the majority of segregating neutral variation arose within the last 600,000 generations. Barring pervasive balancing selection for homoeologs, these data indicate that the majority of segregating fractionation substantially post-dates the last whole-genome duplication. Coalescent theory also predicts that rare deletions should be much younger than those segregating at intermediate frequency. We constructed the unfolded site frequency spectrum (SFS) of fractionation deletions in our curated set of homoeolog pairs and compared this to the unfolded SFS of non-coding SNPs using sorghum to define the ancestral state (**Fig. 2B**). The data reveal a similar frequency distribution in deletions and SNPs with a preponderance of rare variants in both, suggesting that a subset of fractionation may be quite young, potentially continuing in modern-day populations of maize. We also evaluated patterns of co-exon-retention in non-stiff-stalk temperate maize, tropical maize, and flint-derived maize, and observed clear evidence of population-specific fractionation (**Fig. 2C**). This surprising variation in homoeolog retention at the population level may reflect relaxed constraint following domestication and migration of maize to temperate climates.

Analysis of gene ontology terms revealed that fully retained homoeologous loci were enriched (p < 1×10^-05^) for DNA-binding, nucleic acid binding, phosphatase regulation, and transcription factor activity (consistent with prior results; (*32*), whereas segregating fractionated loci were enriched (p < 1×10^-05^) for transporter and catalytic activity (**Fig. S3, Dataset S1**). These results support the hypothesis that fractionated loci have distinct functions from those that are retained, presumably due to differential selection on multi-protein pathways or metabolic networks (*32, 33*).

### The repetitive fraction of the pan-genome

Transposable elements (TEs) were annotated in each assembly using both structural features and sequence homology (*34*). Individual TE libraries from each inbred were then combined to form a pan-genome library, which was used to identify TE sequences missed by individual libraries. The annotations reveal that DNA transposons and LTR retrotransposons comprise 8.5% and 74.4% of the genome, respectively (**Table S4, Fig. S4**). A total of 27,228 TE families were included in the pan-genome TE library, of which 59.7% were present in all 26 NAM founders and 2.5% were unique to one genome (**Fig. S5**). The average percentage of intact and fragmented TEs were 30.5% and 69.5% (SE = 0.06%), respectively. As reported previously, *Gypsy* LTR retrotransposon families are more abundant in pericentromeric regions, while *Copia* LTR retrotransposons are more abundant in the gene-dense chromosome arms (**Fig. S6**) (*35*). Tropical lines have significantly more *Gypsy* elements than temperate lines (p = 0.002, *t*-test), with mean *Gypsy* content of 1,018 Mbp and 988 Mbp, respectively (**Table S4, Fig. S4**). This may reflect increasing constraint on *Gypsy* proliferation in temperate lines that have, on average, smaller genomes (**Table 1**).

In some maize lines, over 15% of the genome is composed of tandem repeat arrays that include the centromere repeat CentC, the two knob repeats knob180 and TR-1, subtelomere, and telomere repeats (*36, 37*). Repeats of this type remain a major impediment to assembly. A mean of 60% of CentC, 70% of the 4-12-1 subtelomeric sequence (*38*)), 28.9% of TR-1, 1% of knob180, and 0.09% of rDNA repeat units were incorporated in the final assemblies (**Table 1**).

A total of 110 (of 260) functional centromeres identified by CENH3 ChIP-seq (*39, 40*) were fully assembled, and of these 88 are gapless ((**Fig. S7A** and (*40*)). Chromosomes with very long CentC arrays (such as chromosomes 1, 6, and 7) often have assembly gaps and the precise location of the centromere could not be determined. However many centromeres either have fully assembled small CentC arrays or the functional centromeres are located to one side of the CentC tracts in regions dominated by retrotransposons (**Fig. 3A**). By projecting all centromere locations onto B73, we were able to identify twelve centromere movement events (three on chr5 and chr9, and two on chr3, chr8 and chr10), clarifying and extending prior evidence for centromere shifting (*39*) (**Fig. 3B, Fig. S7B**). The variation in CentC abundance and positional polymorphism made it possible to gaplessly assemble at least two variants of all ten centromeres (**Fig. S7A**).

**Figure 3.**
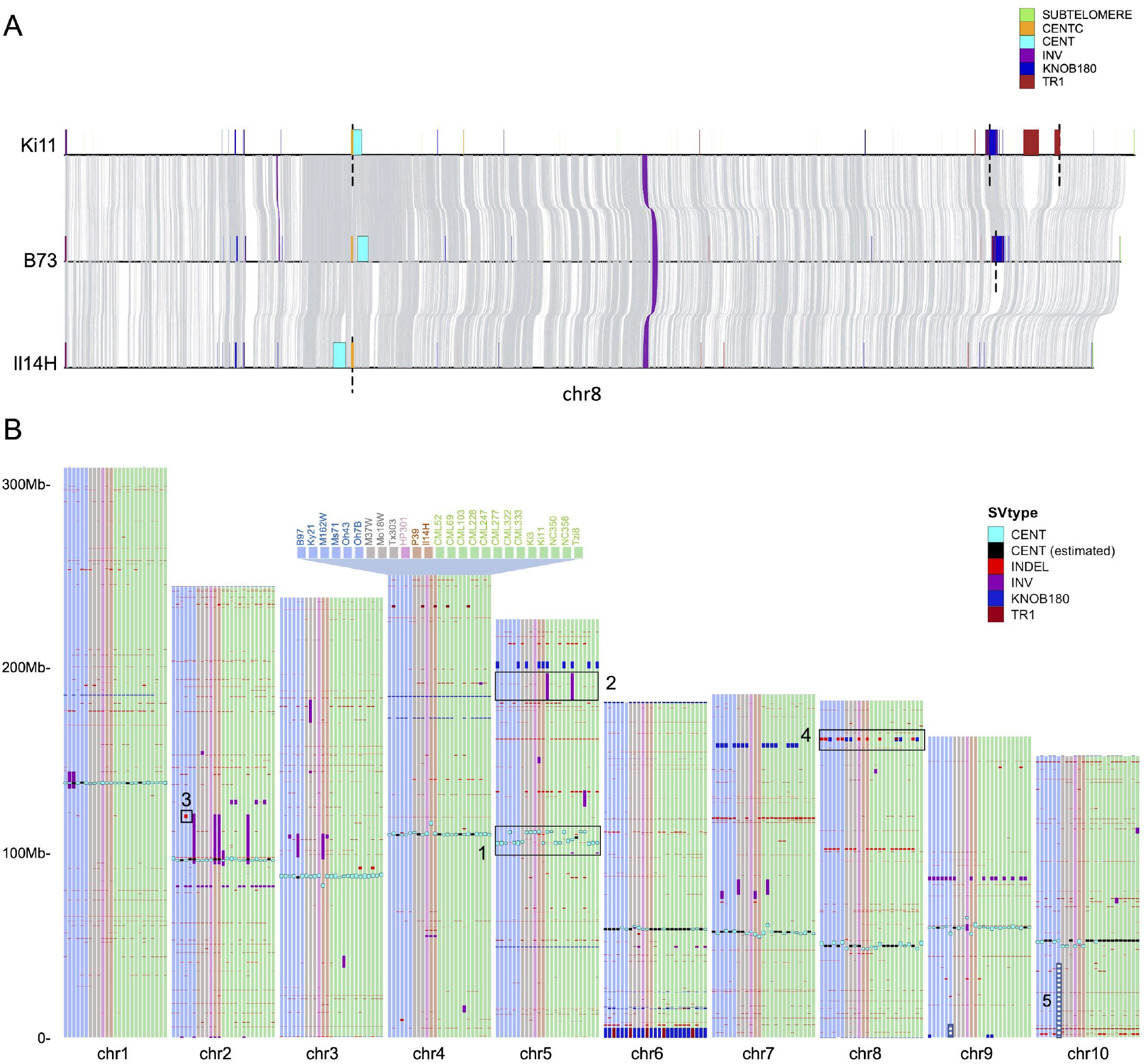
Structural variation in the NAM founders. **A**) Pairwise alignments between Ki11, B73, Il14H on chromosome 8. Grey links represent syntenic aligned regions; gaps of unknown size (scaffold gaps) are marked by dashed lines. **B**) Large (>100 kbp) structural variants, centromeres, and knobs across the NAM lines versus the B73 reference. The subset of SVs larger than 1 Mbp were manually curated, and only those containing genes are represented. Features 1-5 highlight major SVs: 1) Multiple centromere movement events; 2) A major inversion hypothesized to suppress recombination; 3) A large deletion in the Ms71 inbred; 4) Knob polymorphism; 5) Reciprocal translocation between chromosome 9 and 10 in the Oh7B inbred (both segments placed in their standard positions for display).

Both knob180 and TR-1 arrays are subject to meiotic drive and accumulate when a chromosome variant known as Abnormal chromosome 10 (Ab10) is present (*37, 41*). Although Ab10 is absent from modern inbreds, its legacy remains in the form of many large knobs. The majority of knob180 and TR-1 repeat arrays were identified in mid-arm positions (81.9%) where meiotic drive is most effective. Long knob180 and TR-1 repeat arrays can occur separately, but are more frequently intermingled in fragmented arrays along with transposons (**Fig. 3A, Fig. S8**) (*42*). Analysis of classical (cytologically visible) knobs on chromosome 1S, 2S, 2L, 3L, 4L, 5L, 6L, 7L, 8L, and 9S revealed that their locations are syntenic and that several are composed of a series of disjointed smaller knobs (**Fig. 3A, Fig. S9**). In some lines, knobs are not visible cytologically but can still be detected as smaller arrays at the sequence level; however, this is not always the case, as many show strict presence-absence variation among the NAM founder inbreds.

Tandem repeat arrays are also commonly found at the ends of chromosome arms (**Table S5**). Among the 520 chromosome ends, 57.9% contained knob180 repeats and 30.5% contained subtelomere repeats. At least 65.6% of the ends were fully assembled as indicated by the presence of telomere sequences.

### Structural variation and impact on phenotype

Comparative analyses among the NAM genotypes through mapping of long-reads to B73 revealed a cumulative total of 791,101 structural variants (SVs) greater than 100bp in size. Tropical lines, which are the most divergent NAM genomes from B73, include a substantially higher number of SVs than temperate lines (mean = 32,976 versus 29,742; p = 0.00013) **(Tables S6, S7).** Structural variants are more common on chromosome arms where recombination is highest (**Fig. S10**), similar to SNPs and other forms of genetic variation (*43*). Almost half (49.6%) of SVs were <5 kbp in size, with 25.7% being less than 500bp. Across all size classes SVs are skewed toward rare variants (**Fig. S11**). Several large SVs were found segregating within the 26 NAM genomes (**Fig. 3B**), including 35 distinct inversion polymorphisms and 5 insertion-deletion polymorphisms >1 Mbp. For example, a 14.6 Mbp inversion on chromosome 5 in the CML52 and CML322 lines, which was previously hypothesized based on suppressed recombination in the NAM RILs (*11*), is confirmed here based on assembly. Additionally, there is a 1.9 Mbp deletion with seven genes on chromosome 2 in the MS71 inbred, and a 1.8 Mbp deletion with two genes on chromosome 8 found in eight lines. Our data also capture a very large reciprocal translocation (involving >47 Mbp of DNA) between the short arms of chromosomes 9 and 10 in Oh7B that had been previously detected in cytological studies (*38*) (**Fig. 3B**).

The high proportion of rare SVs in maize suggests these may be a particularly deleterious class of variants, as observed in other species (*44, 45*). Indels and inversions occur in regions that have 49.8% fewer genic base pairs than the genomic background. Furthermore, SVs are 17% less likely to be found in conserved regions than SNPs (odds ratios of 0.27 and 0.58 for SVs and SNPs, respectively, Fisher’s Exact Test, p < 0.001). Approximate Bayesian computation modeling revealed that selection against SVs is at least as strong as that against nonsynonymous substitutions (**Fig. S12**; See Supplemental Methods). These results suggest that, when they occur, SVs are particularly consequential and are likely relevant to fitness.

To estimate the phenotypic impact of SVs, we assessed the genetic basis of 36 complex traits (*13*) using 71,196 filtered SVs in 4,027 recombinant inbred lines derived from the NAM founder inbreds (*11*) (**Fig. S13A**). The analysis revealed that SVs explain a high percentage of phenotypic variance for disease traits (60.10% ∼ 61.75%) and less for agronomic/morphological traits (20.04% ∼ 61.04%) and metabolic traits (4.79% ∼ 26.78%). Disease traits are often conferred by one or a few genes, whereas metabolic traits may be more sensitive to the environment and involve epistatic interactions that would not have been detected by our approach (*46*). Much of the phenotypic variation was also explained by SNPs, which were much more numerous (288-fold more) relative to our conservative set of SVs (**Fig. S13A**). When the SNP and SV data were integrated into one linear mixed model, the combined markers only slightly surpassed values from SNPs, consistent with the fact that most SVs are in high linkage disequilibrium with SNPs (**Fig. S13A**). We also carried out genome-wide association analyses (GWAS) to identify specific SVs contributing to phenotypic variation for the same suite of traits (**Fig. S13B-G**). Among the detected GWAS signals, 93.05% overlapped with those identified with SNPs and 6.95% were unique to SVs (no significant SNPs detected within 5 Mbp of significant SVs). The most significant association between a SV and a trait not identified using SNP markers was a QTL for northern leaf blight (NLB) on chromosome 10 (**Fig. S13F**). This SV is within a gene encoding a thylakoid lumenal protein; such proteins could be linked to plant immunity through the regulation of cell death during viral infection (*47*).

Disease resistance in plants is frequently associated with SV in the form of tandem arrays of resistance genes. Complex arrays of resistance genes are retained, potentially through birth-death dynamics in an evolutionary arms race with pathogens, or through balancing selection for the maintenance of diverse plant defenses (*48*). Nucleotide-binding, leucine-rich-repeat (NLR) proteins provide a common type of resistance. Our data reveal that there are fewer NLR genes in maize than other Poaceae (**Fig. S14**) and that most NAM lines have lost the same clades of NLRs as sorghum (**Fig. S15**). Only one line (CML277) retains the MIC1 NLR clade, which is particularly fast-evolving in Poaceae (*49*). Nevertheless, there is clear NLR variation among the NAM lines (**Fig. S16**), and tropical genomes contain a significantly higher number of NLR genes than temperate genomes (p=0.006), suggesting ongoing co-evolution with pathogens, particularly where disease pressure is high.

The annotated NLR genes were significantly enriched relative to random samples of genes for overlap with SVs (boot-strap permutation test, p<0.001). An extreme example is found at the *rp1* (resistance to *Puccinia sorghi1*) locus on the short arm of chromosome 10, which is known to be highly variable (*50*). We observed exceptional diversity in the NAM lines with as few as 4 *rp1* copies in P39, and as many as 30 in M37W (**Table S8**). However, due to its repetitive nature, only 18 NAM lines have gapless assemblies of the *rp1* locus.

SVs linked to transposons have been shown, through the modulation of gene expression, to underlie flowering-time adaptation in maize during tropical-to-temperate migration (*51, 52*). Our SV and TE-annotation pipelines identified the adaptive *CACTA*-like insertion previously reported upstream of the flowering-time locus *ZmCCT10* (*52*). We also surveyed an additional 173 genes linked to flowering-time (*53, 54*) and discovered three genes (*GL15*, *ZCN10*, and *Dof21*) with TE-derived SVs <5 kbp upstream of their transcription start sites. These SVs distinguish temperate from tropical lines (t < 7#x2212;2.346, p < 0.0358) (**Fig. S17**) and show significant correlation (F > 8.658, p < 0.001) with expression levels.

### Discovery of candidate cis-regulatory elements through DNA methylation

Based on sequence alone, it can be difficult to distinguish functional regulatory sequences from the multitude of non-functional and potentially deleterious genetic elements in the intergenic spaces. The problem is complicated by the fact that regulatory regions can be separated from their promoters by tens or hundreds of kilobases (*5, 55*). One way to identify functional regions is to score for unmethylated DNA, which provides both a tissue-independent indicator of gene regulatory elements and evidence that annotated genes are active (*5, 55, 56*). To incorporate DNA methylation to the NAM genomes resource, we sequenced enzymatic methyl-seq (EM-seq) libraries from each line and identified methylated bases in three sequence contexts, CG, CHG, and CHH (where H = A, T, or C). The results are consistent across genes and transposons, demonstrating the quality of the libraries (**Figs. S18, S19**). There is minor variation in total methylation across inbreds, with CML247 being noteworthy for uniformly lower CG methylation in several tissues (**Fig. S20**) pointing to the existence of a genetic variant that compromises mCG methylation in this line.

Each of the three methylation contexts reveal information on the locations of repeats, genes and regulatory elements. mCHH levels are generally low in maize except in heterochromatin borders, whereas mCHG is abundant in repetitive regions and depleted from regulatory elements and exons (**Fig. 4**) (*57*). mCG is also depleted from regulatory elements but can be abundant in exons, especially of broadly expressed genes (*58*). Thus, to identify unmethylated regions (UMRs) corresponding to both regulatory elements and gene bodies, we defined UMRs using a method that takes into account mCHG and mCG but does not exclude high mCG-only regions. Comparison of the 26 methylomes revealed uniformity in number and length of UMRs, averaging about 180 Mbp in total length in each genome (**Figs. S21, S22**). To confirm the accuracy of the UMR data, we also identified accessible chromatin regions (ACRs) using ATAC-seq for each inbred. We expect chromatin to be accessible mainly in the subset of genes expressed in the tissue sampled (primarily leaves) and to show a high level of concordance with UMRs. The data reveal that at least 98% of genic ACRs overlap with UMRs in each genome (**Fig. S23, S24**). For non-genic ACRs, the percent overlap was lower, but typically greater than 90%.

**Figure 4.**
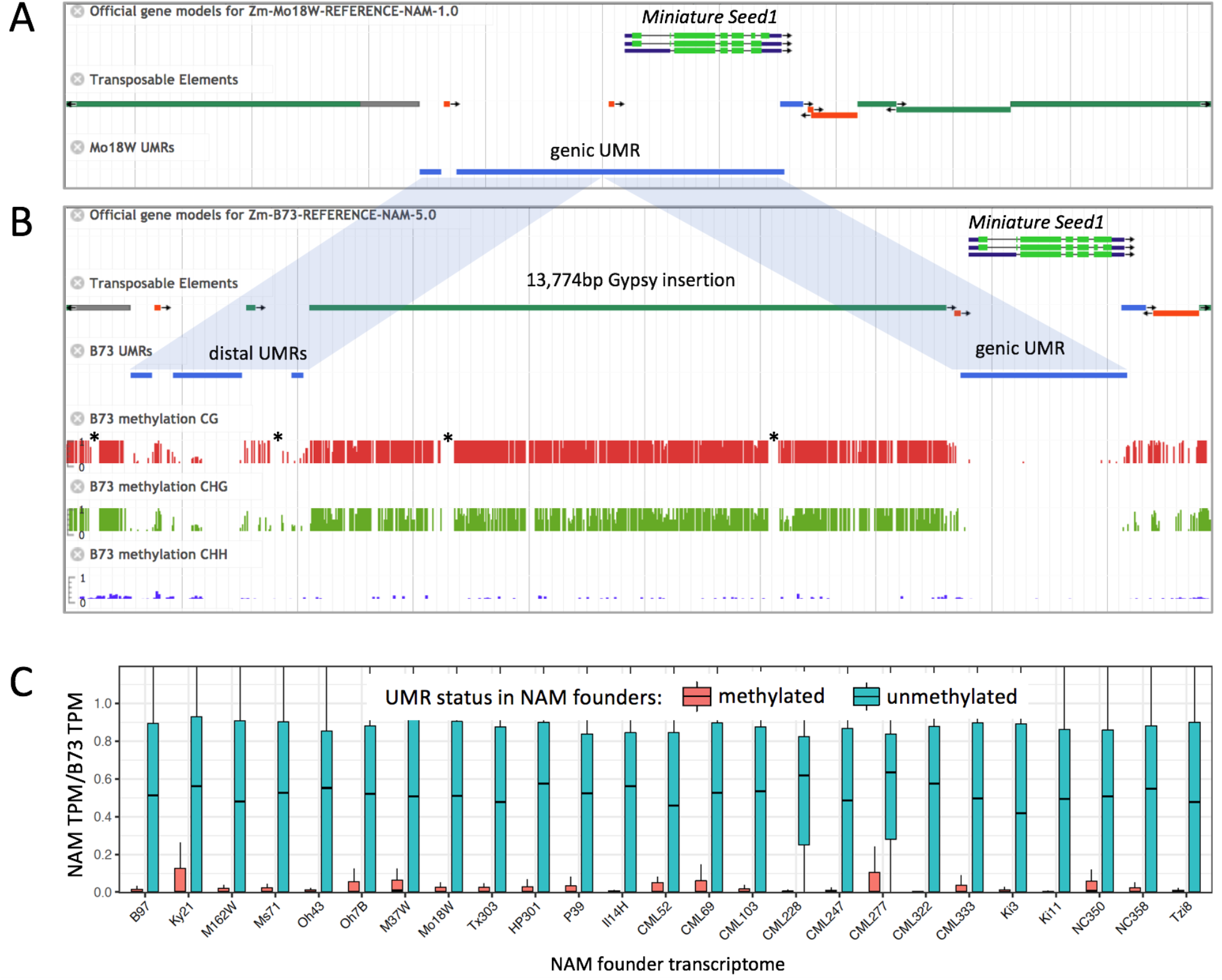
UMR variation across the NAM founders. **A**) Annotation of the *Miniature seed1* gene in the Mo17W inbred. An image from MaizeGDB browser shows gene, TE, and UMR tracks. TE tracks are color-coded by superfamily: green/grey = LTR, red = TIR, blue = LINE. The grey vertical lines show 2.5 kbp intervals. **B**) Annotation and underlying methylation data for *Miniature seed1* in the B73 inbred. The insertion of a *Gypsy* element moved part of the proximal UMR to a position 14 kbp upstream from the transcription start site (TSS). Methylation tracks indicate base-pair level methylation values from 0 to 100%. Asterisks indicate gaps in coverage, which are visible in separate tracks not shown here. **C**) Relationship between methylation and gene expression. UMRs were mapped to B73 to identify UMRs that overlap with TSS. The Y axis indicates the ratio of transcripts per million (TPM, compared to B73) when the region is methylated (red) or unmethylated (teal).

To assess methylation diversity, we mapped UMRs from all inbreds to the B73 genome. The data reveal that ∼95% of genic UMRs identified in one methylome overlap with a UMR in at least one other genome in pairwise comparisons (**Fig. S25**). UMR polymorphism is higher in the intergenic space, particularly among UMRs greater than 5 kbp from genes, where typically ∼75% of UMRs identified in one methylome overlap with a UMR identified in any other (**Fig, S25**). Even when the UMR sequence is conserved, its position relative to the closest gene may vary dramatically among inbreds. This is exemplified by the *Miniature Seed1* gene where a UMR proximal to the promoter in Mo18W is displaced nearly 14 kbp upstream in B73 by a single *Huck* element (*Gypsy* LTR superfamily) (**Fig. 4**). The *Huck* insertion is present in 23 of the 26 genomes, and in two of these (Oh43 and CML322), additional nested TE insertions increased the distance between the gene and the UMR to 27 kbp. Although the overlap of UMRs in pairs of lines is generally consistent with genetic distance across NAM lines (**Fig. S26**), UMRs from the inbred Tzi8 were not substantially shared with other tropical genomes. Tzi8 also has longer ACRs (**Fig. S24**) despite grouping well with other tropical lines in terms of gene expression patterns.

Variation in DNA methylation has been associated with adaptive traits in maize (*59*), most likely through effects on gene expression. To estimate how well UMRs predict transcriptional competency in these genomes, we identified a conservative subset of UMRs overlapping genes that were unmethylated in B73 but methylated in at least one other methylome. These differentially methylated regions were strongly correlated with gene expression in B73 and gene silencing in the other genomes (**Fig. 4, Fig. S27**). We further evaluated the enrichment of significant GWAS SNPs across 36 traits in UMRs. Based on genome-wide estimates, UMRs show 2.50- to 3.26-fold enrichment across traits for significant associations. Roughly 18% of SNPs identified by GWAS lie outside of genic regions but within UMRs (**Table S9**), consistent with the view that UMRs can be used to identify regulatory regions with important roles in determining phenotype (*5, 55, 56*).

## Summary

Our analysis of 26 genomes has uncovered previously unknown variation in both the genic and repetitive fractions of the pan-genome and provided new evidence of genome reorganization both before and after domestication. The available data will have broad utility for genetic and genomic studies and facilitate rapid associations to phenotyping information from the NAM lines. More generally, these new resources should motivate a shift away from the single reference mindset to a multi-reference view where any one of 26 inbreds, each with different experimental and agronomic advantages, can be deployed for the purposes of basic discovery and crop improvement. All data including annotations for genes, transposons, repeats, centromeres, UMRs, and ACRs are available with browser support at the maize community database, www.MaizeGDB.org.

## Acknowledgments

We appreciate the sequencing services provided by the University of Arizona, Oregon State University, Brigham Young University and the University of Georgia, as well as coordination among sequencing centers provided by Pacific Biosciences. The authors further acknowledge the High Performance Computing facility at Iowa State University (partially funded by NSF 1726447), Minnesota Supercomputing Institute, the Georgia Advanced Computing Resource Center, BlacknBlue high performance computing center at Cold Spring Harbor Laboratory and the participants of the Virtual Maize Annotation Jamboree who evaluated the initial gene predictions for benchmarking and improvements in the final gene annotations.

## Funding

Primary support for this work came from a generous grant from the National Science Foundation (IOS-1744001). Additional support came from NSF IOS-1546727 to CNH, USDA 2018-67013-27571 to CNH, USDA-ARS 8062-21000-041-00D and NSF IOS-1127112 to DW, NSF IOS-1546719 to MBH, NSF IOS-1822330 to JRI and MBH, USDA Hatch project CA-D-PLS-2066-H to JRI, NSF IOS-1856627 to RJS, an NSF Postdoctoral Fellowship in Biology DBI-1905869 to APM, NSF Graduate Research Fellowships 1650042 to AIH and 1744592 to SJS, NSF Research Traineeship (DGE-1545463) to Iowa State University (Trainee SJS), USDA-ARS 58-5030-8-064 to MBH and CMA, USDA-ARS project 5030-21000-068-00D to CMA and MW and NSF IOS-1546657 to JY

## Author contributions

Conceptualization – RKD, DW, MBH, CNH, JIG; Data curation – MW, AS, KMC, SO, JL, SW, ZL, BW, MKT-R, JLP, EKSC, CMA; Formal analysis – AS, MW, KMC, SO, JL, WAR, TG, AO, YQ, RDC, ST, AIH, SW, ZL, BW, MKT-R, RDP, YZ, CHO, XL, AMG, EB, JLP, NM, SJS, QJ, SP, MLS, KF, JIG; Funding acquisition - RKD, DW, MBH, CNH, JIG; Investigation – DK, DAK, APM, NW, DEH, VL, KF, JIG; Methodology – MBH, DW, CNH, AS, MW, KF, WAR, JL, JIG, JR-I, JY, RKD; Project administration – RKD, MBH, DW, CNH, JIG; Software – AS, DEH, NM, SO; Supervision – MBH, DW, RKD, CNH, JIG, KVK, JIG, RJS, JR-I, JY; Visualization – MW, AS, JL, WAR, YQ, KMC, SO, WAR, RDP, SJS, CNH; Writing – MBH, RKD, CNH, JIG.

## Competing interests

RJS is a co-founder of REquest Genomics, LLC, a company that provides epigenomic services. All other authors declare no competing interests.

## Data and materials availability

Links to all data are provided in supplementary materials.

## List of Supplementary Materials

Materials and Methods

Figs. S1 to S27

Tables S1 to S9

Dataset S1

References

## SUPPLEMENTARY DATA

### Materials and Methods

#### Data Availability

Browsers: The NAM assemblies and gene models can be accessed through their genome assembly pages https://maizegdb.org/NAM_project, which provide the genome browser metadata and links to downloads for each assembly.

Downloads: Downloads can be accessed directly from the MaizeGDB download site (https://maizegdb.org/download). NAM gene models can be downloaded, viewed on the genome browsers, and searched via the gene center (https://maizegdb.org/gene_center/gene). BLAST targets for the NAM assemblies and their gene models are available for the MaizeGDB BLAST tool (https://blast.maizegdb.org).

Raw sequence data: Raw data used for all the assemblies including the PacBio Sequel reads, Illumina short reads, BioNano optical maps are available through ENA BioProject IDs PRJEB31061 and PRJEB32225. RNA-Seq reads for various tissues can be found through ENA ArrayExpress IDs E-MTAB-8633 and E-MTAB-8628 and EM-Seq reads are available through ENA ArrayExpress under ID E-MTAB-10028.

Other data: Other files, tables and supplemental data can be found in CyVerse /iplant/home/shared/NAM/NAM_genome_and_annotation_Jan2021_release. Links to the NLR trees can be found at https://itol.embl.de/shared/xCJbI9ndshEK.

Scripts: Scripts used to generate and analyze data are at https://github.com/HuffordLab/NAM-genomes.

#### USDA funding statement

This research was supported in part by the US. Department of Agriculture, Agricultural Research Service. Mention of trade names or commercial products in this publication is solely for the purpose of providing specific information and does not imply recommendation or endorsement by the U.S. Department of Agriculture. USDA is an equal opportunity provider and Employer.

#### Plant Material

Inbred NAM lines were obtained from GRIN Global (**Table S1**) and tissue collected as previously reported (*60*). Briefly, original accessions were selfed for one generation at Curtiss Farm at Iowa State University. Using single-seed-descent ears derived from this propagation, 144 seedlings were greenhouse grown to V2 vegetative growth stage at Iowa State University. After 48hr etiolation, 30 grams young leaf tissue was harvested, flash frozen, and submitted for CTAB, or nuclei-based high molecular weight DNA isolation for downstream analysis. Remaining seed from our single-seed-descent ears has been deposited and is publicly available through GRIN Global (**Table S1**).

#### DNA preparation for sequencing

High molecular weight DNA was isolated using either a standard CTAB protocol or a modified version in which nuclei were first isolated, thereby removing the plastid and mitochondrial genomes (**Table S1**). The CTAB procedure was a slightly modified version of the original method (*61*). Nuclei isolations were based on the method of Luo and Wing (*62*), with collected and washed nuclei then being resuspended in CTAB buffer and isolations completed following (*61*).

#### PacBio Sequencing

Sequencing libraries were constructed following PacBio’s template prep protocols for the Express Template Prep Kit 2.0. For all lines except Ki11, NC350, and B73, samples were sequenced using Sequel binding and sequencing chemistry v2.1. Ki11, NC350, and B73 were sequenced using Sequel binding and sequencing chemistry v3.0.

#### Illumina Sequencing

The same DNA used for PacBio sequencing was used for Illumina sequencing. PCR-free DNA sequencing libraries were prepared using the Kapa HyperPrep PCR-free kit (#KK8505). The sequencing libraries were checked for quality on an Agilent Fragment Analyzer and the final concentrations estimated using qPCR. PE150 libraries were sequenced on the Illumina NextSeq 500 using the 300 cycles high output kit.

#### Optical Map Generation

DNA was extracted for optical map construction using the Bionano Prep™ Plant Tissue DNA Isolation Kit and a slightly modified protocol. For each inbred, approximately 0.5 g of etiolated leaf tissue was harvested from young seedlings germinated under soil-free conditions and grown in the dark for approximately 2 weeks. Leaves were treated with a 2% formaldehyde fixing solution, washed, cut into small pieces, and homogenized with a Qiagen Tissueruptor probe. Free nuclei were then concentrated through centrifugation at 2000 x g, washed, isolated by gradient centrifugation, and embedded in a low-melting-point agarose plug. The plug was treated with proteinase K and RNase A and washed four times in Bionano Wash Buffer and five times in TE buffer. Finally, purified, ultra-high-molecular-weight nuclear DNA (uHMW nDNA) was recovered by melting the plug, digesting with agarase and subjecting the sample to drop dialysis against TE.

Labeling was performed using the DLS Kit (Bionano Genomics Cat.80005) following manufacturer’s recommendations with slight modification. In total, 1 ug uHMW nDNA was incubated along with DLE-1 Enzyme, DL-Green and DLE-1 Buffer for 2:20 h at 37 °C, followed by 20 min at 70 °C. Subsequently, a second proteinase K digestion and cleanup of unincorporated DL-Green label was performed, and labeled DNA was combined with Flow Buffer, DTT, and incubated overnight at 4 °C. DNA was stained and quantified by adding Bionano DNA Stain to a final concentration of 1 microliter per 0.1 microgram of DNA. The labeled sample was then loaded onto a Bionano chip flow cell where molecules were separated, imaged, and digitized in the Saphyr System according to the manufacturer’s recommendations (https://bionanogenomics.com/support-page/saphyr-system/). Data visualization, processing, and DLS map assembly were conducted using the Bionano Genomics software Access, Solve and Tools.

#### Genome Assembly and Hybrid Scaffolding

Raw illumina reads were first used to verify homozygosity of inbreds by comparing percent heterozygosity of SNPs using BWA-MEM (*63*) and GATK (*64*) to publicly available HapMap2 maize SNP data (*43*). Loci that were monomorphic across lines were removed for this analysis. The data were also subsampled to 10,000 to 50,000 SNPs in order to generate a phylogenetic tree using SNPhylo (*65*) for the purpose of verifying line identity.

PacBio subreads were error-corrected with Falcon (*66*) using the longest 50x coverage and an average read correction rate set to 75% (-e 0.75) with local alignments at a minimum of 3000 bp (-l 3000). The usage of -l 3000 instead of the default -l 2500 performs better for highly repetitive genome species such as maize. We required a minimum of two reads and a maximum of 200 reads for error corrections (--min_cov 2 --max_n_read 200). For sequence assembly, the exact matching k-mers between two reads was set to 24 bp (-k 24) with a read correction rate of 95% (-e 0.95) and local alignments of at least 1000 bp (-l 1000). Corrected reads ranged from 32x-56x coverage and were characterized by N50s ranging from 16.2 – 23.2 kbp. These reads were trimmed and assembled with Canu (v1.8) (*67*) with the following modification of default parameters: correctedErrorRate=0.065 corMhapSensitivity=normal ovlMerThreshold=500 utgOvlMerThreshold=150. This version of Canu fixes a bug in previous versions that generated truncations in large contigs during the consensus stage. The resulting contigs were filtered to a minimum contig length of 30 kbp.

Sequence polishing of contigs was conducted using both PacBio and Illumina data sets. First, raw PacBio reads were aligned to contigs using the software pbmm2 (a PacBio wrapper for minimap2 (*68*)). The PacBio consensus algorithm tool Arrow was then run under default parameters (https://github.com/PacificBiosciences/pbbioconda). PacBio polished contigs were then polished with either PE 150 bp Illumina reads (the majority of samples) or 10X Chromium linked reads (CML52 and Il14H). The PE Illumina reads ranged from 26x-73x depth and were aligned to contigs using minimap2. Subsequently, the assembly tool Pilon v1.22 (https://github.com/broadinstitute/pilon) was used to correct individual base errors and small indels under the following modifications to default parameters: --fix bases --minmq 30 -- mindepth 10. Chromium linked reads were aligned to contigs using Longranger v2.2.2 (https://support.10xgenomics.com/genome-exome/software/downloads/latest<u>?</u>) with Pilon run as described above.

The PacBio sequence assembly was merged with the optical map using the hybrid scaffolding module of BionanoSolve (v3.4.0) and Bionano Access (v1.3.0). Default parameters from optArguments_nonhaplotype_noES_DLE1_saphyr.xml were used. At this stage three forms of gaps were generated: 1) N gaps of various sizes (not 100Ns or 13Ns). These are rough estimates of missing sequence where the Bionano map was contiguous but there were no PacBio contigs that matched. Sizes are calibrated by the Bionano software and are generally accurate within 500 bp. 2) 100N gaps. These represent gaps of unknown size between scaffolds. They generally occur in centromeres and knobs. 3) 13N gaps. These are assembly artifacts associated with repetitive regions. They occur when two contigs are aligned to the same optical map and they overlap on the ends, indicating that they are independently assembled parts of a single contiguous region (however due to the repetitiveness or residual heterozygosity, were not assembled together at the sequence level). Bionano software does not remove this overlap and instead joins the contigs end-to-end and marks the join by 13Ns. This creates a software-induced sequence duplication of several hundred bp to several kb. For the B73 assembly **only** (version 5.0) the contig overlaps marked by 13Ns were hand curated and removed.

We emphasize that any segment of a genome containing a 13N gap, when aligned to any other genome, will show apparent structural variation that does not reflect a biological difference, but instead reflects an assembly artifact associated with contig overlap. These can be identified by scanning the sequence for 13N gaps.

#### Pseudomolecule Construction

Pseudomolecules were constructed from the hybrid scaffolds using ALLMAPS (v0.8.12; (*69*) as described in our previous assembly of the B73-Ab10 line (*42*). Briefly, we used pan-genome anchor (*4*) and Golden Gate (*11*) markers for all NAM lines and the IBM (Intermated B73 x Mo17) genetic map (*70*) in the case of B73 for pseudomolecule construction. Pan-genome anchor markers were downloaded from the CyVerse Data Commons (*71*) and processed to obtain coordinates 50 bp upstream and downstream of the marker position, and sequences from the B73 V3 assembly were then extracted. These sequences were mapped to an indexed NAM assembly using HiSat2 (v2.1.0) (*72, 73*) with fine-tuning to map short sequences reliably. By disabling splicing (--no-spliced-alignment), forcing global alignment (-- end-to-end), and including high read, reference gap open, and extension penalties (--rdg 10000,10000 and --rfg 10000,10000), full-length mapping of marker sequence was achieved. Only reads with mapping quality higher than 30 and tag XM:0 (unique mapping) were retained as the final set of mapped marker sequences. These markers were then combined with the metadata to generate a pan-genome marker input file for ALLMAPS (predicted distance information with their mapped position) in CSV format. For preparing the IBM and the Golden Gate genetic maps, the marker information was downloaded from MaizeGDB (IBM: https://www.maizegdb.org/complete_map?id=887740; GoldenGate: https://www.maizegdb.org/data_center/map?id=1160762) and processed to yield markers in fasta format and metadata in a tsv file. Methods for mapping and processing these markers were identical to pan-genome anchor markers.

ALLMAPS was run using CSV files as inputs (pangenome.csv and goldengate.csv) and configured to use scaffolds with more than 20 uniquely mapped markers (--mincount=20). Gap inserts between the scaffolds was set to 100 (--gapsize-100). Pseudomolecules were finalized after inspecting the marker placement plot and the scaffold directions. Any small scaffolds nested within the large scaffolds were identified as heterozygous and were excluded from the final pseudomolecule. These scaffolds were named with the prefix “alt-scaf” and were saved as unplaced scaffolds. Synteny dotplots were generated using the scaffolds as well as pseudomolecule assemblies against the B73 genome by following the ISUgenomics Bioinformatics Workbook (https://bioinformaticsworkbook.org/dataWrangling/genome-dotplots.html). Dot plots helped confirm the placement and orientation of scaffolds. Briefly, the repeats were masked using RepeatMasker (v4.0.9) (*74*) and the Maize TE Consortium (MTEC) curated library (https://github.com/oushujun/MTEC) (*75*). RepeatMasker was configured to use the NCBI engine (rmblastn) (*76*) with a quick search option (-q) and GFF as a preferred output. The repeat-masked genomes were then aligned using minimap2 (*68*) (v2.2) and set to break at 5% divergence (-x asm5). The paf files were filtered to eliminate alignments less than 1 kbp and dotplots were generated using the R package dotPlotly (https://github.com/tpoorten/dotPlotly). The AGP construction method along with the scripts are detailed in the “agp-generation” section of the companion GitHub site.

#### Genome Quality Assessment

To assess the contiguity and gene space completeness of the NAM genome assemblies, different quality metrics (**Table S2**) were calculated using the GenomeQC tool (*77*). Embryophyta odb9 dataset (n = 1,440) and Augustus species ‘maize’ were provided as the input parameters to calculate the BUSCO metrics.

The LTR Assembly Index (LAI) (*18*) was used to assess the contiguity of TE assembly. First, intact LTR retrotransposon (LTR-RT) candidates of each genome (pseudomolecules only) were identified using LTRharvest (v1.6.1) (*78*) and LTR_FINDER_parallel (v1.1) (*79*), then filtered by LTR_retriever (v2.9.0) (*80*) with default parameters. The LAI program (beta3.2) was used to calculate LAI values of each genome based on a total LTR content of 76.34%, an LTR identity of 94.854% (-totLTR 76.34 -iden 94.854), and the intact LTR-RTs identified from the genome. The LAI was comparable among NAM lines with an average of 28 (SD = 0.23), which is considered “gold” quality (*18*). The percentage of structurally annotated TEs was lower than previously reported (*21*) due to more effective filtering of false positives (*80*) and the fact that only intact TEs were structurally annotated in this study.

#### RNA-seq

Total RNA was extracted using the Qiagen RNeasy plant mini kit from ten tissues. These were (1) primary root and (2) coleoptile at six days after planting, (3) base of the 10^th^ leaf, (4) middle of the 10^th^ leaf, (5) tip of the 10^th^ leaf at the Vegetative 11 (V11) growth stage, (6) meiotic tassel and (7) immature ear at the V18 growth stage, (8) anthers at the Reproductive 1 (R1) growth stage, (9) endosperm and (10) embryo at 16 days after pollination. With a few exceptions, for each tissue in each NAM founder, mRNA was sequenced from two biological replicates that were composed of mRNA from three individual plants. In the case of endosperm and embryo, 50 kernels per plant were used (for a total of 150 per biological replicate). For tissues 1-5, plants were grown in University of Minnesota greenhouses in Metro-Mix300 (Sun Gro Horticulture) at 27°C/24°C day/night and 16h/8h light/dark. For tissues 6-10, plants were grown outdoors at the Minnesota Agricultural Experiment Station in Saint Paul, MN with 30-inch row spacing at ∼52,000 plants per hectare.

For each sample, total RNA was assayed by Bioanalyzer to determine the quantity and integrity of the sample. Concentrations were normalized in 25uL of nuclease-free water and sequencing libraries prepared using KAPA’s Stranded mRNA-seq kit (#KK4821). The mRNA was enriched using oligo-dT beads, fragmented, and converted to double stranded cDNA using random hexamer priming and amplification. Libraries were pooled and sequenced on NextSeq 500 instruments using the PE75 protocol.

#### Gene Model Annotation

The 26 NAM genomes were annotated using a hybrid evidence and *ab initio* based gene prediction pipeline (*81*). Evidence-based predictions were directly inferred from the assembled transcripts, which were generated using five different genome-guided transcript assembly programs, Trinity (v2.6.6) (*76, 82*), StringTie (v1.3.4a) (*83*), Strawberry (v1.1.1) (*84*), Cufflinks (v2.2.1) (*83, 85*) and Class2 (*83, 85, 86*)) and processed using Mikado (v1.2.4) (*87*) to pick the optimal set of transcripts for each locus. To generate assembled transcripts, quality inspected RNA-seq reads from each library were mapped to their respective NAM genomes using STAR (v2.5.3a) (*88*) with an iterative 2-pass mapping approach in which splice junctions generated from the first round were used to refine alignments in the subsequent round. STAR was configured to output SAM format (with options --outSAMattributes All, --outSAMmapqUnique 10, --outFilterMismatchNmax 0) to ensure downstream analysis compatibility. Mapped reads from each library were merged, sorted, and indexed using SAMTools (v1.9)(*89*) to generate input for transcript assembly programs. All programs were run with default options with the exception of the minimum transcript length setting (when allowed), which was set to 100 bp (Trinity using -- min_contig_length 100, StringTie using -m 100 and Strawberry using -t 100) and enabling of RNAseq strandedness (Trinity using -SS_lib_type FR, Cufflinks using --library-type fr-firststrand), when available. Maximum intron size was also set to 10000 (-- genome_guided_max_intron 10000) in Trinity. While most of the assembly programs generated a GFF3 as the final output, Trinity provided fasta format transcripts. These transcripts were mapped back to the gmap (v2019-05-12) indexed genome to generate a GFF3 file (by setting the output format option -f to gff3_match_cdna).

In order to pick the final transcripts, Mikado uses assembled transcripts combined with high-confidence splice junctions generated by Portcullis (v1.1.2) (*90*) with the mapped reads as input (merged and sorted), predicted ORFs for the assembled transcripts generated by TransDecoder (v5.5.0) (*91*), and homology results of transcripts to SwissProt (viridiplantae) sequences generated by NCBI-BLAST (blastx) (v2.9.0) (*76*). While default options were used for Portcullis and TransDecoder, for blastx, maximum target sequences were set to 5 (-max_target_seqs 5) and output format to xml (-outfmt 5). The following were provided as inputs for Mikado: all transcript assemblies (with strandedness marked as True for all except for Trinity, and with equal weights) in GFF3 format, portcullis generated splice sites in bed format, TransDecoder results in bed format, homology results in XML format, and a scoring matrix in yaml. Final transcripts from Mikado were exported in GFF3 format, and transcripts and proteins were then converted to fasta format using the gffread utility of the Cufflinks package.

*Ab initio* predictions were performed using BRAKER (v2.1.2) (*92*) with both evidence-based predicted proteins and mapped RNA-seq reads as input. BRAKER was run iteratively, with the first round using the hard-masked genome (primarily to speed-up the protein alignments and to generate a hints file from the BAM file) and the second round using a soft-masked genome with proteins/RNA-seq hints for finalizing the *ab initio* predictions. Default options were used in BRAKER, with the exception that gth was substituted as the protein aligner (--prg=gth), models trained using protein alignments (--gth2traingenes), the soft-masked genome was provided as input (--softmasking), and output predictions were generated in GFF3 format (--gff3).

A working set (WS) of models was generated for each NAM line to capture the complete gene space by combining evidence based and non-overlapping BRAKER gene models using BEDtools (v2.17.0) (Aaron. A et al 2010). Additional structural improvements on the WS models were completed using the software PASA (v2.3.3) (*93*) iteratively with default options. 69,163 B73 full-length cDNA (*94*) and an additional 46,311 transcripts from 11 developmental tissues (*95*) were filtered for intron retention and then used in combination with ∼2 million maize ESTs from genbank with the Mikado generated transcripts as evidence to update WS gene models with PASA. PASA was run with default options, with a first step of aligning transcript evidence to the masked NAM genomes using GMAP (v.2018-07-04) (*96*) and Blat (v.36) (*97*). The full-length cDNA and Iso-seq transcript IDs (*98*) were passed in a text file (-f FL.acc.list) during the PASA alignment step. Valid, near-perfect alignments with 95% identity were clustered based on genome mapping location and assembled into gene structures that included the maximal number of compatible transcript alignments. PASA assemblies were then compared with NAM-generated transcript models using default parameters. PASA on average updated 12,927 protein coding models across the NAM lines (Supplementary Table3) with the majority of updates being UTR modifications (73.8%), followed by alternative isoforms (35.1%) and novel genes (5.5%). Transposable element (TE) related genes were filtered from the evidence and non-overlapping BRAKER sets using the TEsorter tool (*99*), which uses the REXdb (viridiplantae_v3.0 + metazoa_v3) database of TEs. The TE filtered WS had 110,498 gene models on average across the NAM lines (lowest of 101,754 in B73 and highest of 118,596 in Tzi8).

The TE filtered WS models were given Annotation Edit Distance (AED) scores using MAKER-P (v.3.0) (Campbell. M et al, 2014). Only models with AED < 0.75 passed to the high-confidence set (HCS). The number of gene models dropped to an average of 45,768 transcripts per NAM accession in the HCS (lowest of 44,424 in B73 and highest of 47,262 in Mo18w) (Supplementary Table4). The HCS gene models were further classified based on homology to related species, and assigned coding and non-coding biotypes. Protein sequences were aligned to the canonical translations of gene models from *Sorghum bicolor*, *Oryza sativa*, *Brachypodium distachyon*, and *Arabidopsis thaliana* obtained from Gramene release 62 (*100*) using USEARCH v11.0.667_i86linux32 (*101*). The HCS gene models were checked for missing start and stop codons. On average 8,078 out of 32,470 conserved genes and 5,003 out of 8,862 lineage-specific genes had incomplete CDS. The CDS boundaries of the transcripts were modified based on conserved start codon positions or extended to a start or stop codon whenever possible. All conserved genes in addition to lineage-specific genes that had a complete CDS were marked as protein-coding. The remaining lineage-specific genes were marked as non-coding. HCS gene models were checked and potentially split or merged using the GFF3toolkit (2.0.1) (*102*). Gene ID assignment was made as per MaizeGDB nomenclature schema (https://www.maizegdb.org/nomenclature) for each line. Functional domain identification was completed with InterProScan (v5.38-76.0) (*103*). TRaCE (*104*) was used to assign canonical transcripts based on domain coverage, protein length, and similarity to transcripts assembled by Stringtie. Finally, the gene annotations were imported to ensembl core databases, verified, and validated for translation using the ensembl API (*105*). The exported GFF3 annotation files were validated and reformatted again using GFF3toolkit.

#### Centromere annotation

Functional centromere regions were annotated using ChIP-seq with antisera to maize Centromeric Histone H3 (CENH3) as described (*40*). CENH3 ChIP-seq data from B97, CML228, CML322, CML247, CML52, CML69, Ky21, Mo18W, M37W, M162W, Ms71, NC358, Oh43, and Tx303 are from (*39*) and can be obtained from GenBank (SRP067358); and ChIP-seq reads for B73, CML103, CML277, CML333, HP301, Il14H, Ki11, Ki3, NC350, Oh7B, P39 and Tzi8 are from (*40*) and available under project PRJNA639705.

Centromere positions of each NAM line were projected to B73 by mapping both CENH3 ChIP-seq data and genomic input data to the B73 genome with bwa-mem (v0.7.17) (*63*). ChIP enrichment was calculated by normalizing RPKM values from the ChIP data against the genomic input in 5 kbp windows with deeptools (v3.3) (*106*). Enriched islands with a ratio above 2.5 were identified and merged with a distance interval of 1 Mbp using bedtools (v2.29) (*107*). The final centromere coordinates were determined by visual inspection of the ChIP-seq peaks in IGV (v2.8) (*108*). Centromeres that were not mappable by CENH3 ChIP were defined as the midpoint of the largest CentC array in B73.

#### DNA methylation and identification of unmethylated regions (UMRs)

DNA methylome sequencing libraries were prepared from the second leaves of 5 to 9 plants (at a stage before the unfurling of the first leaves) using the NEBNext® Enzymatic Methyl-seq Kit (New England Biolabs #E7120S). At least two biological replicates were prepared and analyzed in this way for B73 each NAM founder. The input for each sample consisted of 200 ng of genomic DNA that had been combined with 1 pg of control pUC19 DNA and 20 pg of control lambda DNA and sonicated to fragments averaging ∼700 bp in length using a Diagenode Bioruptor. All libraries were amplified with 4 or 5 PCR cycles. The libraries were Illumina sequenced using paired-end 150 nt reads, with a minimum of 300 million reads per NAM founder, divided between biological replicates. Reads were trimmed of adapter sequence using cutadapt (version 2.6, default parameters except -q 20 -a AGATCGGAAGAGC -A AGATCGGAAGAGC -O) (*109*). Reads were aligned to each genome and methylation values called using BS-Seeker2 (version 2.1.5, default parameters except -m 1 --aligner=bowtie2 -X 1000) (*110*). The previously separate replicates were merged together for subsequent analyses. Methylation averages were calculated for whole genomes and for specific sets of genetic elements using CGmapTools (*111*). UMRs were identified as described in (*112*). Briefly, reference genomes were segmented into 100-bp intervals. Intervals lacking at least four covered CHG-context cytosines (CHGs) were discarded. Coverage was calculated on a per-cytosine basis and summed over each interval, and any interval with less than 20 reads covering CHGs was discarded. Intervals with methylation of greater than 20%, calculated using the weighted methylation formula (*113*), were also discarded. This was repeated on 20bp sliding increments, and all overlapping intervals or intervals separated by only 20 bp were merged to define larger UMRs. UMR edges were then trimmed such that their boundaries were defined by CHGs with less than or equal to 20% methylation. At this stage UMRs that overlapped with blacklisted regions (identified based on abnormally high coverage of the 150nt paired end Illumina reads that were used in each genome’s assembly) were discarded. This process was repeated using CG/CHG methylation combined rather than CHG methylation alone and both sets of UMRs were merged. Finally, UMRs that were shorter than 150 bp in length were discarded.

A conservative approach was used to identify UMRs present within B73 that were either methylated or unmethylated in other NAM lines at homologous loci. B73 UMRs were divided into quarters of equal length. Based on EM-seq reads mapped to B73, a minimum CHG coverage of ten and a minimum covered CHG count of four was enforced in each UMR quarter. UMRs that satisfied these criteria were separated into those with >= 50% mCHG in all quarters (methylated) or < 20% mCHG in all quarters (unmethylated). UMRs in which all four quartiles were methylated were classified as high-confidence differentially methylated regions (DMRs), and UMRs in which all four quartiles were unmethylated were classified as high-confidence conserved UMRs. For each B73-NAM pair, the DMRs and conserved UMRs were compared to corresponding pan-gene expression levels averaged across the ten tissues and replicates. TPM was used for normalized pan-gene expression. Pan-genes that were absent from B73 were excluded from this analysis. A subset of TSS-overlapping pan-genes were selected as those where a region from -10 to +400 bp of the TSS was at least 98% overlapped by a DMR or conserved UMR. The NAM founder TPM/B73 TPM ratio was calculated for each selected pan-gene. This analysis was performed separately on each NAM founder-B73 pan-gene pair.

#### UMR enrichment analyses

A collapsed set of UMRs identified in all NAM lines using B73 as reference was generated using the bedtools (v2.27.1) (*107*) merge function. These UMRs were then intersected with significant SNPs (p-value >0.05) from GWAS analyses using bedtools intersect. Enrichment of significant associations was calculated by shuffling UMR intervals using the bedtools shuffle function. To estimate genome-wide enrichment of significant associations in UMRs, shuffling was permitted in all regions except for sequencing gaps. To assess enrichment in low-copy, genic regions, shuffling was limited to pan-gene coordinates, plus 15-kb flanking regions (bedtools slop), allowing overlap with known UMRs. Summary statistics of intersecting SNPs were tabulated using bash scripts and GNU datamash (v1.3) (*114*). The interval size distribution, feature overlap and other metrics were computed using the GenomicRanges package (*115*).

UMRs identified in B73 were also examined to assess intersection with coding features using the GFF files. With the bedtools intersect function, the number of significant SNPs (p- value <0.05) from the GWAS that are present in the B73 UMR region and also in the genic feature were computed and tabulated.

#### ATAC-seq and identification of accessible chromatin regions (ACRs)

Three biological replicates were included in each ATAC-seq sample, from two tissues sources. The first tissue source was V1 stage, above-ground tissue, excluding most of the exposed 1st and 2nd leaf blade but including coleoptile, sheath and ligule portions of 1st and 2nd leaves, developing inner leaves, and shoot apical meristem. The second was the same 2nd leaf tissue used for EM-seq. Oh43 and Mo18w were exceptions in that they only included two biological replicates from the first tissue source and none from the second. Finely-ground, frozen tissue was suspended in 500 uL of LB01 buffer (15mM Tris pH 7.5, 2mM EDTA, 0.5mM Spermine, 80mM KCl, 20mM NaCl, 15mM 2-mercaptoethanol, 0.15% Triton X-100). The lysate was filtered through two layers of miracloth (Millipore #475855), stained with ∼1 uM DAPI and loaded onto a Beckman Coulter Moflo XDP flow cytometer instrument. A total of 20,000 nuclei were sorted from each replicate and NAM founder and combined into a single tube containing ∼350 uL of LB01. Sorted nuclei containing all NAM founders within a single tube were spun in a swinging bucket centrifuge (5 minutes, 500 rcf) and resuspended in 10 uL of LB01, visualized and counted on a hemocytometer under a fluorescent microscope, and adjusted to a final concentration of 3,200 nuclei per uL using diluted nuclei buffer (10X Genomics #1000176).

For each replicate, a total of 16,000 nuclei were loaded per well on the Next GEM Chip H (10X Genomics #1000162), targeting a final recovery of ∼10,000 single nuclei per library. Single-cell ATAC-seq libraries were prepared according to the manufacturer’s instructions (10X Genomics #1000176, Chromium Next GEM v1.1) using the Chromium Controller (10X Genomics #120223). Libraries were sequenced using 100-bp paired-end reads on an Illumina S2 flow cell (NovaSeq 6000) in dual-index mode with 8 and 16 cycles for i7 and i5, respectively. Replicated (3x) libraries were demultiplexed from single-cell ATAC-seq binary base call sequences files (BCL) output from the Illumina S2 NovaSeq 6000 with 10X Genomics *cellranger-atac mkfastq* software (v1.2) and aligned to the B73 RefGen_V4 reference genome (*21*) using *cellranger-atac count* (v1.2), resulting in three distinct sets of FASTQ files containing pooled NAM founders for each replicate. To assign genotypes to individual cells, a VCF file containing NAM founder SNP information mapped to RefGen_V4 (*116*) was used to partition reads by their respective genomes. Specifically, genotype probabilities for individual cells were estimated using *demuxlet* with non-default values (--min-total 100) (*117*). Cells with genotype probabilities less than 0.95 were removed from the analysis. Cell genotype classifications were taken as the genotype with the maximum probability. Finally, raw reads from cells corresponding to the same genotype were concatenated into forward and reverse FASTQ files.

Demultiplexing, genotyping and FASTQ concatenation were repeated for each pool of biological replicates separately. Reads were then processed with fastp (version 0.20.0) (*118*), with the parameters --detect_adapter_for_pe --correction --length_required 35. Reads were aligned to the NAM reference genomes and to the B73v5 genome with Bowtie2 (version 2.3.5.1) (*119*), with parameters --local --very-sensitive-local --seed 1 -q --no-mixed --no-discordant -- maxins 1000. Aligned SAM files were converted to BAM files with SAMtools (version 1.10) (*89*), with the parameters view -b -h -S. Duplicate reads were removed with Sambamba (version 0.7.1) (*120*), with the parameters markdup --remove-duplicates and reads were filtered for MAPQ scores of 30 or higher with sort -F “mapping_quality >= 30”. ATAC-seq peaks were called with MACS2 (version 2.2.7.1) (*121*), with the parameters callpeak --format BAMPE -- gsize 1.8e+9 --keep-dup all --qvalue 0.005.

#### Pan-genome Analysis

The pipeline described in (*122*) was used to identify homoeologous gene pairs using the canonical transcript for each gene (/iplant/home/shared/NAM/NAM_genome_and_annotation_Jan2021_release). This method requires that genes have high sequence similarity and fall within the same syntenic block. Syntenic blocks were identified by whole-genome alignment using MUMmer4 version 4.0.0.beta2 (*123*) with --mum -c 1000 option. As a result, any genes unanchored to scaffolds would have been excluded.

To compare gene content among genomes, we first created a blast database of all canonical gene model transcripts using the makeblastdb command in ncbi_blast+ version 2.8.1 with default settings. An all-by-all blast was then performed between each pair of genomes. The results were parsed to retain hits between genes within a syntenic block that had a p-value of no more than 1×10^-10^. Gene pairs from the 26 genomes were added stepwise into a matrix using the custom R script stepwise_add_to_matrix.R and executed in R version 3.6.3 (*124*). Tandem duplicate genes as defined in (*122*) were compressed into semicolon-separated values in the matrix and counted as a single pan-gene for downstream analyses. Lines in the initial pan-genome matrix that had redundant transcripts were compressed such that each transcript was contained in a single line. Additional tandem duplicates identified during this process were also merged and all tandem duplicates are presented as semicolon-separated values in the pan-gene matrix. There remain cases where two biologically separate gene models are annotated as a single combined gene model, as well as genes that are incorrectly split (i.e. one biological transcript annotated as two separate transcripts) within the final annotation that can cause genes to be incorrectly identified as tandem duplicates in the pan-gene matrix.

To recover pan-genes that exist in a genome but were not annotated, representative pan-gene sequences for all pan-genes were mapped to each NAM genome excluding scaffold sequences using GMAP version 2015-09-29 (*96*) with output one path option. Alignments were filtered to have greater than 90% coverage and 90% identity and to be in the same syntenic block to the pan-gene. GMAP canonical transcripts with CDS larger than 200 bp were intersected with annotation gff CDS files containing only the canonical transcript using intersectBed from bedtools v2.29.2 (*107*) with -f 0.90 -r option. GMAP coordinates that intersected with a canonical transcript at these thresholds were replaced by the canonical transcript name. Pan-genes that overlap with a non-canonical annotated transcript are still represented as GMAP coordinates in the matrix.

#### Transposable Element Annotation

For each genome, both structurally intact and fragmented transposable elements were annotated using the Extensive *de-novo* TE Annotator (EDTA v1.9.0) (*34*). The curated and updated Maize TE Consortium (MTEC) library (https://github.com/oushujun/MTEC) was used as the base library, so that EDTA could identify novel TE families in each genome (--curatedlib maizeTE02052020). The high-confidence, evidence-based *de-novo* gene annotation of each genome was used to remove genic sequences in the TE annotation (--cds genome.cds.fasta). The species parameter was set to Maize (--species Maize) to use the maize-specific classification model for terminal-inverted repeat (TIR) elements in the TIR-Learner pipeline (*125*) that was included in the EDTA package. To further control false annotations, novel TE families that were single-copy in the source genome were identified using RepeatMasker (v4.0.9) (*126*) and further removed. The remaining novel TE families of all NAM founder genomes were aggregated following the removal of redundant sequences using the “cleanup_nested.pl” script in the EDTA package. The non-redundant, novel TE library was aggregated with the MTEC library to form the pan-NAM founder TE library, which was used to annotate all NAM founder genomes using RepeatMasker (v4.0.9) with parameters “-q -div 40 - cutoff 225”. The homology-based annotations (by RepeatMasker) were combined with the structure-based annotation (by EDTA) and formed comprehensive TE annotations for each NAM founder genome. TEs found by structure-based annotations were classified into families using the pan-genome TE library based on the 80-80-80 rule, that is 80% of the TE sequence was covered by a library sequence with more than 80% identity and longer than 80 bp. Annotation statistics were summarized and plotted using custom Perl and R scripts.

#### Characterization of tandem repeat arrays

The coordinates of CentC, knob180, TR-1 and rDNA repeat arrays were determined by blasting consensus sequences to the assemblies as described previously (*42*). Arrays were defined as ≥100 kbp clusters composed of at least 10% repeat sequences with no more than 100 kbp spacing between repeat units. The completeness of assembled repeat arrays was evaluated by comparing the amount of repeats incorporated in the pseudomolecules with that estimated with 150bp Illumina reads from the same genomes. Assembled repeats in NAM genomes were identified with BLAST (v2.2.26) and quantified by counting non-overlapping repeat monomers. The absolute repeat abundance for each NAM line was estimated with Illumina reads. Paired-end short reads were subsampled to approximately 3X coverage and aligned as single-end sequences against consensus repeats with BLAST (v2.2.26; -b 5000 -F F). Non-overlapping fragments (≥ 30bp) mapped to repeat sequences in each read were summed as the total repeat abundance. The total repeats were then normalized by read coverage and genome sizes measured by flow cytometry (*40, 43*).

Knob arrays were categorized as lying in a mid-arm position if they were farther than 2 Mbp from either chromosome end. To identify conserved knob positions, the syntenic positions for each array were defined by the up and downstream sorghum orthologous gene from their respective genome. The knob arrays that correspond to classical knobs were identified by comparing relative coordinates based on karyotypes (*127*) to genomic coordinates of knob arrays in IGV. For the subset of knobs displayed for structural variation (Fig. 3), only arrays that were syntenic to knobs of at least 100 kbp in length in B73 were considered.

Arrays of telomeric 7-mer repeat units (5’-TTTAGGG-3) were identified using the motif search algorithm of the Tandem Repeat Finder tool (version 4.09 with parameters 2 7 7 80 10 50 500 -f -d -m -h) (*128*). To identify the boundaries of subtelomeric repeat arrays, fasta files of the maize subtelomeric sequences were first downloaded from the NCBI database with the following accession numbers: EU253568.1, S46927.1, S46926.1, S46925.1, CL569186.1, AF020266.1, AF020265.1. Subtelomeric sequences were blasted (BLAST v2.7.1+) against each chromosome of the pseudomolecule assembly for each NAM line; blast hits were then filtered for query coverage (≥80%) and percentage identify (≥80%). The coordinates of the filtered blast hits were clustered using bedtools (version 2.27.1) (*107*) to identify the start and stop coordinates of the repeat clusters. IGV was then used to manually check and refine the boundaries of telomeric and subtelomeric repeats located on the ends of the short and long arm of each chromosome for each of the NAM lines.

#### Fractionation Analysis

For fractionation analyses, the exons from the outgroup Sorghum bicolor (Sbicolor_313_v3.1 from Phytozome) were aligned to the previously described repeatmasked NAM and B73 genomes; annotated maize genes were not used. Tandem arrays for primary Sorghum CDS transcripts were filtered out with the script s.paralog_clusters.pl by selecting gene model paralogs (i.e., sharing the same gene tree) that were clustered with four or fewer non-paralogous intervening genes as determined by the file tree_id.sorghum_bicolor.txt generated by Gramene. Exons from this filtered Sorghum CDS set were extracted using the Sorghum gff file and the Sorghum genomic fasta file using bedtools getfasta (*107*) and were aligned to the repeatmasked maize genomes using BLAST (*76*), -task dc-megablast, no max target sequences (see project GitHub for scripts and detailed parameters). Sorghum and all the NAM founders plus B73v5 were also filtered for tandem arrays using Tandem Repeat Finder (*128*), parameters 2 7 7 80 10 50 2000 -l 1 -d –h. The coordinates of these filters were applied to the blast outputs and all blast hits that fell within these coordinates in either Sorghum or maize were removed using bedtools intersect with the parameter –v to select only blast hits with no tandem repeat overlap. All sorghum genes with a tandem repeat homeolog in any NAM/B73 were removed from consideration; this was found by running the same bedtools intersect command except with –wa –wb instead of –v for Sorghum hit coordinates that corresponded to any NAM/B73 tandem duplication. Only Sorghum genes that had clear and distinct homeolog associations were used; those that mapped to more than two syntenic regions were removed.

DagChainer (*129*) was run using parameters optimized for the large size and complexity of maize and its large distance between genes and between syntenic orthologs: -s -I -D 1000000 -g 40000 -A 15 (-A being much higher than the default value since exon collinearity was being determined, not whole-gene collinearity). Orthologs were scored in each NAM line based on alignment of at least one Sorghum exon to a single gene-space locus syntenic with the query Sorghum gene. Total Sorghum exon alignment counts per locus per maize genome post-DagChainer were deduced using bedtools groupBy (*107*). Fully retained orthologs were considered to be those that had all expected Sorghum exons aligned to the orthologous region in each maize genome. Partial deletions were those where fewer than the total number of exons of the Sorghum ortholog aligned. Cases where no Sorghum exons aligned at the expected orthologous region in each maize genome were scored as fully fractionated; an ortholog is considered not fully fractionated even if only one exon in one NAM line is present. Sorghum exon alignments were used instead of gene model alignments in order to capture partially deleted loci which may not be represented by a gene model annotation.

DagChainer results were then filtered for Sorghum exon alignments falling within the identified subgenome blocks of B73 version 4 associated with syntenic coordinates of Sorghum gene models from the file B73v4.subgenome_reconstruction.gff3 from Gramene (/iplant/home/yjiao/B73_RefGen_V4/Annotation). Since maize underwent a genome duplication event after diverging from Sorghum, there would be two expected sorghum orthologs in each maize line; therefore, each Sorghum exon orthologous in maize would have been expected to have two syntenic copies unless fractionation had ensued. Only blast outputs in each NAM founder that share the same B73 subgenome chromosome as the sorghum orthologs were selected, such that if an exon is retained on both subgenomes, it would have two alignments to one Sorghum exon, differentiated in part by maize chromosome ID. Most inversions within the various maize lines were contained within subgenomic blocks, so they would not be excluded by this method. However, special consideration had to be made for Oh7B’s translocation of distal chromosome 10 to chromosome 9; all alignments that fell within that translocated region were given the identity associated with the subgenome identity of distal chromosome 10 for the purposes of fractionation assignment. The fractionation pipeline was tested multiple times for accuracy using CoGe’s GEvo visualization platform (*130*) and the pipeline was changed as needed to increase true positive alignments and reduce false fractionation calls, resulting in the finalized fractionation dataset (**Supplemental Dataset 1**). Segregating fractionation loci were manually checked in CoGe, and pipeline errors (i.e. false exon deletion calls) or missing exons associated with sequencing gaps as well as loci where flanking syntenic sequence could not be confirmed or exons were too fragmented to make a confident call were removed.

GO enrichment of both homeologs for unfractionated and segregating fractionating pairs was generated in AgriGOv2 (*131*) (http://systemsbiology.cau.edu.cn/agriGOv2/) using the B73 v4 Ensembl gene model dataset corresponding to the B73 NAM gene models (associations generated by CoGe SynFind, default parameters, using the B73 NAM gene model set as query), with parameters SEA, FDR 0.05, Bonferroni correction, and a minimum of 5 mapping entries.

#### Structural Variant Detection

Structural variants (SV) were characterized using data generated from 1) long reads of each NAM mapped to B73, 2) chromosomal genome assemblies of each NAM aligned to B73, and 3) *in silico* digested assemblies (to simulate a Bionano optical map) of each NAM line aligned to the B73 map.

For the long-read-based SV characterization, error corrected reads from each NAM line were mapped to B73 using NGMLR (v0.2.7) (*132*) with the “--presets” option set to “pacbio” and with “--bam-fix” enabled. The mapping step was trivially parallelized by splitting the input files (PacBio reads) and mapping them simultaneously to the reference genome, followed by merging the output bam files to a single bam file using samtools merge (v1.9). The merged BAM file was then used with SNIFFLES (v1.0.11) (*132*) for calling structural variants in a two round process. The first round of SNIFFLES used stringent parameters (--max_num_splits 2, -- min_support 20, --min_zmw 2, --min_seq_size 5000, --max_distance 5000, --cluster, and -- cluster_support 2) with minimum SV size set to 100 (--min_length 100) and generated a VCF format output for each NAM line separately. The individual VCF files were then merged using SURVIVOR (v1.0.6) (*133*), with the max distance between breakpoints set to 1000, taking the SV type and strand into account, without using the estimating SV size option or taking the minimum size of SV into account. Since this merged SV set does not have genotype information, another round of SNIFFLES was run to force SV calls across all NAM lines. In the second round, the merged SVs were provided as input (--Ivcf) along with the BAM files (mapped reads). The final genotyped SVs were combined using SURVIVOR with the same options.

Whole genome sequence alignments of each NAM against the B73 reference were generated using minimap2 (v2.17-r941) (*68*). The PAF-formatted alignments were generated using default options along with -c, (output cigar string), -x asm5 (use of ∼0.1% sequence divergence preset) and --cs (encode bases at mismatches and INDELs) options. The generated paf file was sorted using the core utilities sort command, followed by paftools (k8 paftools.js call) (*68*) to characterize variants. The output format was then converted to a bed file in order to visualize SV in IGV (*134*) using a simple awk command.

For characterizing large SVs, each NAM genome was subjected to *in silico* digestion with the fa2cmap_multi_color.pl script from the BioNano solve program, using CTTAAG as the enzyme motif. This generates a simulated, assembled BioNano map in cmap format. The cmap files were aligned against the B73 cmap file using RefAligner tool from runCharacterize.py and runSV.py script of BioNano solve. Default options were used for both steps, with the arguments supplied through an XML file (optArguments_nonhaplotype_noES_DLE1_saphyr.xml). The resulting smap file (with the list of structural variants detected between query maps and reference maps in tsv format), was then converted to VCF format using the smap_to_vcf_v2.py script. The final SV file in VCF format was filtered to only include SVs greater than 1 Mbp using an awk command. Due to lack of resolution near the breakpoints, the SVs were subjected to manual inspection using the paftools alignment in IGV and synteny dot-plots, to refine the start and stop of the SVs called using this method. Calls of Bionano SVs across all NAM lines were made by selecting common boundaries across the lines. The most 5’ start position and the most 3’ end position were used as the coordinates for the collapsed SV, and the genotypic calls for these overlapping SVs from the same individual were merged. The final curated SVs were combined to generate a joint SV file using SURVIVOR, with similar options as explained before. The final SV set was generated by merging the SNIFFLES SVs with the curated BioNano SVs.

#### Analysis of Flowering-time Genes

As proof-of-concept that SVs affect important traits, we closely investigated 39 known flowering-time genes (*53*). We found the B73v5 coordinates for these 39 flowering-time genes and extracted the high confidence SVs of those gene coordinates (genic regions) plus 5 kb upstream (promoter regions) using bedtools (*107*). SVs for each genome relative to the B73v5 genome were further filtered to include only insertions or deletions. These data were formatted for the IGV browser (*134*). For each promoter and genic region of a flowering-time gene across all genomes, unique insertion or deletion events were catalogued manually. These candidate SVs were investigated for association with changes in gene expression using t-tests between lines with and without a unique indel.

Transcripts Per Million (TPM) was calculated for each candidate gene across six specific tissues: V11 leaf base, middle, and tip; V18 tassel; and R1 anthers and ears. The presence or absence of a candidate SV was used to predict the TPM of the candidate gene for a specific tissue (t-tests, accounting for (un)equal variances between groups). Out of a total of 62 unique indels and 372 tests while using the Benjamini-Hochberg procedure for multiple testing correction at alpha equal to 0.05 (*135*), we found 18 unique indels significant and 24 significant tests. Focus for intense study was on those significant indels that were present in at least 2 or more NAM lines and those genes which had multiple significant indels.

Additionally, we inspected the previous 39 candidates as well as an additional 134 known flowering time genes (Li et al 2016) for differences in gene expression between the temperate and tropical lines without tissue specificity, i.e. the TPM value was averaged across all tissues for a given line. Similar cut-off criteria were used as before. While no candidates surpassed the multiple testing cut-off, there were candidates with greater than +/- 2 log2 fold change between temperate and tropical lines. Candidates that met the log2 fold change cutoff were manually scanned for indels using the IGV browser as before (*134*). If an indel was found segregating between lines, an ANOVA determined if there were significant differences between indel haplotypes (Figure S17). Further confirmation was achieved using CoGe (*130*) to manually inspect these loci. TE annotations gave support to a TE origin for these candidate SVs.

Using a permutation test, the Li et al 2016 candidates were significantly enriched with GWAS SNPs for Days to Silking, Anthesis Silking Interval, and Days to Anthesis (exact p-value ranged from 0 - 0.028). Neither the Li et al 2016 candidates or the Dong et al 2012 candidates were more variable (i.e. had greater coefficient of variation in expression) as random subsets of genes (p-value 0.677-0.995).

#### Glossy 15 analysis

Two insertions were identified as candidates, 337 bp and 881 bp in size, associated with changes in gene expression changes (short insertion: t = -3.932, p = 6.354×10^-04^, long insertion: t = 3.151, p = 2.923×10^-03^). The shorter insertion passed a log2 fold change cut off of (+/-) 2 at 2.5 while the longer one did not at -1.78. Those lines that contained only the shorter insertion had significantly higher expression in V11 middle (F = 24.51, p = 4.39×10^-10^) and tip of the leaf (F = 24.51, p = 4.24×10^-10^) tissue than any of the other haplotypes. Lines solely containing the shorter insertion were Oh43, Il14H, P39, M37W, and CML277. This insertion was confirmed with a local alignment in COGE where Oh43, Il14H, M37W, and CML277 all showed alignment with the P39 assembly while lines without the insertion were missing alignment.

#### ZCN10 analysis

*ZCN10* had higher expression levels in tropical NAM lines compared to temperate NAM lines (est. difference = 8.49 TPM, t = -2.346, raw p = 0.0358, log2fc = 2.940). There is a single large insertion in CML247 and NC350, but this could not be verified by manual inspection with CoGe. The local alignment of CoGe did detect many deletions relative to NC350 in the upstream region of *ZCN10* in temperate lines with the exception of B73, Il14H, and Oh7b. Deletions were detected in the tropical lines CML333, CML52, and Ki11. Those with these deletions appear to have less expression than those without, but it is difficult to parse if these deletions are correlated with TPM and if so, which deletions are the most strongly correlated.

#### Dof21 analysis

*Dof21* had higher expression in tropical NAM lines compared to temperate (est. difference = 32.123 TPM, t = -2.542, raw p = 0.01898, log2fc = 1.540). P39, B73, and Il14H were temperate outliers with higher expression while CML52, NC350, NC358, and CML247 were tropical outliers with lower expression. There were 2 insertions and 1 deletion with segregating haplotypes in the promoter window. Those lines with only one of the insertions had significantly lower expression than lines with both, neither, or the deletion (F = 8.658, p = 0.000317). Lines with only one of the insertions included most of the temperate lines (except P39) and the tropical outliers. These insertions were confirmed by manual inspection with CoGe.

#### ZmCCT10 analysis

*ZmCCT10* had higher expression in tropical NAM lines than in temperate lines (est. difference = 0.2459, t = -1.844, raw p = 0.0895, log2fc = 2.063). CML247 was an outlier for high expression. There was an insertion and deletion segregating between the NAM lines, but there were no significant differences in TPM between the different haplotypes (F = 1.252, p = 0.307). These deletions likely correspond to the *CACTA* insertion found in B73 (*52*). Because of the cyclical expression pattern of *ZmCCT10*, it is likely our method of calculating TPM across tissue with a single time sample limits our ability to connect these deletions to flowering time.

#### Analysis of Disease Resistance Genes

The NLRs were extracted from the genomic DNA sequences using NLR-Annotator (*136*) and from proteomes using hmmalign with reference HMM of the grass NB-ARC (*49*). Additionally, NLRs and NLR-IDs were characterized in the Brachypodium (*137*) and maize annotations using the plant_rgenes pipeline (https://github.com/krasileva-group/plant_rgenes) (*138*) (e-value cutoff 1×10^-03^). The number of NB-ARC containing proteins was compared to those previously identified in Arabidopsis (*139*) and plotted using R package ggplot2 (*140*). The NB-ARC domain alignment was manually curated for the presence of NB-ARC domain functional motifs including Walker A, WALKER-B, RNBS-C, GLPL and RNBS-D. The NLR phylogeny was determined using RAxML MPI (v8.2.9,-f a, -x 12345, -p 12345, -# 100, -m PROTCATJTT) (*141*). The phylogeny was visualised and re-rooted on the longest internal branch in iTOL (*142*).

#### Population Genetic Analysis

*GERP*. Soft masked copies of 13 angiosperm genomes were aligned to the unmasked B73v5 reference genome using LAST (*143–148*). Repetitive elements in B73v5 were then masked in the aligned sequences. A tree with neutral evolutionary rates was estimated from four-fold degenerate sites in the alignment using rphast with default parameters (*149*). We then used the tools gerpcol and gerpelem from GERP++ (*150*) to estimate conservation scores at aligned base pairs and identify conserved elements. For gerpcol we excluded the B73 genome from the alignment to avoid reference bias.

*Enrichment analysis.* To test whether structural variants were depleted in conserved elements, we measured the overlap between structural variants and conserved GERP elements and performed Fisher’s exact tests. For tests involving combined deletions and insertions, we measured the overlap of base pairs in conserved elements with the presence of a structural variant in any of the NAM parental lines. We also tested for the depletion of deletions and insertions in conserved coding sequence, conserved noncoding sequence, and conserved non-genic sequence. In all three of these cases, the Fisher’s exact test was testing depletion compared with non-conserved elements. For tests involving insertions, we measured the overlap of GERP elements with insertion start sites. As insertions may simply move conserved elements while maintaining their function, we speculated that insertion start sites may be more meaningful than base pairs of overlap with conserved elements. Insertions were also subdivided into quartiles based on size to test whether the size of insertions was associated with its depletion in GERP elements.

To test the relationship between genomic features and the presence of SVs, we used quasi-Poisson regression in 10kb windows to explain the number of overlapping SVs based on overlap with GERP elements, accessible chromatin (*5*), recombination rate (*151*), and the number of masked base pairs in B73 (see supplemental Transposable Element Annotation).

The model takes the following form:

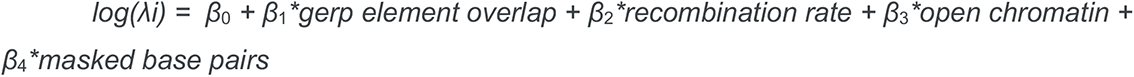

Where λ_i_ is the number of occurrences of SVs within the i^th^ window. As this is a quasi-Poisson model, the expected value of λ_i_, λ, is equal to the expected number of SVs in a window, and θλ is equal to the variance of the number of SVs in a window, where θ is a dispersion parameter.

*Simulations*. We used SLiM (*152*) to generate simulations of a 20-Mbp region consisting of two genomic element types that represented coding and non-coding sequence. The size and number of the element types were based on the approximate median values of B73v5 genome annotations described in the main text. The simulated 20-Mbp region consisted of 300 genes, each separated by 30 Kbp of non-coding bases. Each gene consisted of four 200 bp exons, and three 300 bp introns. Three types of mutations were simulated to represent neutral, 0-fold non-synonymous, and structural variants. 4-fold and 0-fold mutation types were restricted to the simulated exonic regions, where structural variants were allowed to occur anywhere along the 20 Mbp segment. The total mutation rate in exonic regions was modeled as the sum of the rate for single nucleotide mutations (*μ*) and structural variants (*μ_sv_*). Each of the three types of exonic mutations occurred in proportion to the average number of 4-fold, 0-fold, and total number of exonic bases, which were 200 kbp, 57 kbp, and 240 kbp, respectively. The distribution of deleterious fitness effects for both non-neutral mutation types were modeled using a gamma distribution with parameters for the mean (*s*_0_ and *s_sv_*) and shape (*sha*pe_0_ and *sha*pe*_sv_*), and a dominance coefficient of 0.5.

We reduced the computation time by simulating 1000 individuals in the ancestral population. We maintained the population scaled mutation rate (*θ* = 4*N_e_μ*) estimated from median pairwise diversity (*π*) in maize populations as ≈ 0.008 (*153*) by increasing the mutation rate from previous estimates of 3 × 10^-8^ (*154*) to 2 × 10^-6^. Following recommendations in the SLiM manual (*155*), we rescaled recombination rate to match the change in mutation rate using *r_scaled_*_=_(1/2) ∗ (1 − (1 − 2 ∗ *r*)*^n^*), where *r* is the original recombination rate and *n* is the rescaling factor determined by the ratio of the increased and original mutation rates. Previous estimates of median recombination in maize are 1.6 × 10^%&^(*151*); following the equation above, our simulations used a constant recombination rate of 1.05 × 10^-6^.

In addition to modeling the distribution of fitness effects, our simulations incorporated a simple demographic scenario based on previous studies of maize domestication (*156, 157*). We assume a single panmictic ancestral population of constant size (*N_a_*) that underwent an instantaneous bottleneck during domestication (*N_b_*), which we assume occurred *B_T_* = 0.067*N*_a_ generations ago based on archaeological and genetic data (*156, 158*). After the domestication bottleneck, we assume the population size grew exponentially to its present size *N*_0_, where the growth rate was derived from the change in population size as *log*(*N*_0_/*N_b_*)/*B_T_*.

*Parameter Inference with ABC.* We used Approximate Bayesian Computation (ABC) implemented in the R package *abc* (*159*) to jointly infer the distribution of fitness effects (DFE) and demographic parameters of our model. We used the folded site frequency spectra of variant sites from each mutation category generated from our simulations as input summary statistics to predict the joint posterior distribution of our model parameters. We normalized frequencies by their sum within each window and simulation. We accepted 0.5% of simulation draws with the smallest distance between simulated and observed mutation frequency bins. The posterior distribution was then inferred from the accepted draws using a neural network architecture with two hidden layers using the “neuralnet” method from the *abc* package in R. We conducted a total of 90,492 independent simulations by drawing parameters values from minimally informative prior distributions reported in the table below. Our Snakemake (*160*) pipeline and SLiM code to reproduce the simulations are available here: https://github.com/HuffordLab/NAM-genomes/tree/master/abc.

#### ABC model parameters and prior distributions

*U is short for Uniform. The prior distribution for s*_0_ *and s_s_v is a mixture, where 90% of draws are from a uniform and the remaining 10% were fixed with a selection coefficient of zero*.

| Parameter | Prior | Description |
| --- | --- | --- |
| $\mu$ | $2 \times 10^{-6}$ | Neutral mutation rate per base pair. |
| $\mu_{sv}$ | $U(10^{-10}, 10^{-7})$ | Structural variant mutation rate per base pair. |
| $r$ | $1.05 \times 10^{-6}$ | Recombination rate (scaled to match mutation rate). |
| $N_a$ | $1 \times 10^3$ | Ancestral effective population size. |
| $N_b$ | *discrete* $U(0.01N_a, N_a)$ | Instantaneous bottleneck effective population size. |
| $N_0$ | *discrete log* $U(N_a, 20N_a)$ | Modern effective population size. |
| $B_T$ | $0.067N_a$ | Bottleneck time (generations before present). |
| $s_0$ | $0.9 U(-0.1, 0) + 0.1 (s = 0)$ | Mean selection coefficient of 0-fold non-synonymous mutations for the 0-fold Gamma DFE. |
| $shape_0$ | $U(0, 100)$ | Shape parameter for 0-fold Gamma DFE. |
| $s_{sv}$ | $0.9 U(-0.1, 0) + 0.1 (s = 0)$ | Mean selection coefficient of structural variant mutations for the 0-fold Gamma DFE. |
| $shape_{sv}$ | $U(0, 100)$ | Shape parameter for structural variant Gamma DFE. |

*Model validation.* We validated our approach by testing the accuracy of 100 randomly selected simulation runs. In each case, we held out the results of one simulation and predicted its parameters using the remaining simulated data. We evaluated the accuracy and reliability of the model across all 100 runs by calculating: 1. proportion of posterior draws greater than true value (prop_gt), 2. proportion times true values fell within the 95% credible interval (w.in_cred), 3. proportion of times the mean posterior values fell within the prior (w.in_prior), and 4. The natural log of the ratio of standard deviations of the prior and and posterior distributions (log(var_sc)).

*Analysis of empirical data*. To fit our model to empirical data, we constructed 103 20-Mbp windows along the B73v5 genome. We excluded the remainder of bases at the end of each chromosome, which varied from approximately 1 Mbp to 18 Mbp. We developed a script to categorize sites as 0-fold and 4-fold Using the B73v5 reference genome and gff annotation file (https://github.com/silastittes/cds_fold). We also developed a script to calculate the folded allele frequency spectrum of each of the three mutations types in each window (https://github.com/HuffordLab/NAM-genomes/blob/master/abc/predict/src/get_nam_sfs.py). We followed the same ABC approach that was used in our model validation methods above to infer the DFE and demography parameters from the empirical data, fitting each of the 103 windows independently. To summarize across 20-Mbp windows, we used the average value of each parameter from each of the 103 posteriors.

To assess the degree of similarity between SFS data generated by the model and the empirical data, we ran simulations using 20 random draws from the posterior distributions of each genomic window. Before sampling, we excluded posterior draws that fell outside of parameter domains, and rounded demographic parameters to the nearest whole integer. From these 20 draws per window, we calculated the proportion of mutation counts in each frequency bin of the simulated SFS that were greater than observed counts, where 50% of the simulated draws should be greater than the observed under an adequate model of the data.

#### Mean and standard deviation of average posterior predictions across the 103 genomic windows

*Population size and mutation rate estimates are reported on the original scale, 100 times that of the simulated values*.

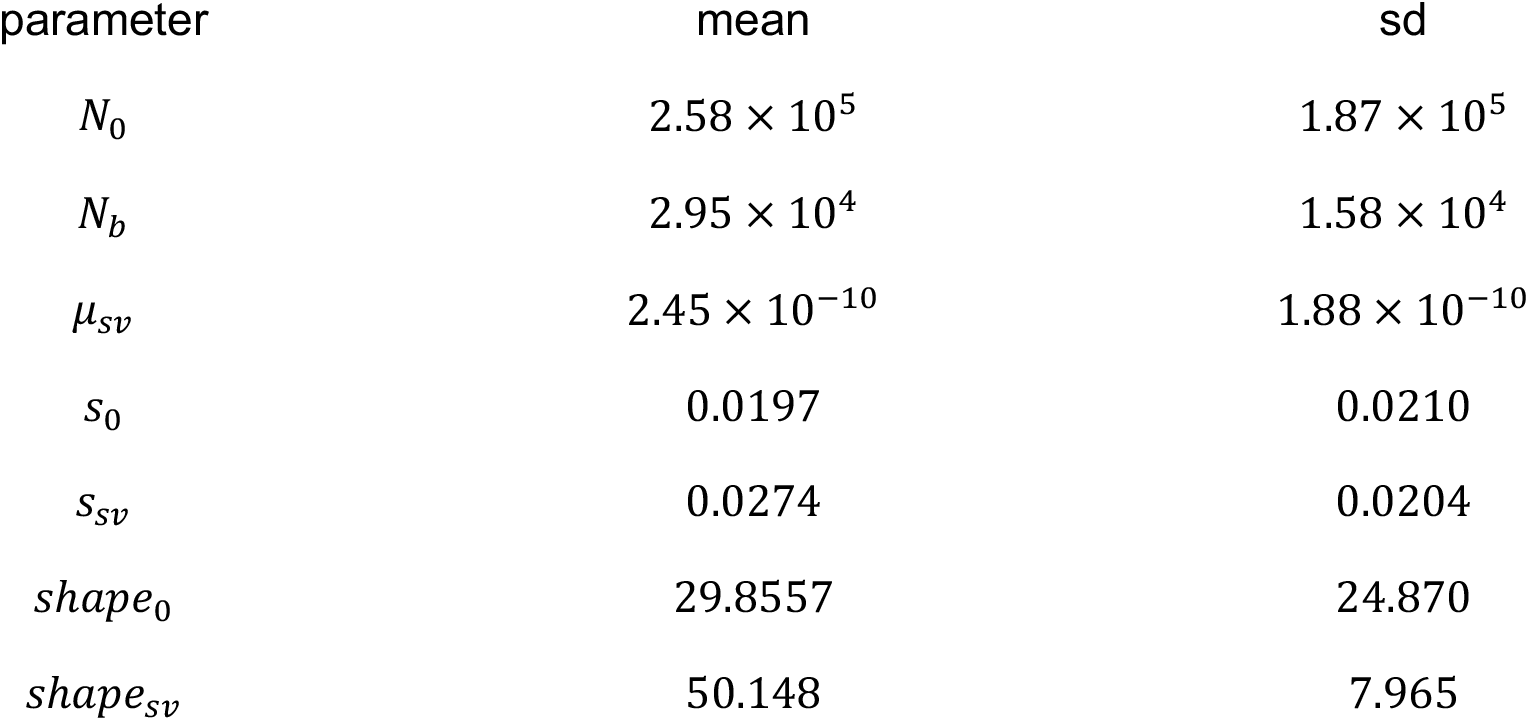

#### Genome-wide Association Study and Variance Component Analysis

We collected NAM phenotype datasets from eight publications (*13, 161–167*). Seven of the datasets are available at https://www.panzea.org/data. The phenotypic data include 36 traits, covering agronomic, developmental, domestication-related, and metabolic characteristics. Traits had already been processed by fitting a model of best linear unbiased predictions (BLUPs) on the multi-environment trial for each trait within each study. A total of 4,027 NAM RILs were used for genome-wide association study and variance component analysis. Genotype projection from NAM parents onto RILs was carried out as follows:

1. *Parental SV and Marker Identification*. Markers were identified in the parental genotypes using the PacBio and Illumina sequence data described above. During the merge step of the SNIFFLES SV calling pipeline some SVs with non-perfect overlapping boundaries were not merged. If the genotypic calls for overlapping SVs were the same across all of the parents that had genotypic calls, the genotypic information was subsequently collapsed. The boundaries were retained for the SV with the least amount of missing data or the largest one (if they had the same amount of missing data). If there was a disagreement between genotypic calls across all parents, both SVs were retained.
2. *Dataset for SV and SNP projections*. All SNIFFLES SV markers were reduced to a binary state (SV is the reference state (A) or SV is the alternate state (T)) and converted to hapmap format for projection to the RIL progeny using the middle position of the SV as the variant point position. The identified SNPs were filtered on a per family (RIL population) basis and all families were combined after per-family projections were completed. The per family filters included 1) remove parental SNPs within the boundaries of deletions using vcftools v0.1.17 (*168*) and 2) remove monomorphic SNPs.
3. *GATK SNP calling for NAM founders.* Short reads (PE150 libraries sequenced on the Illumina NextSeq 500 for polishing NAM genomes) were used for calling SNPs by mapping to the B73 genome as reference. The Genome Analysis Toolkit (GATK v4.1.3.0) HaplotypeCaller (*64, 169*), and best practices published by the Broad institute (*170*), were used along with numerous utilities in the Picard Toolkit (v2.23.3) for SNP discovery and final variant filtering (http://broadinstitute.github.io/picard/). For each read pair in fastq format, Picard was used to convert to SAM format through the FastqToSam utility. MarkIlluminaAdapters was run on SAM files to mark the Illumina adapters and generate metrics files. The SAM formatted files were converted back to interleaved fastq files using the Picard SamToFastq utility and these were mapped to the BWA-MEM-indexed B73 genome using recommended options (-M) (*171*). The obtained SAM file was converted to BAM using samtools and aligned reads were merged with unaligned reads using Picard’s MergeBamAlignment utility, marking duplicates with the MarkDuplicates utility. In the last step of processing BAM files, AddOrReplaceReadGroups was used to add the correct read-group identifier before calling variants with HaplotypeCaller. HaplotypeCaller was trivially parallelized by running simultaneously on 1-Mbp intervals of the genome (2,813 chunks, including scaffolds), and the VCF files were gathered to generate a merged, coordinate-sorted, unfiltered set of variants (SNPs and INDELS). Stringent filtering was performed on the raw set of SNPs using the expression (QD < 2.0 || FS > 60.0 || MQ < 45.0 || MQRankSum < -12.5 || ReadPosRankSum < -8.0 || DP > 5916), where DP was estimated from the DP values of the SNPs (standard deviation times 5 + mean). This filtered set of SNPs was used as “known-sites” with Picard’s BaseRecalibrator and ApplyBQSR for recalibrating the processed BAM files from the previous round. The second round of GATK HaplotypeCaller was run using the same method as before and the variants were separated (SNPs and INDELS), quality filtered, and finalized for downstream analyses.
4. *GBS SNP calling for RILs using stacks*. We followed methods, along with commands and parameters for GBS SNP calling using Stacks, from the online workbook (https://bioinformaticsworkbook.org/dataAnalysis/VariantCalling/gbs-data-snp-calling-using-stacks.html). Briefly, metadata obtained from the CyVerse Data Commons and data downloaded from NCBI-SRA (BioProject ID: SRP009896) were processed using the Stacks (v2.53) (*172*) recommended pipeline. Barcodes were formatted and used with the “process_radtags” function to demultiplex the data. The demultiplexed reads were then aligned to the B73 genome using BWA-MEM under default parameters. Output SAM files were converted to BAM, sorted, and indexed after adding the correct Read-Group for each sample with the Picard Toolkit (v2.23.3). The Stacks program command “gstacks” was run using all bam files together, followed by the “populations” command (default options except --vcf, for VCF-formatted output) to generate the final GBS SNPs file. Redundant positions were collapsed to a single line in this file.
5. *RIL Genotyping-by-Sequencing Anchor Markers*. SNPs identified from GBS data were used to define haplotype blocks for projection of our dense SV and SNP parental markers to the 4,950 NAM RILs. The GBS SNPs were filtered prior to conducting the projections. These filters and subsequent projections were applied on a per family (RIL population) basis and then all families were combined after the per-family projections were complete. The per family filters included: 1) remove SNPs that were contained within a parental deletion of 100 kbp or less (95% of all deletions) using vcftools v0.1.17 (*168*), 2) remove monomorphic SNPs, and 3) remove SNPs with greater than 70% missing data. Finally, a sliding window approach was applied to correct for possible errors during genotyping as described by (*173*). For this, a 15-bp window, with 1-bp step size, and minimum of five markers per window was used. Only SNPs with allele frequency between 0.4 and 0.6 were retained. After these filtering steps, approximately 13,000-52,000 SNPs were retained per family and used to define haplotype blocks for the parental SV and SNP projections.
6. *Parental Marker Projection to RILs*. The FILLIN plugin from TASSEL v 5.2.56 (*174*) was used to project SVs in a two-step process. First, haplotypes were created based on SNP and SV information in the parents using FILLINFindHaplotypesPlugin (-hapSize 3000 -minTaxa 1). Then, the parental haplotypes were projected onto missing genotypes in the RILs with FILLINImputationPlugin (-hapSize 3000 -hybNN false). The projections were done for each NAM family independently. Projections of the polymorphic SNPs were completed using the same methods except the haplotype size was set to a larger size (-hapSize 70000). A sliding window was again applied to the projected genotypes to correct possible errors in the projection using a 45-bp window slide, 1bp step size, and a minimum of 15 markers per window. Finally, all monomorphic SNPs were filled back into each family and all SV and SNP markers across the families were combined into a single file.
7. *Additional marker filtering for GWAS*. A genome-wide association study (GWAS) was performed by using the mixed linear model implemented in GCTA-MLMA (*175*). A total of 71,196 SVs with missing rate < 20% were included to estimate the genomic relationship matrix used for SV-based GWAS and a total of 20,470,711 SNPs with missing rate < 20% were included in the SNP-based GWAS. While the first three principal components (PCs) were calculated to correct for the population structure, we excluded the fixed terms of PCs from the GWAS models for all the traits, due to the equal to or slightly lower genetic variances compared to those in the original models. The GCTA-GREML (Genome-wide Complex Trait Analysis-REstricted Maximum Likelihood) method (*175*) was used to estimate the ratio of genetic variance to phenotypic variance. Differing from trait heritability, this method is to estimate the variance explained by genome-wide markers. We estimated three ratios from this analysis: phenotypic variance explained by all the SVs (SV-based heritability), all the SNPs (SNP-based heritability), and both SVs and SNPs (Combined-genetic heritability). The last estimation uses a method to estimate SVs-based and SNP-based heritability simultaneously in one model that was implemented with the function “mgrm”.

## Supplementary Figures

**Figure S1.**
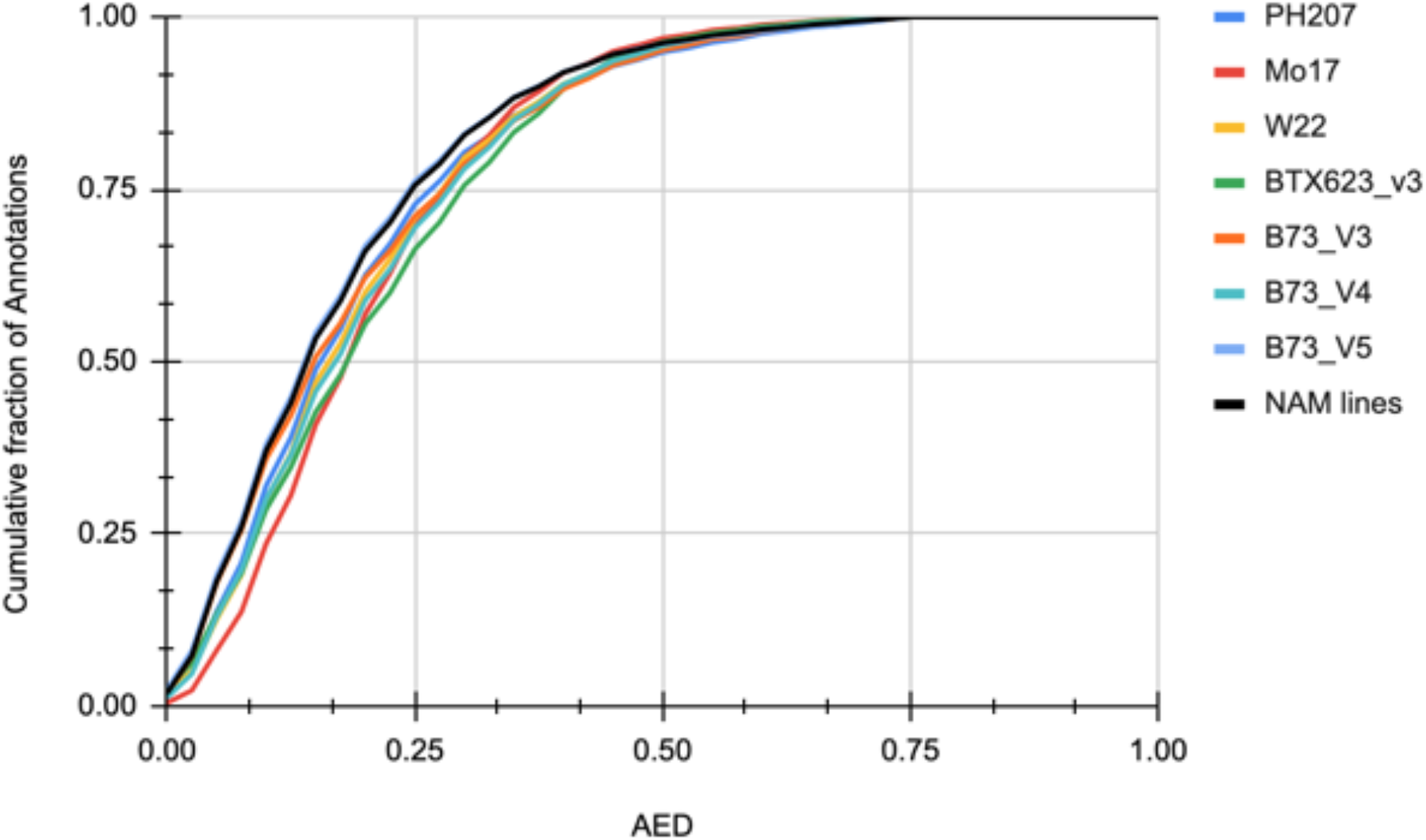
Cumulative Annotation Edit Distance (AED) scores in multiple recent genome assemblies. An AED score closer to zero indicates that more evidence supports the gene models. 83% of B73_V5 (blue) and NAM (black) gene models showed better AED values than other maize or sorghum reference annotations (*2, 6, 10, 20–22*). BTx623 is the sorghum reference genome. All others are maize assemblies.

**Figure S2.**
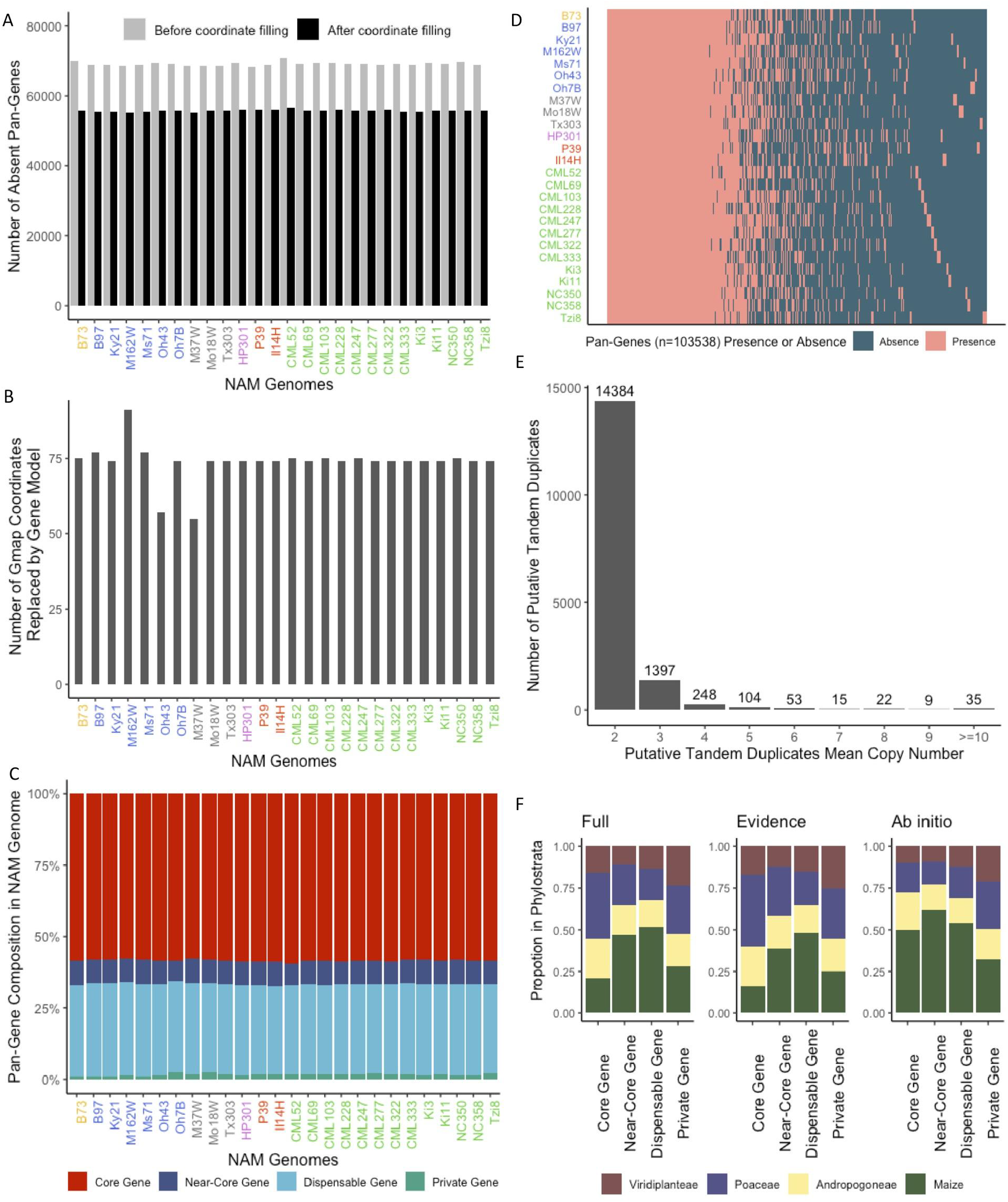
Pan-genome analysis of the gene space. **A**) Number of absent pan-genes in each genotype before and after coordinate filling. **B**) Number of GMAP coordinate fills that overlapped an annotated gene model at greater than 90% coverage. The coordinate fill was then replaced with the annotated gene model in the final pan-genome matrix. **C**) Proportion of the genes in each genome that are part of the core, near-core, dispensable, and private fractions of the pan-genome. **D**) Presence/absence (PAV) variation of each pan-gene in each genotype with pan-gene order sorted by core, near-core, dispensable, and private. In C and D, tandem duplicates were counted as a single pan-gene and coordinates were filled in when a gene was not annotated but an alignment with greater than 90% coverage and 90% identity was present within the correct homologous block. **E**) Distribution of mean copy number across genotypes that had ≥ 2 tandem copies for the 16,267 pan-genes that had a tandem duplicate in at least one genotype. Values over bars indicate the number in each copy number class. **F**) Proportion of annotated genes in each phylostrata level broken down by pan-gene frequency categories (i.e. core, near-core, dispensable, and private genes). Full is the full set of annotated gene models, Evidence is the set of gene models that were generated based on RNAseq expression evidence from 10 unique tissues, and *Ab initio* are the augmented set of *ab initio* annotated gene models.

**Figure S3:**
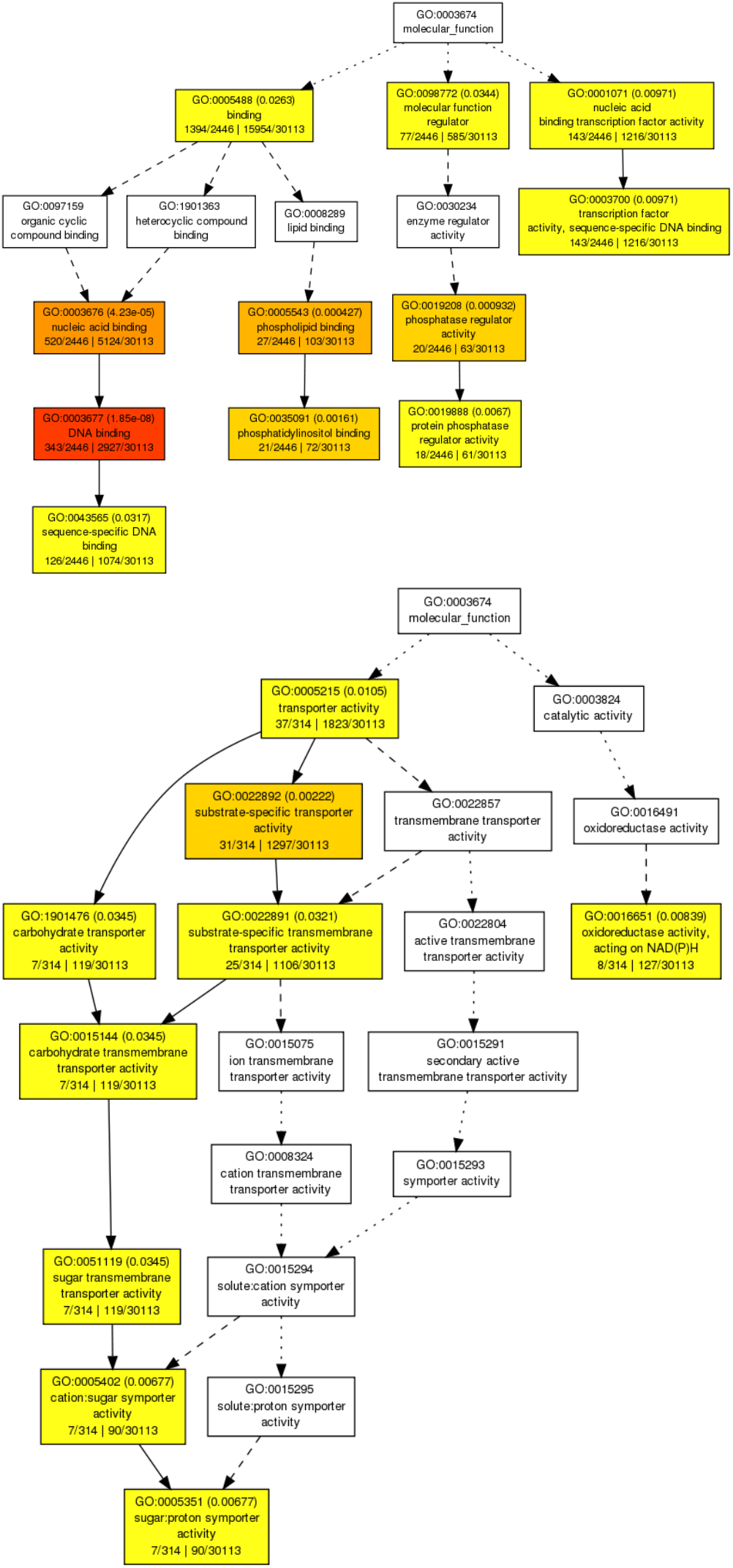
Bonferroni-corrected molecular function GO term enrichment (FDR 0.05) for loci in fully retained homeologs (top) vs loci in fractionating homeologs (bottom). Red shows strongest enrichment; yellow shows weaker (though still statistically significant) enrichment.

**Figure S4.**
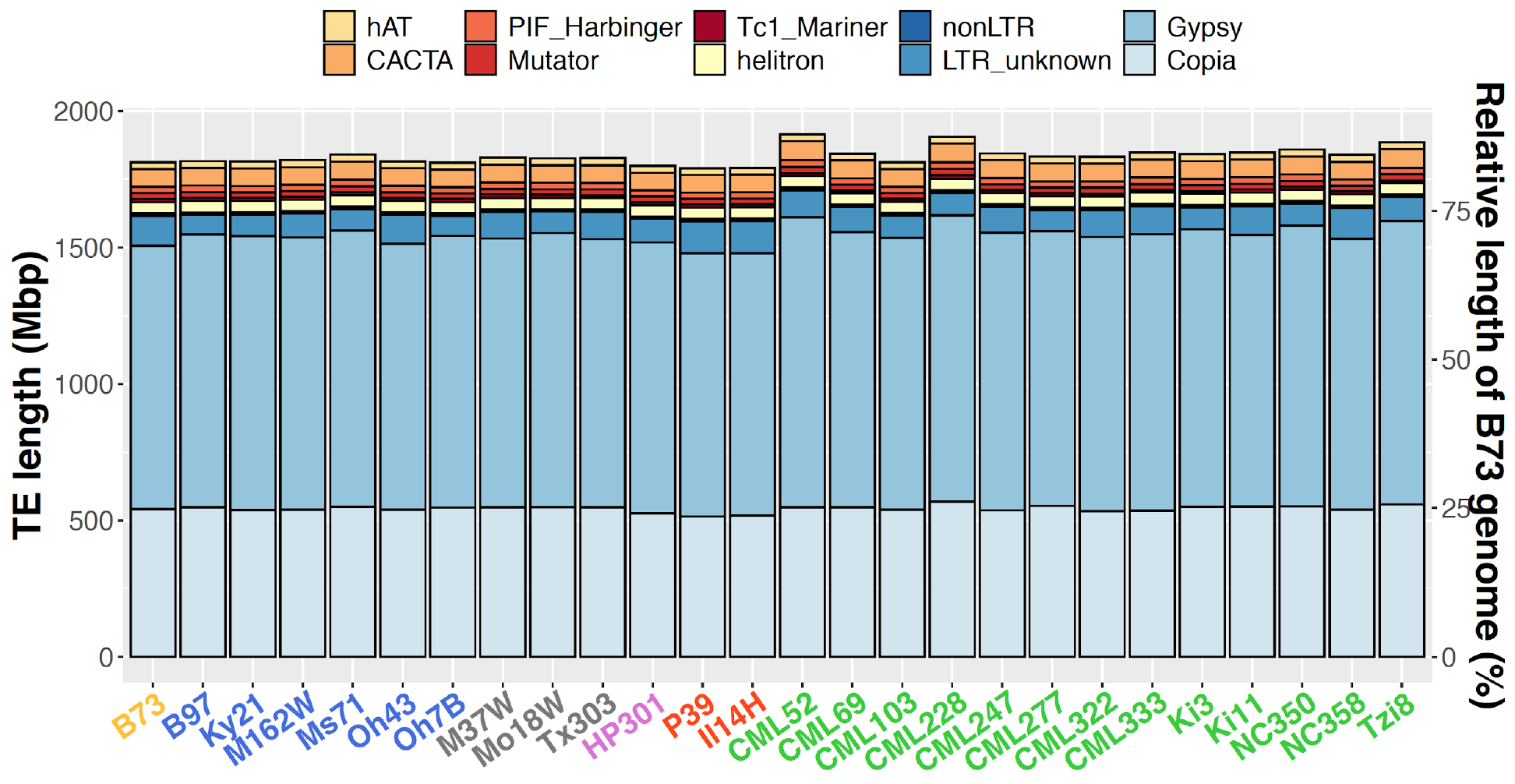
The cumulative length of repetitive sequences in NAM genomes. The right y-axis indicates the size relative to the B73v5 assembly (listed first). Genes and low-copy intergenic regions make up the rest of the assembly.

**Figure S5.**
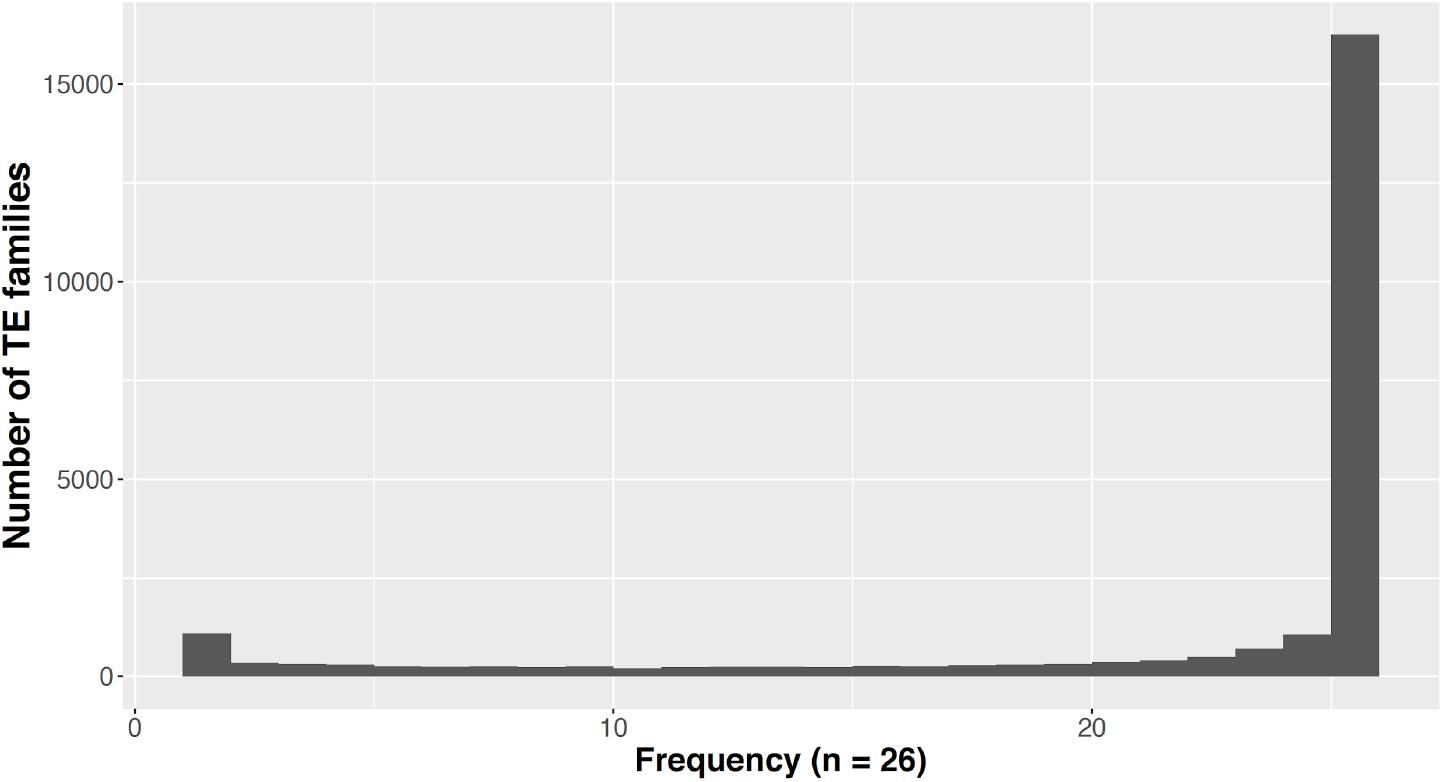
Distribution of TE families in the 26 genomes. The X axis shows the number of genomes, where 1 indicates the number of TE families found in only one genome, 2 indicates the number of TE families found in two genomes, etc.

**Figure S6.**
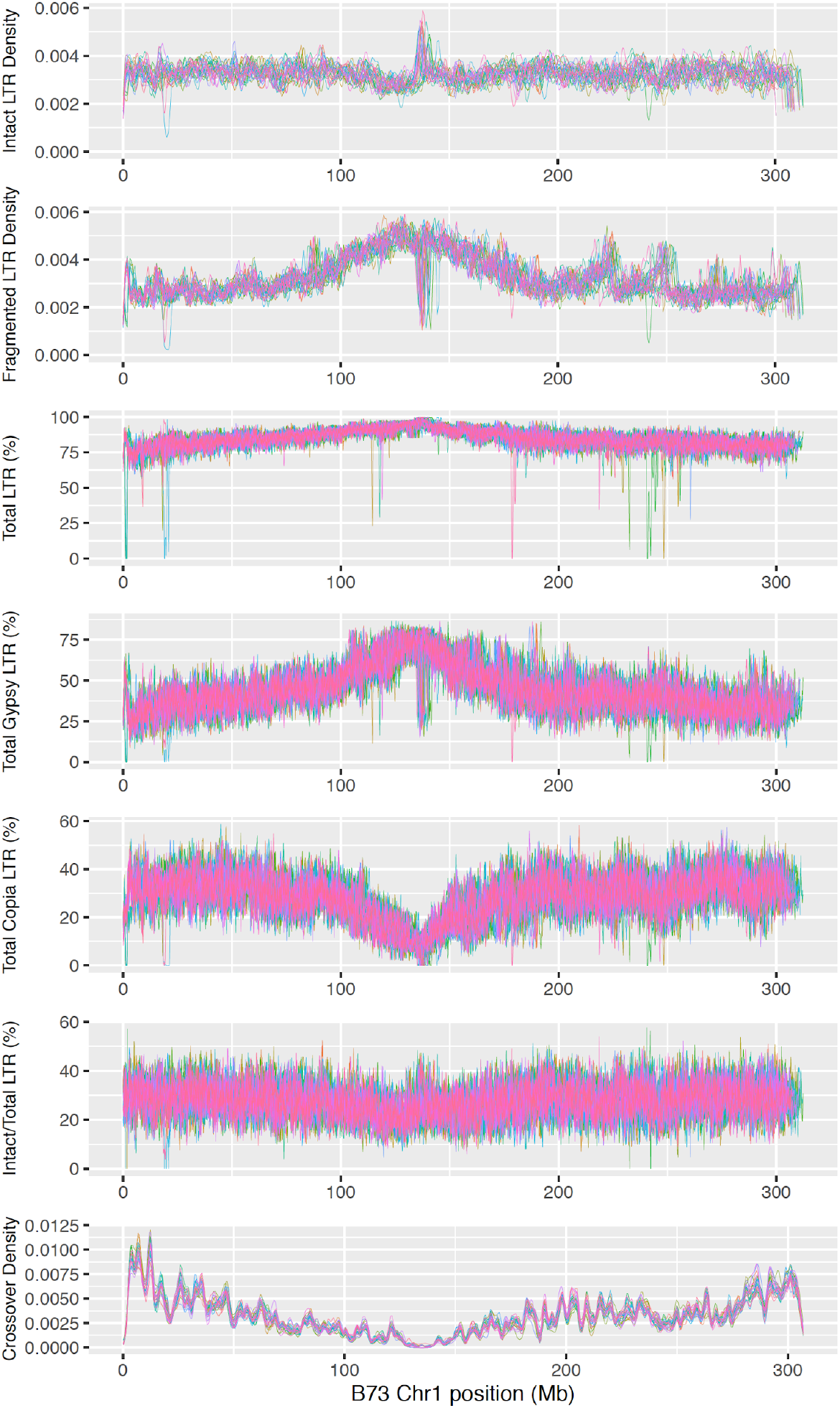
Distribution of LTR retrotransposons on chromosome 1. Each genome is represented by one color. Densities are non-parametric probability densities of the target variable (e.g., the number of intact LTR-RTs). The area under a density line sums to 1. Total LTR (including *Copia*, *Gypsy*, and unknown LTR), Total *Copia*, and Total *Gypsy* percentages are the proportion of respective LTR sequences (including both intact LTR retrotransposons and associated fragmented sequences) of the total assembled sequence length calculated in 500-kbp windows and 100-kbp steps. Intact/Total LTR percent is calculated with Intact LTR percentage (in 500-kbp windows and 100-kbp steps) divided by Total LTR percentage. The LTR makeup is very similar among lines at the Mbp scale. Crossover density (with B73) for each NAM line was calculated using data from (*13*).

**Figure S7.**
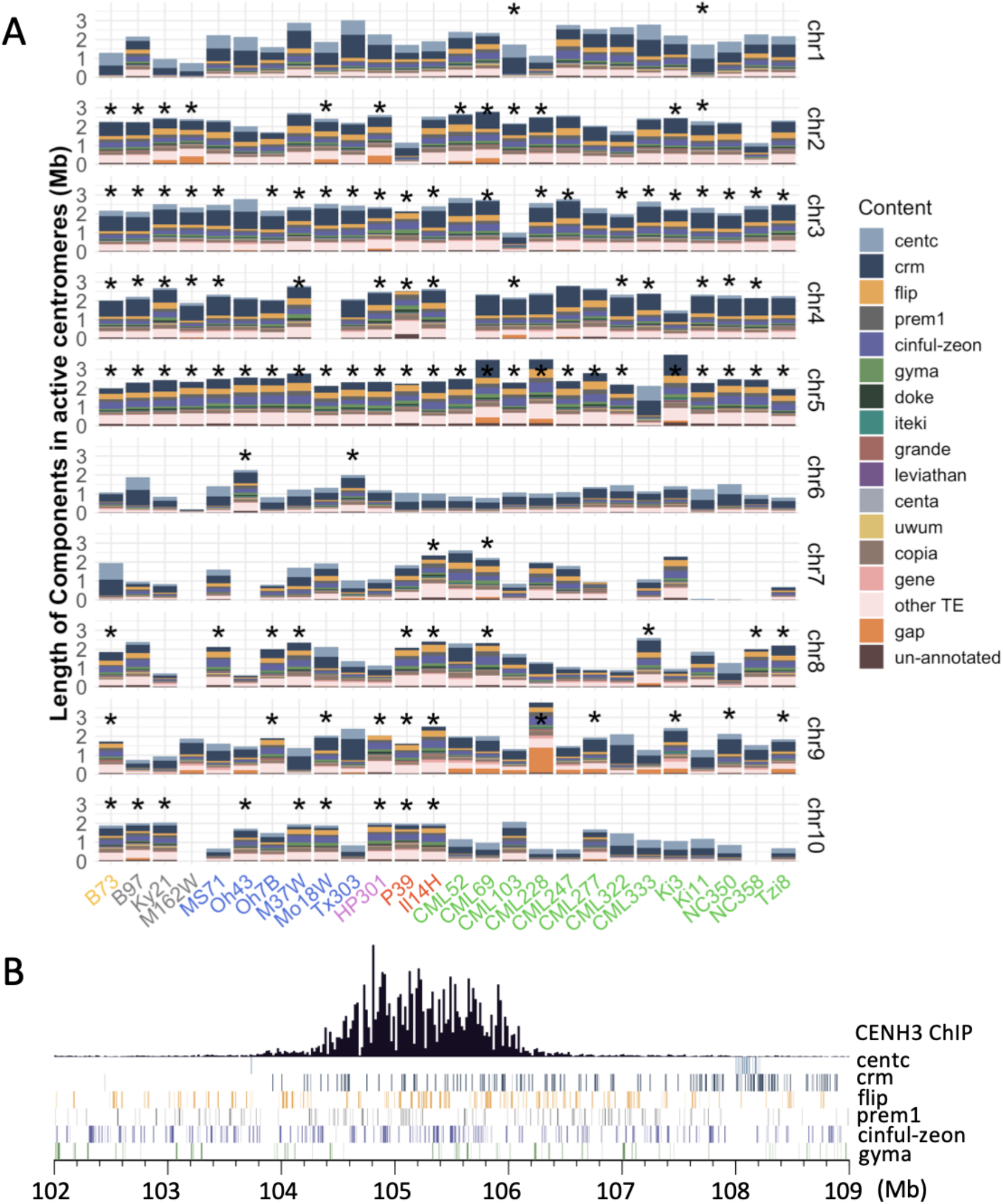
Assembly and components of functional centromeres. **A**) Distribution of transposable elements and repeats in 260 active centromeres among NAM lines. Asterisks depict fully assembled centromeres. Gap only includes gaps of known sizes. **B**) Active centromere on chromosome 5 in B73. CentC and the five most abundant transposable element families are shown as tracks in the lower panel.

**Figure S8.**
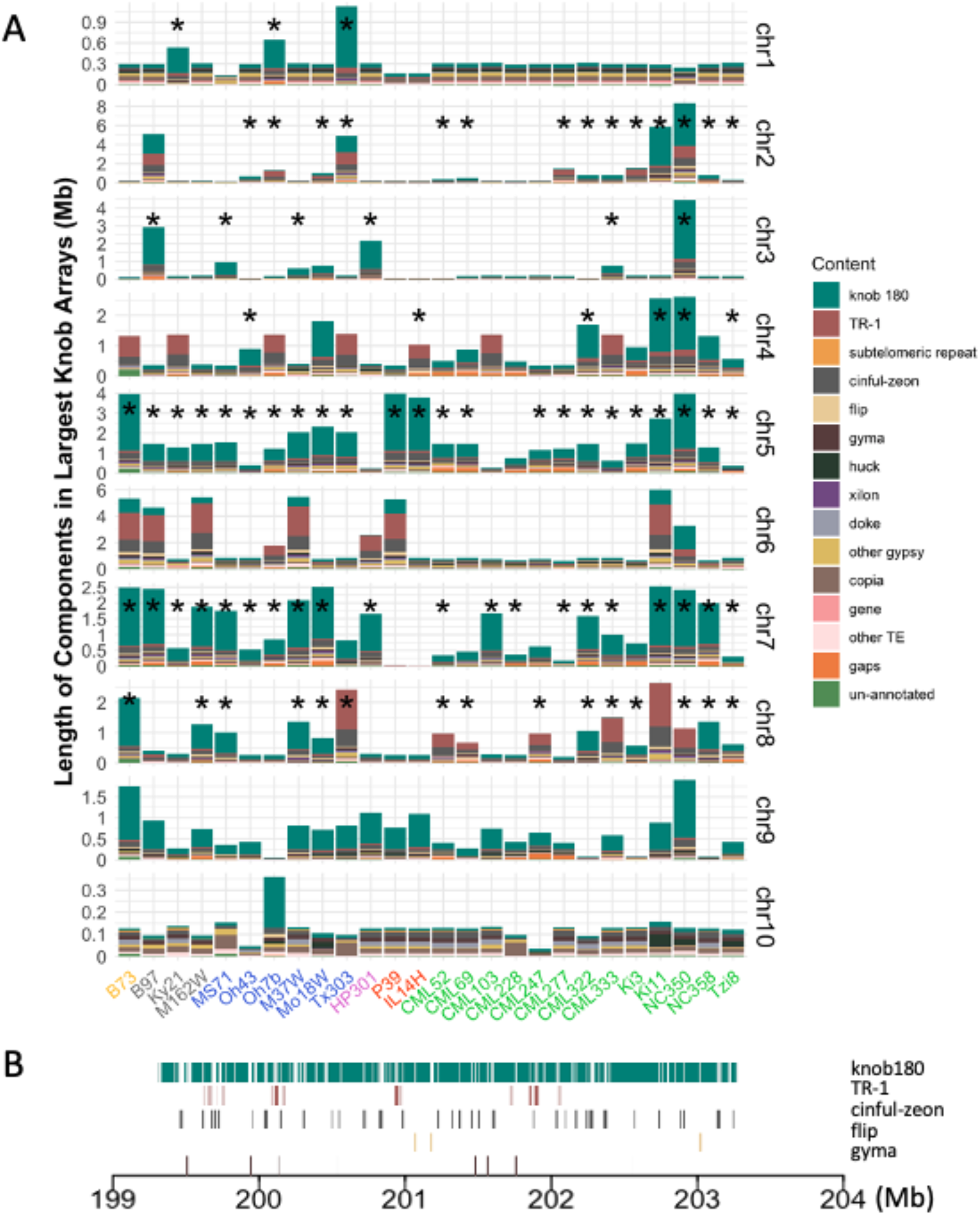
Assembly and components of repeats and transposons in the single largest knob array on each chromosome. **A**) Distribution of transposable elements and repeats in the single largest knob array on each chromosome. Lengths are based on assemblies and only include gaps of known sizes. Asterisks depict fully assembled knobs. Unknown is unannotated. **B**) Largest knob on chromosome 5 in B73. Knob repeats and the three most abundant TE families are shown as tracks.

**Figure S9.**
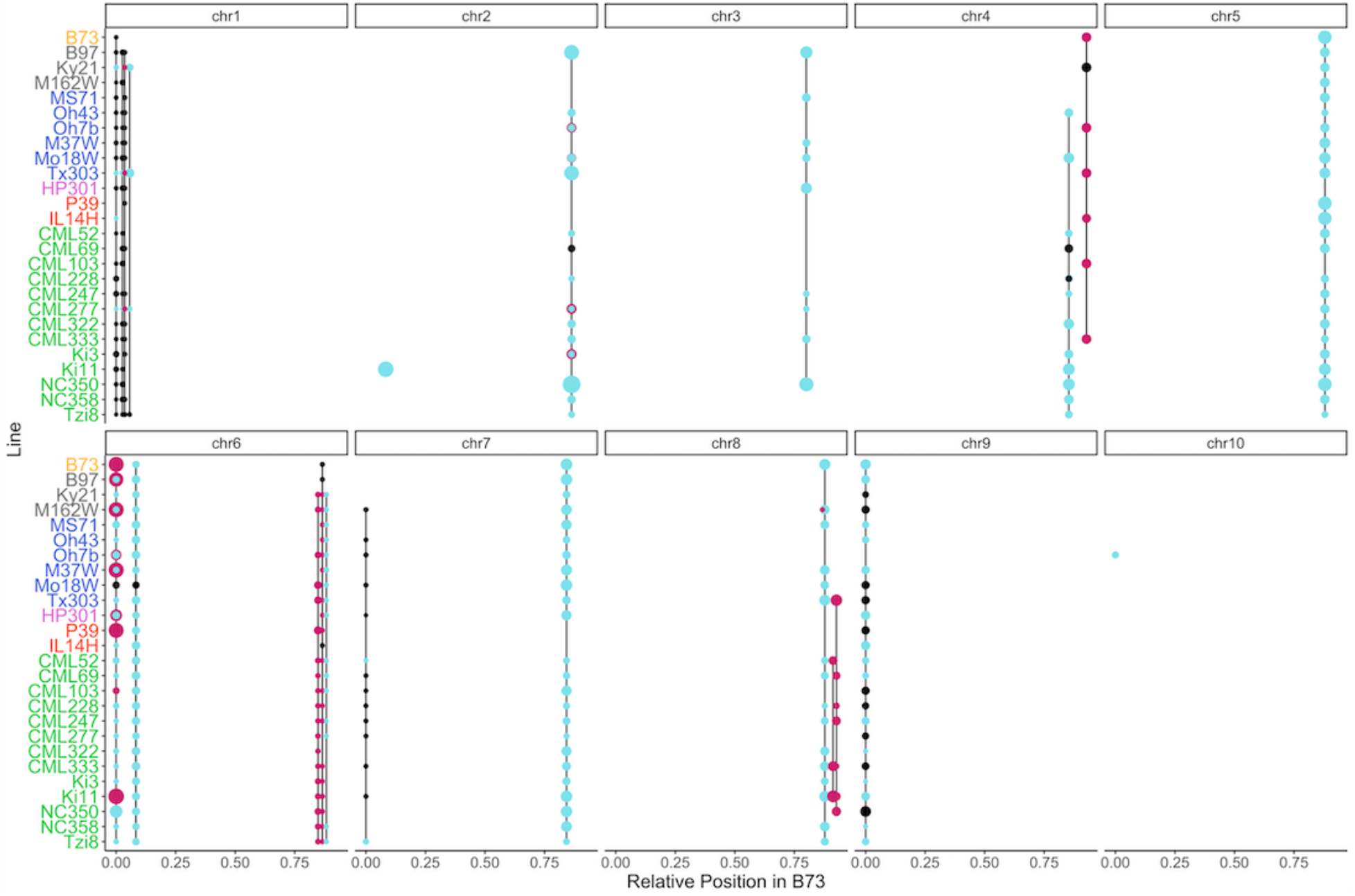
Synteny of classical knobs. Knob arrays corresponding to classical knob180 knobs (blue) and TR-1 knobs (red) are shown. Dot size corresponds to assembled array size. Syntenic arrays that are not cytologically visible are represented in black. The absence of a dot indicates there is no knob array present at the syntenic location.

**Figure S10.**
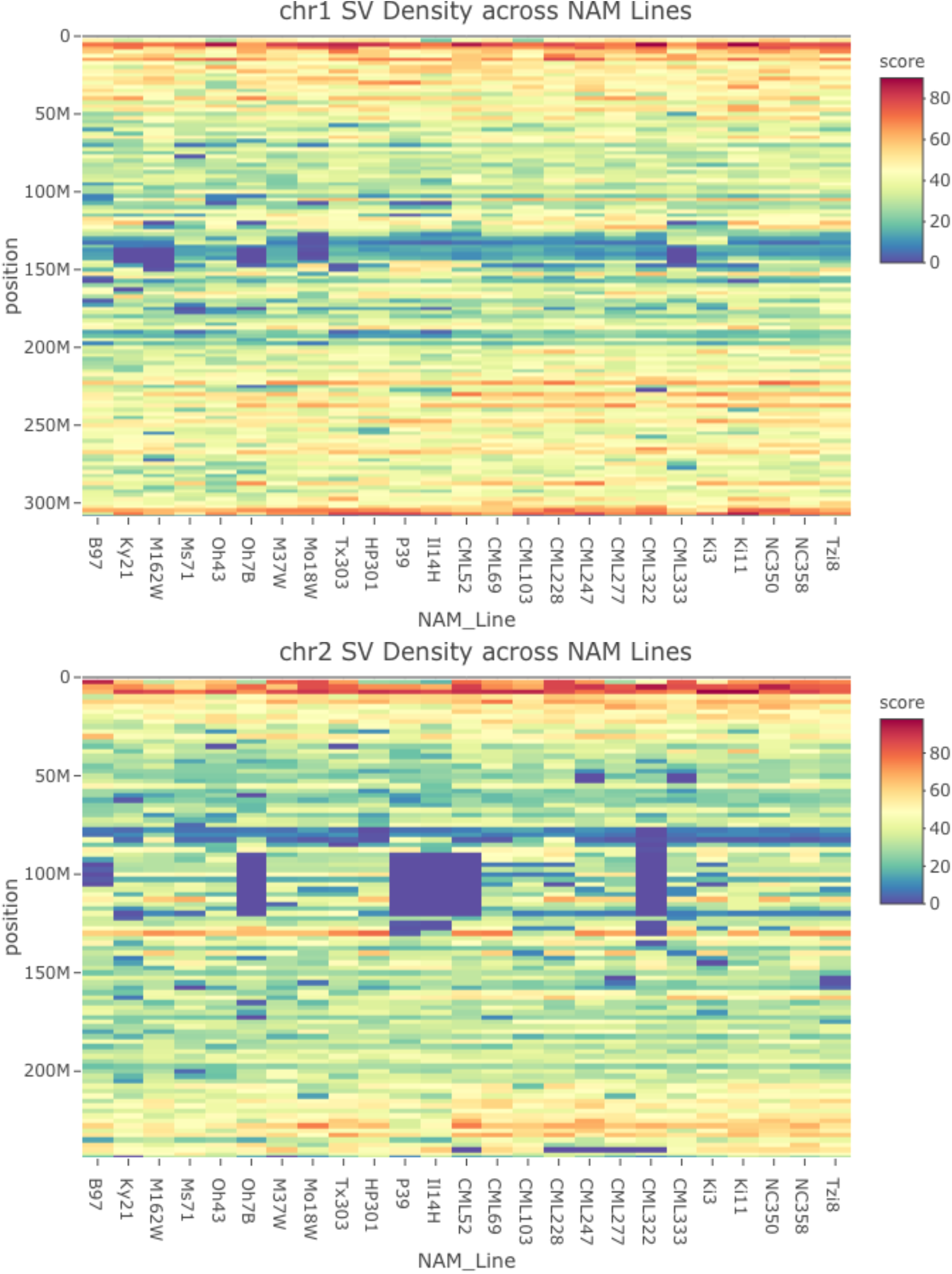
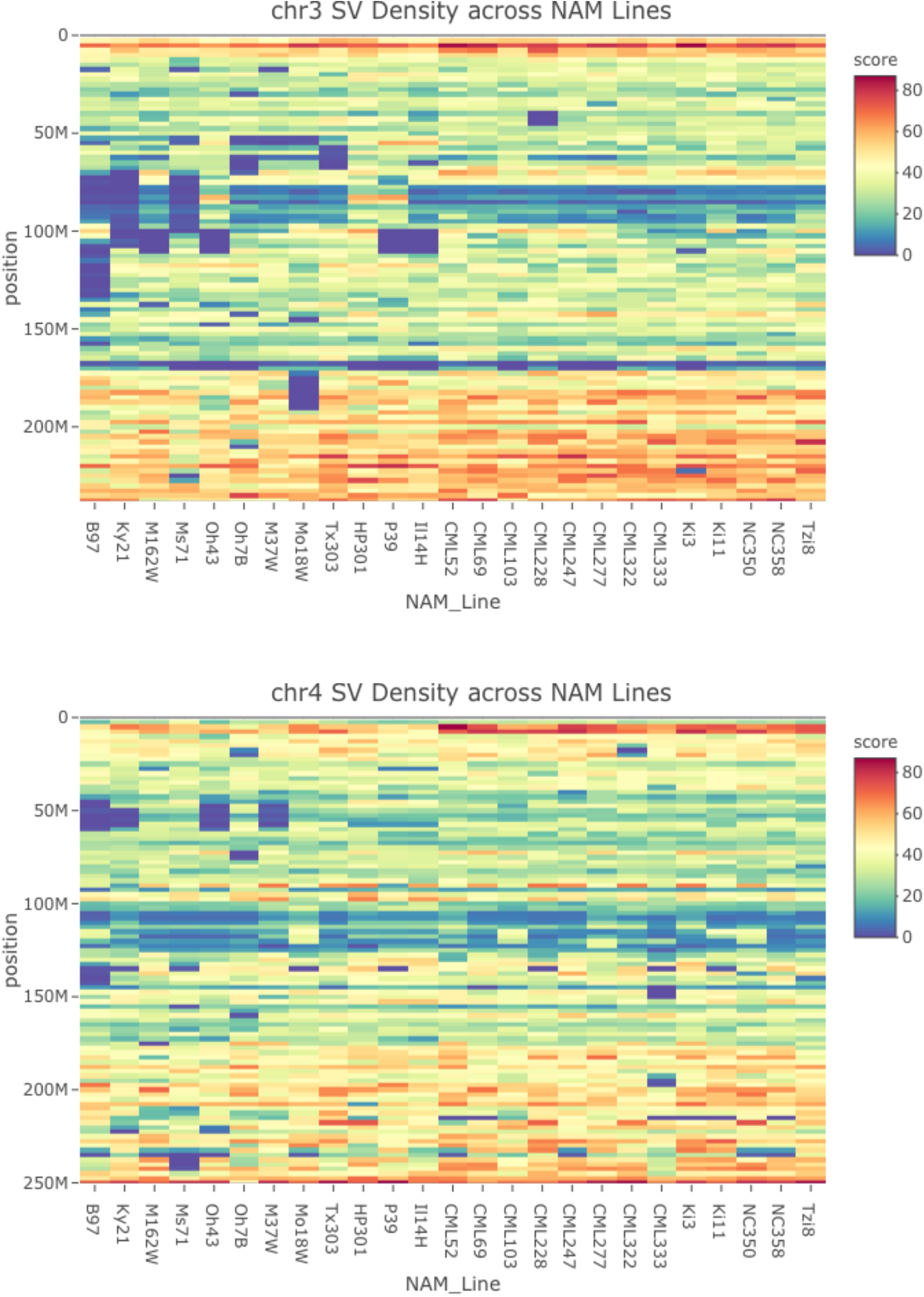
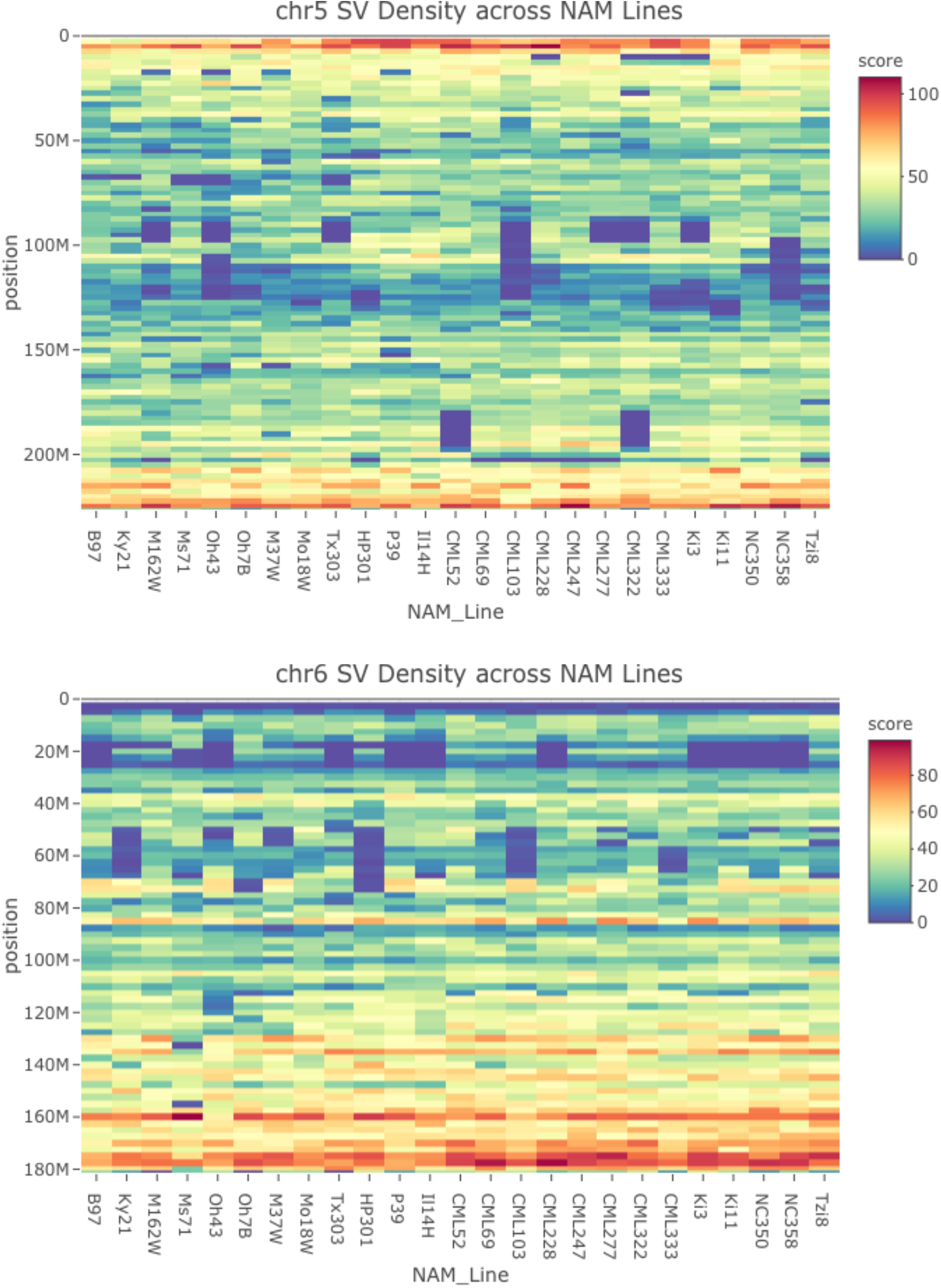
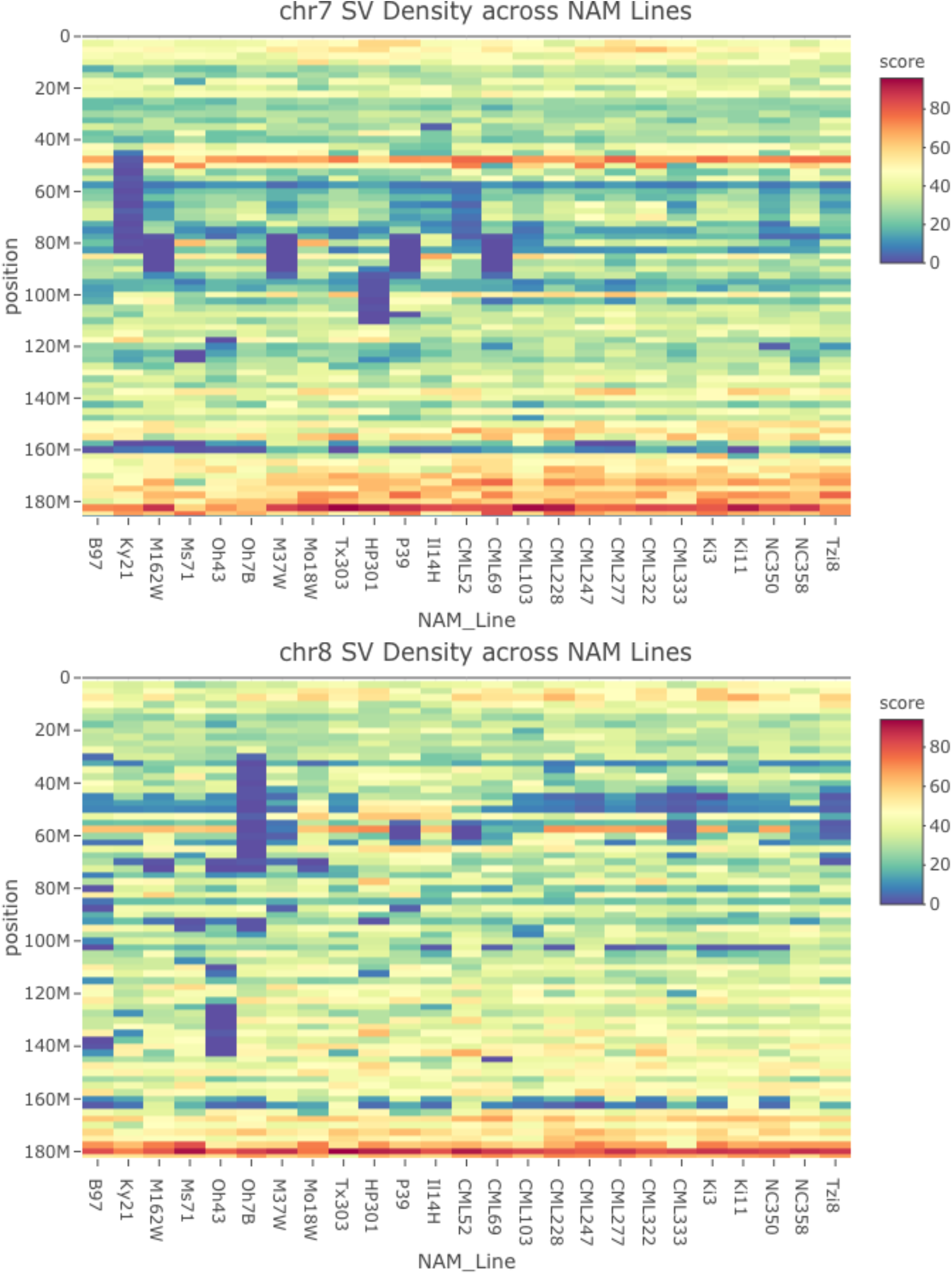
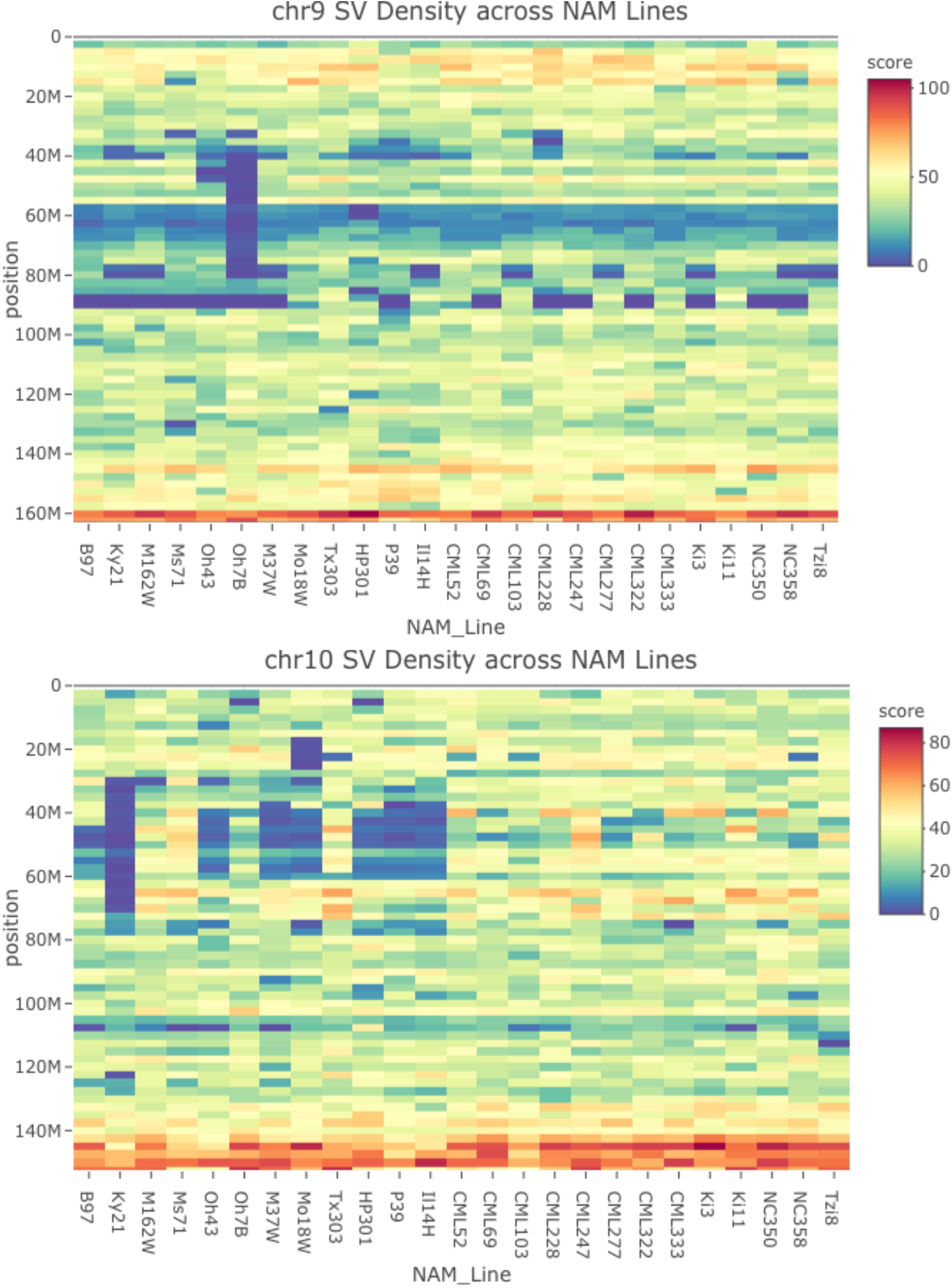
Density of structural variation across the NAM genome assemblies relative to the B73 genome. The scores represent the number of SVs per 2500kb. The minimum size of SVs in this analysis is 100bp or larger. Warmer color indicates higher density of SVs and cooler colors indicates lower density of SVs.

**Figure S11.**
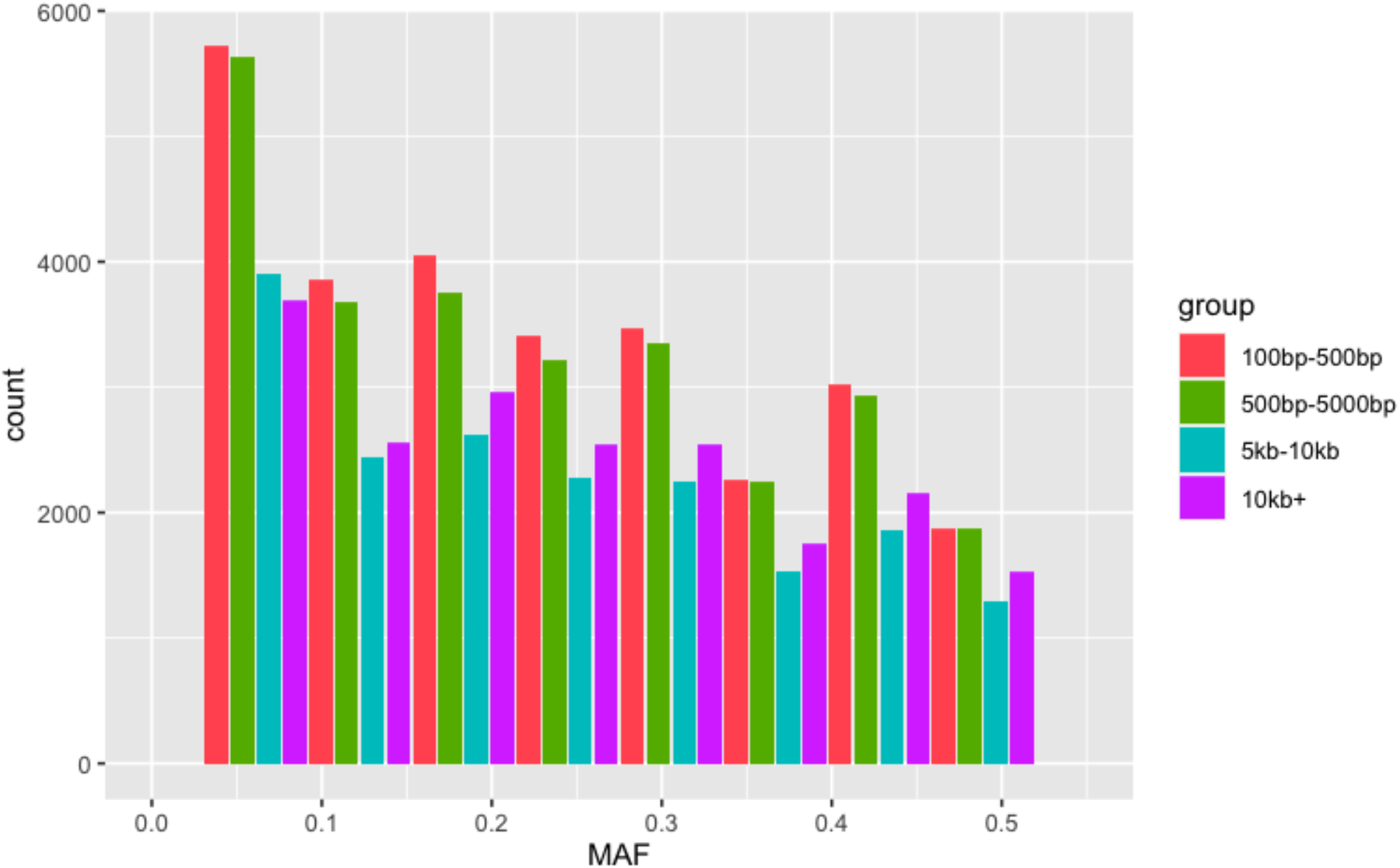
Distribution of structural variation across minor allele frequency (MAF) bins for various classes of size.

**Figure S12.**
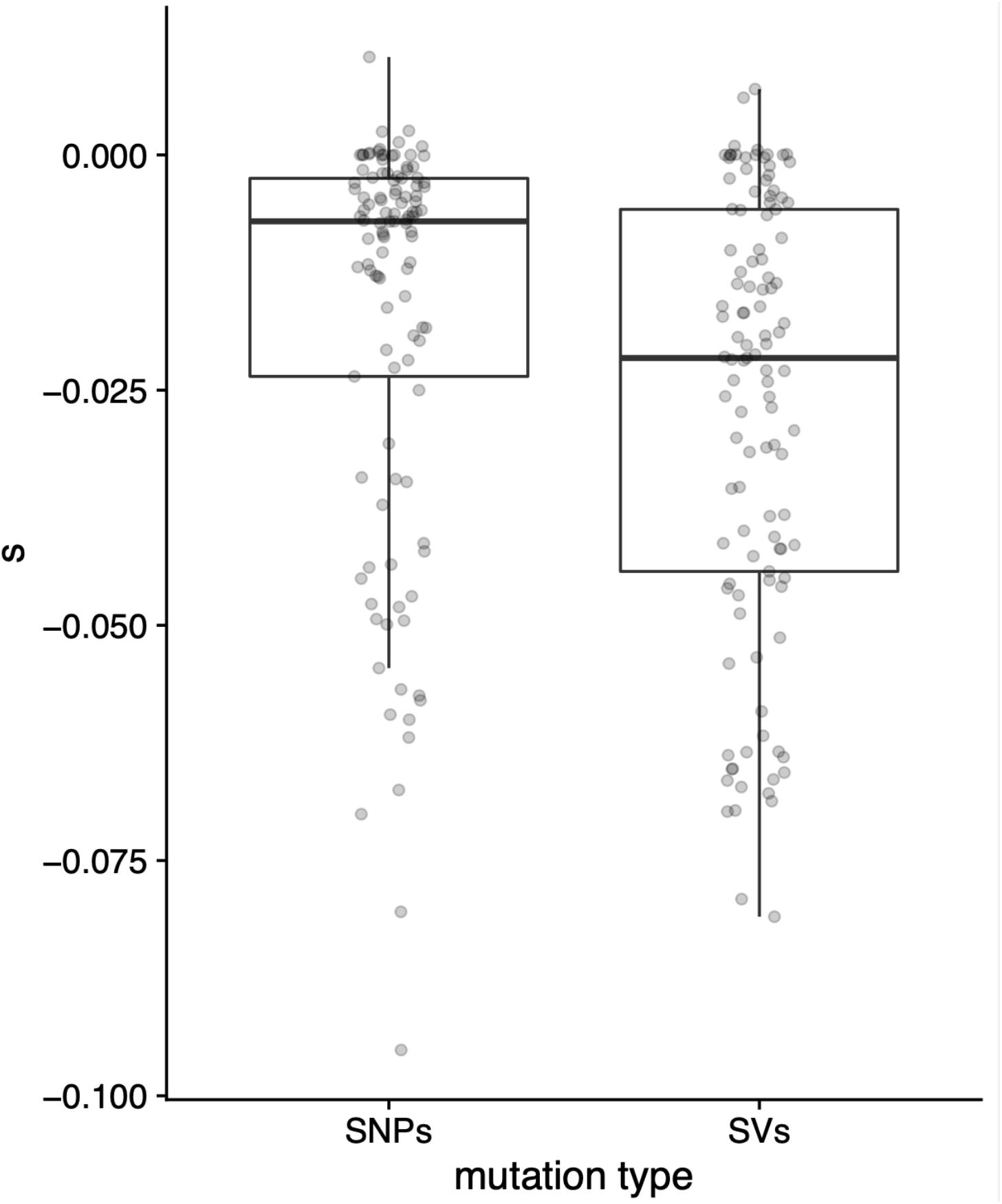
Distribution of mean selection coefficients (s) from 20-Mbp windows.

**Figure S13.**
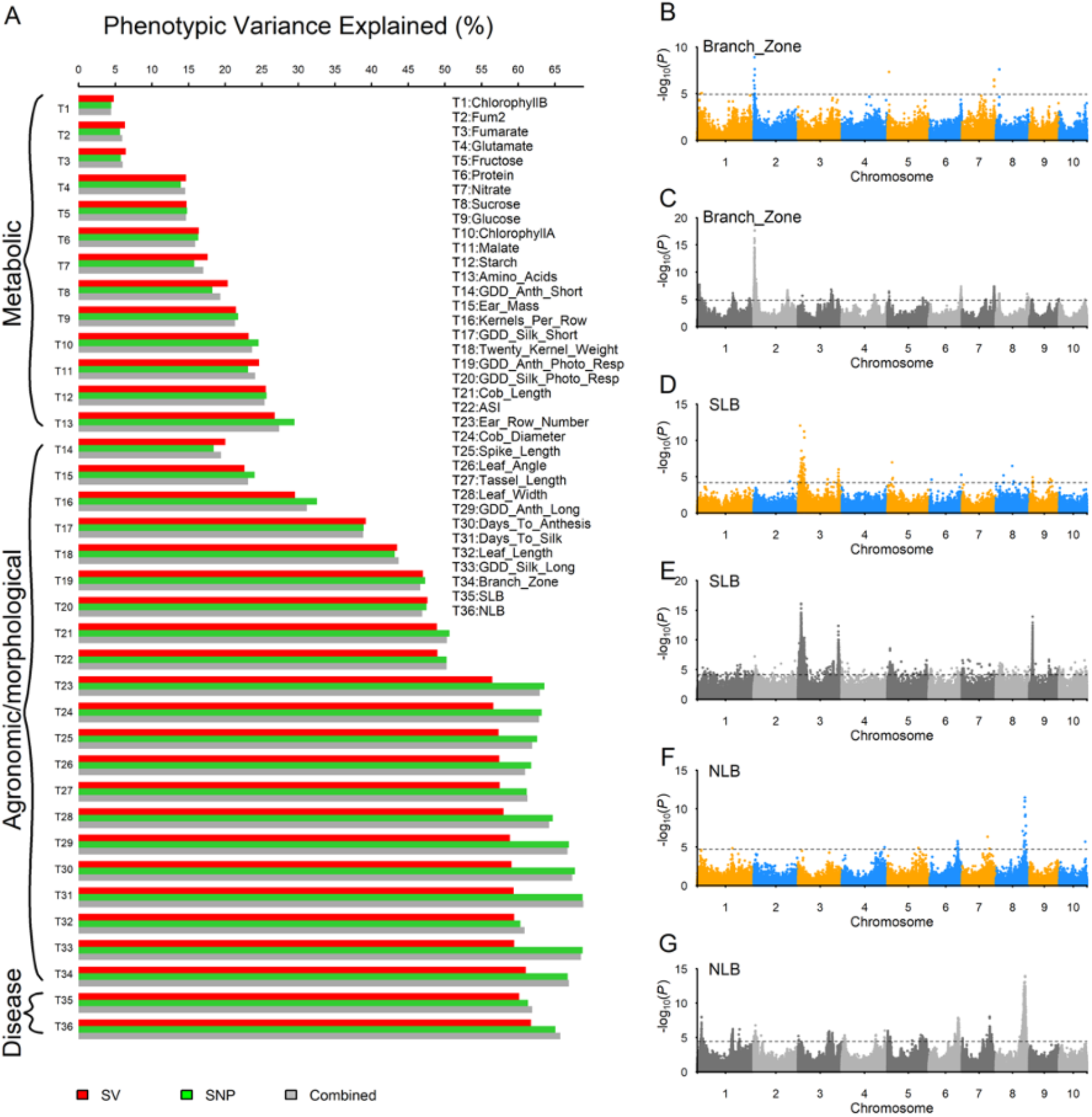
Genetic contributions from SVs and SNPs to complex traits in the population of NAM RILs. **A**) Phenotypic variance explained (PVE) by genome-wide SVs, SNPs, and combined. The 36 traits are organized into three groups: metabolic, agronomic/morphological, and disease-related traits. **B-G**) Manhattan plots of genome-wide association analyses (GWAS) of three traits with the highest PVEs by SVs. The GWAS with SVs (B, D, and F) detected significant QTLs, most of them overlapping with QTLs detected with SNPs (C, E, and G), but one on chromosome 10 for NLB was unique to SVs. The statistical significance thresholds on the Manhattan plots were obtained by controlling FDR on p-value 0.05. NLB and SLB are northern leaf blight and southern leaf blight, respectively.

**Figure S14.**
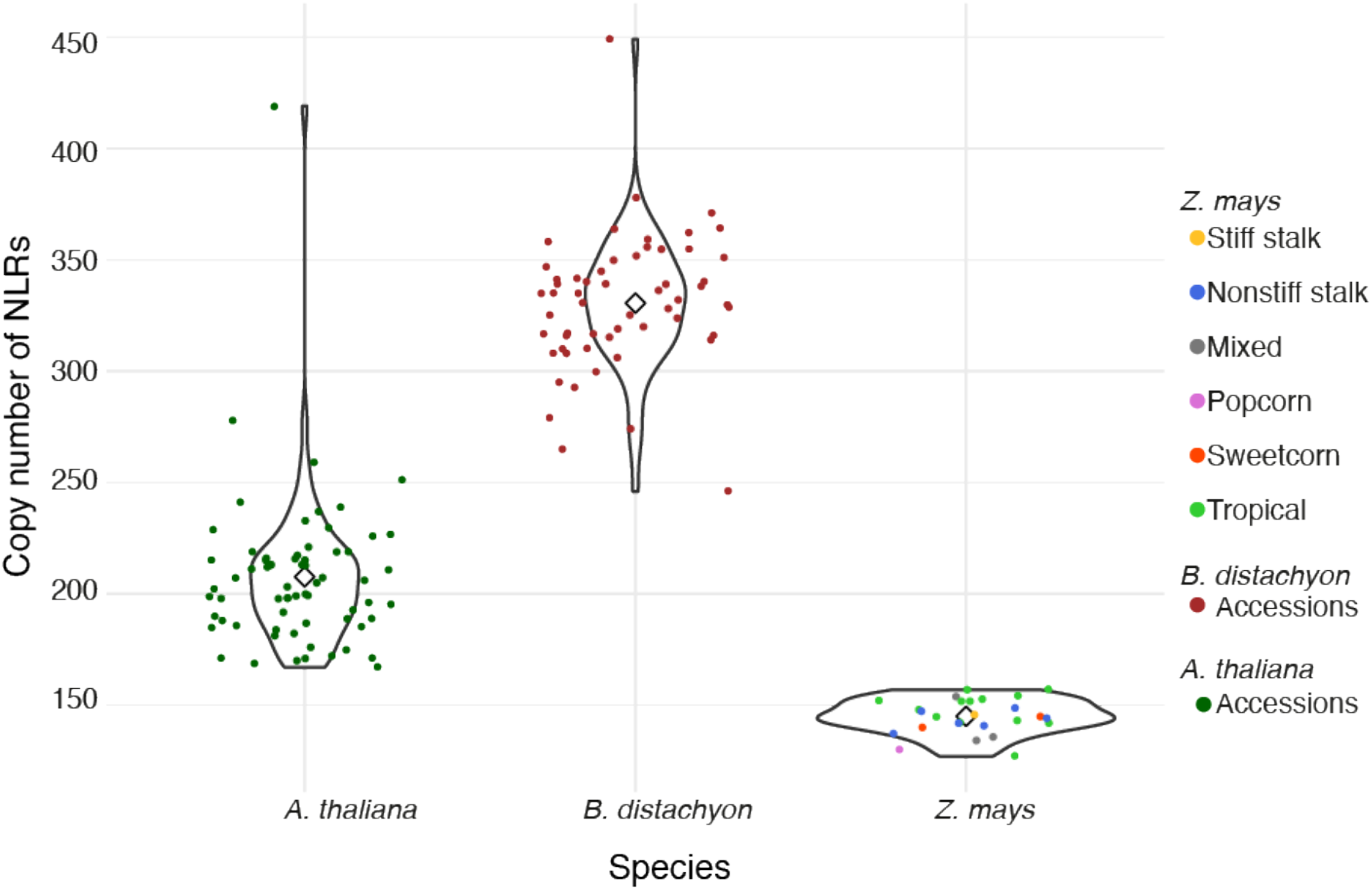
Violin plot of NLR variation in the pan-genomes of a eudicot (*A. thaliana*) and two monocot species (*B. distachyon and Z. mays*).

**Figure S15.**
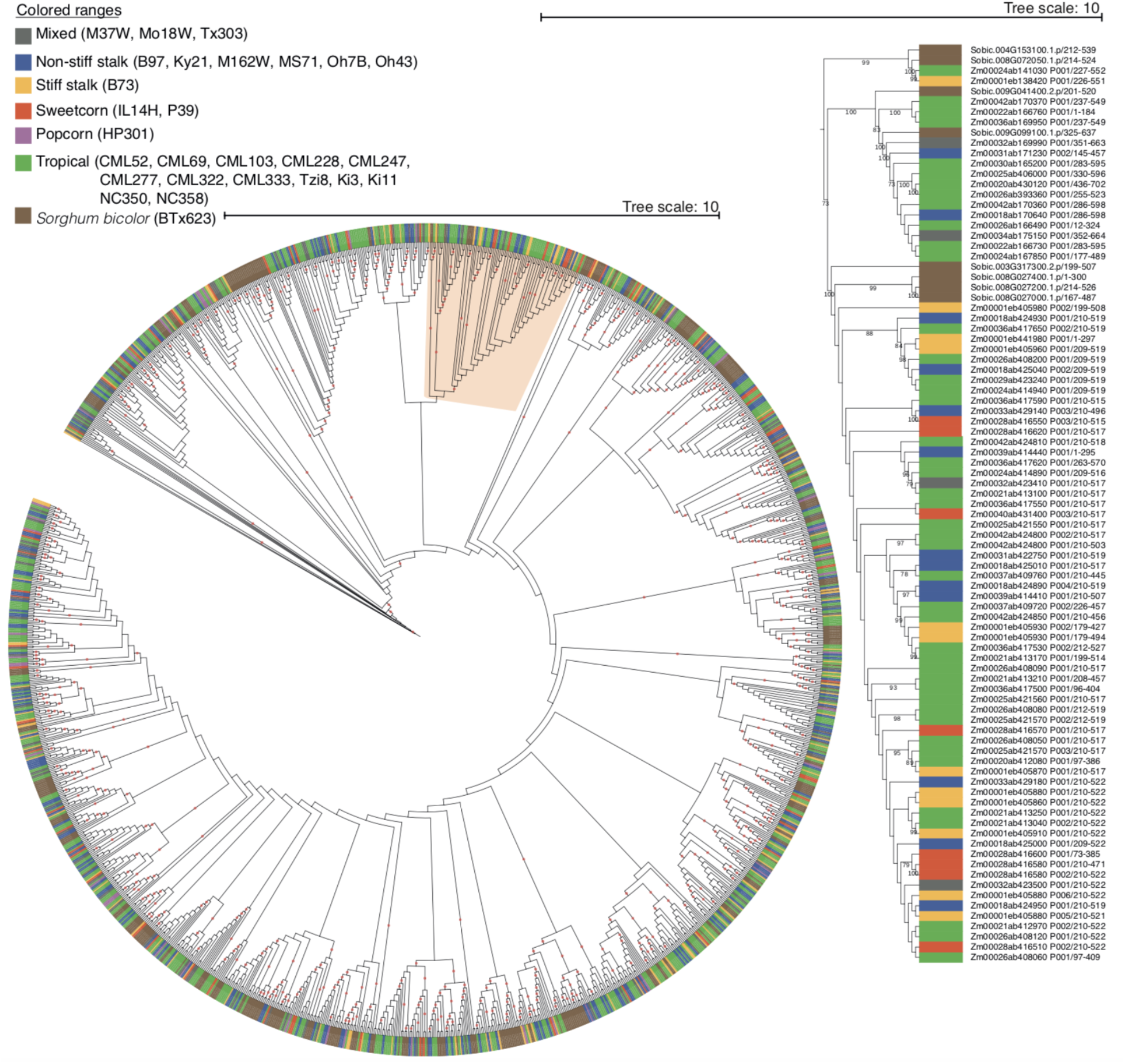
Maximum likelihood phylogeny of all NLR containing transcripts from NAM maize lines and *S. bicolor.* Dots indicate bootstrap values >80. The circle phylogeny shows all NAM NLRs. The linear phylogeny to the right is a zoom of the rose colored region illustrating the general trend that the NLR clades are broadly distributed across the maize NAM founder groups.

**Figure S16.**
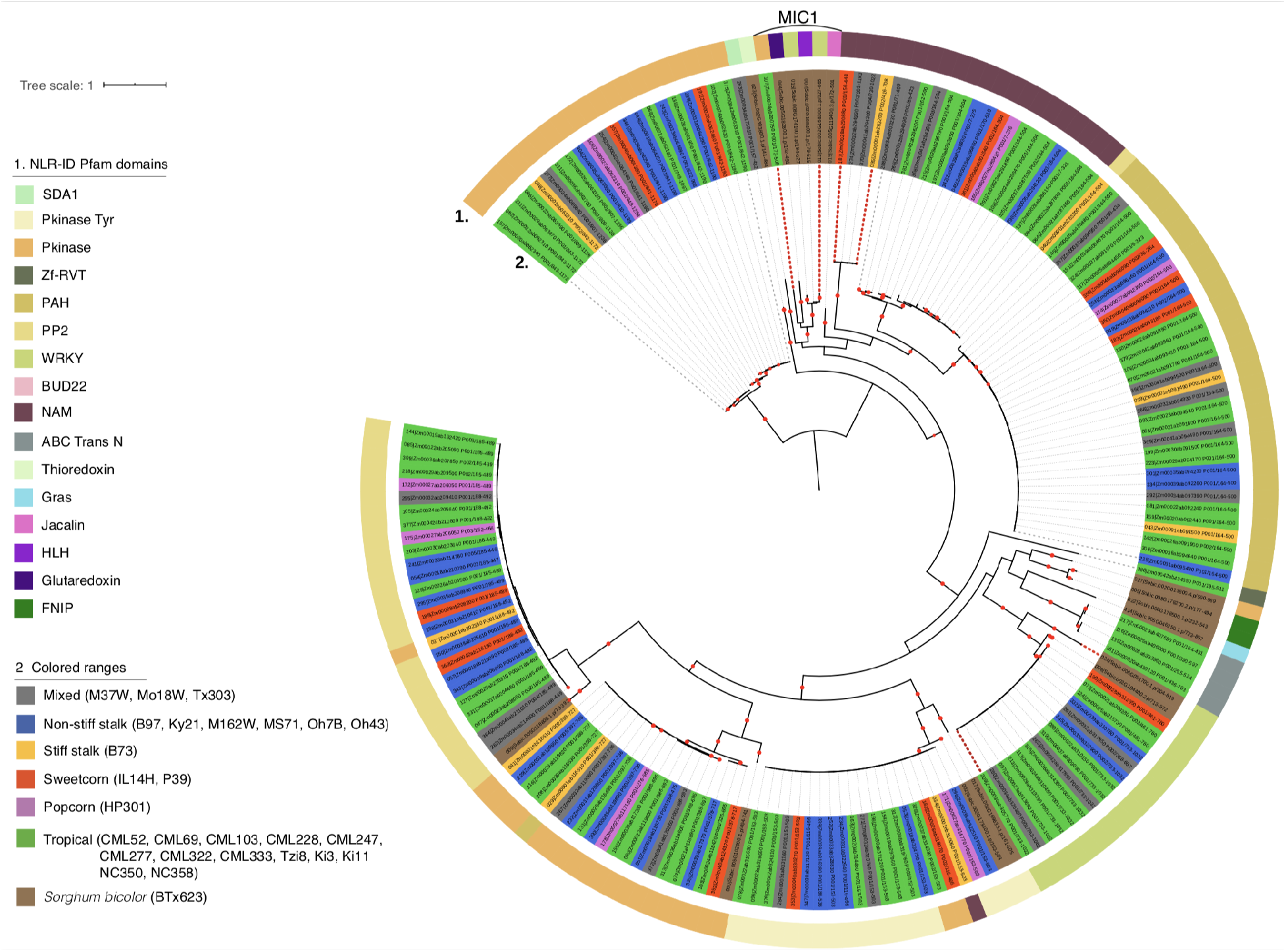
Maximum likelihood phylogeny of NLRs from NAM maize lines and *S. bicolor.* Single red dots on branches indicate bootstrap values >80. Ring one shows NLR-ID Pfam domains. Ring two shows the genes that have the corresponding Pfam domains. The colors represent the NAM founder groups (or Sorghum). Clades delimited by red dotted lines are segregating and not present in all NAM founders within a group.The MIC1 NLR clade (highlighted at top) is particularly fast-evolving in Poaceae (*49*).

**Figure S17.**
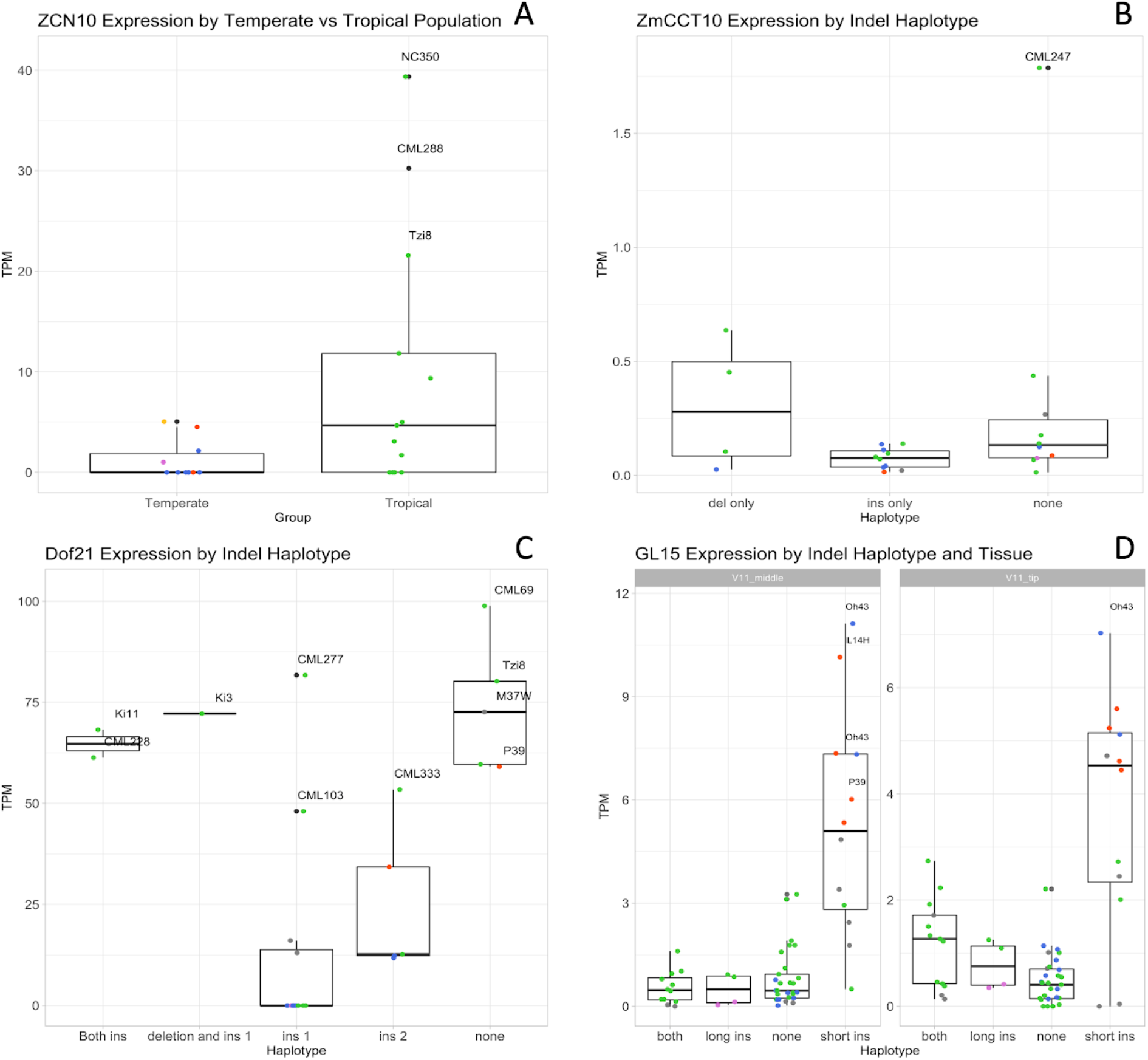
Candidate flowering time gene expression (Transcripts Per Million, TPM) by indel haplotypes. Point colors represent the population (green = tropical, orange = sweet corn, pink = popcorn, blue = non-stiff stalk, yellow = stiff stalk, and grey = mixed). For A-C, each point represents the average TPM across tissues and replicates. **A**) *ZCN10* expression in temperate and tropical groups (indel haplotype in the promoter region is unresolved). **B**) *ZmCCT10* expression in promoter haplotypes containing a deletion, an insertion, or neither indel. **C**) *Dof21* expression in promoter haplotypes containing two insertions and a deletion. **D**) *GL15* expression in promoter haplotypes with insertions. *GL15* showed significant expression differences in V11 middle (left) and tip leaf tissue (right), which was not detected when all tissues were averaged together. Since *GL15* is active for a short window of development in early vegetative stages, this fits with established knowledge of this gene. Expression is plotted based on the haplotypes created by presence/absence of a short and long insertion.

**Figure S18.**
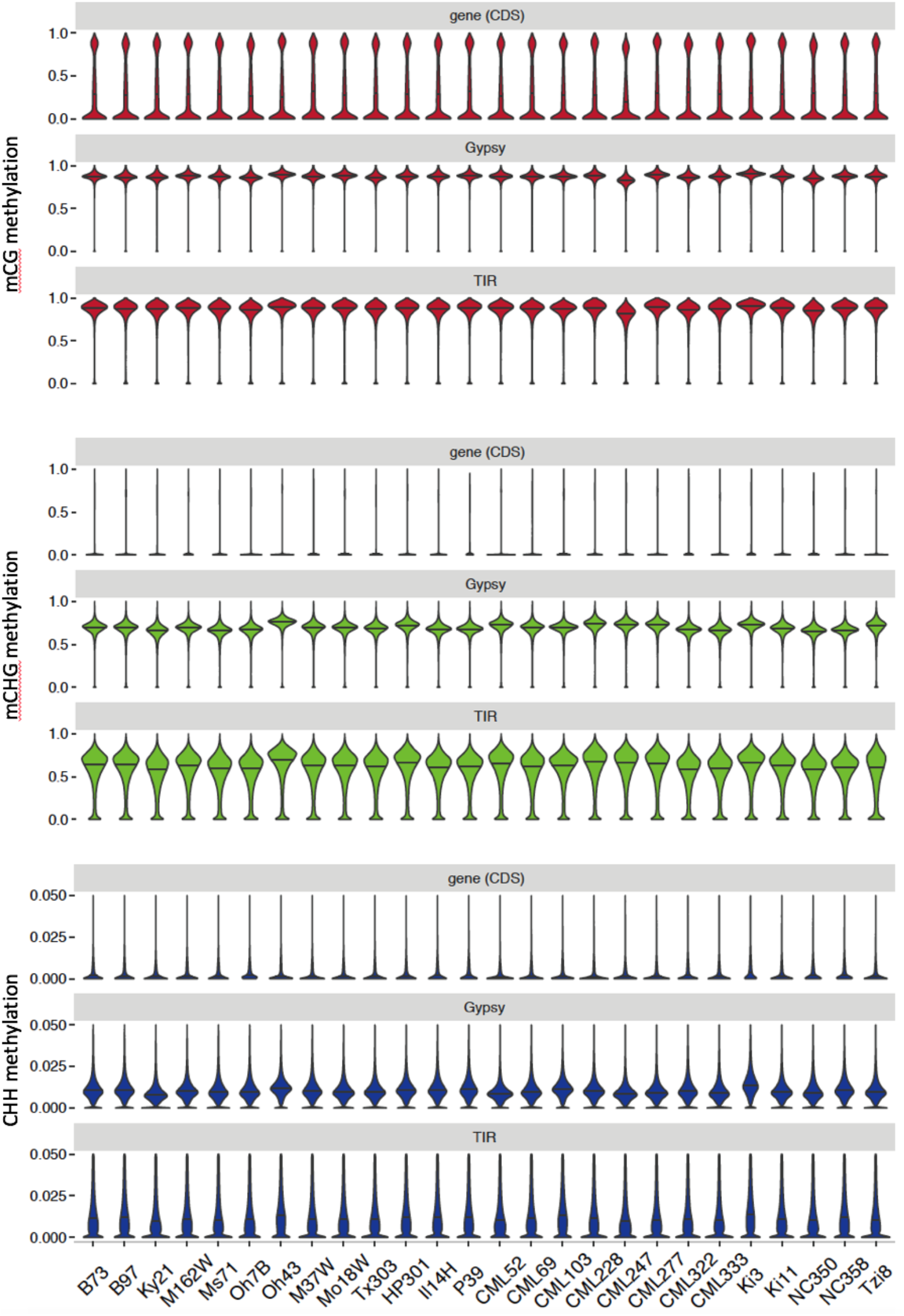
Spread of methylation levels for three representative genetic elements, genes (coding DNA only), *Gypsy* LTR retrotransposons, and TIR DNA transposons (*Tc1*/*Mariner*, *hAT*, *Harbinger*, and *Mutator*). Methylation is mC/total C for each sequence context. Horizontal lines indicate medians. To be included in this analysis, loci had to have a minimum of 10 cytosines in the specified context (CG, CHG, or CHH) that were covered by EM-seq reads. EM-seq reads from each methylome were mapped to their own genomes.

**Figure S19:**
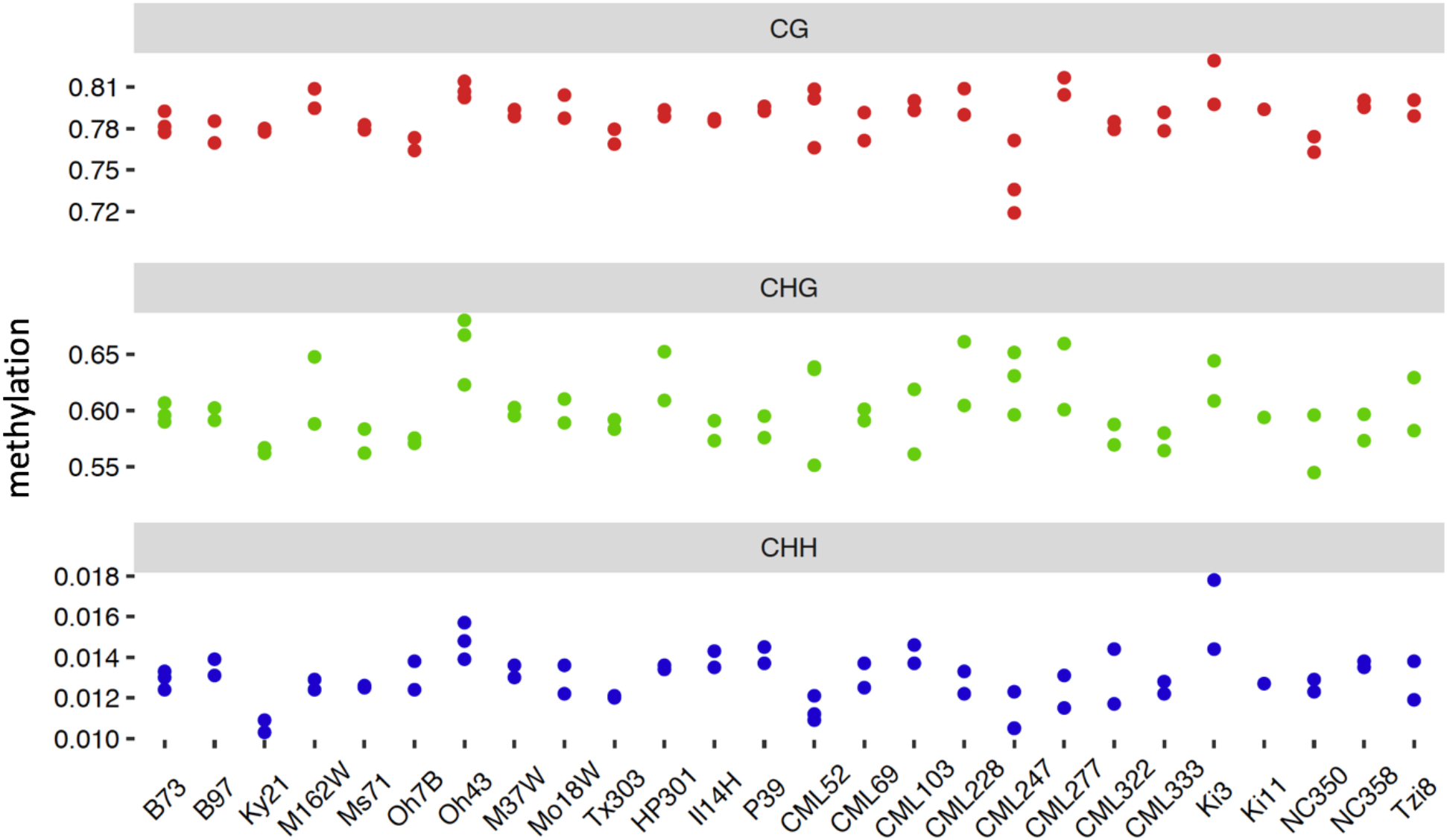
Whole-genome methylation levels for individual biological replicates. Methylation is mC/total C for each sequence context. EM-seq reads from each methylome were mapped to their own genomes.

**Figure S20.**
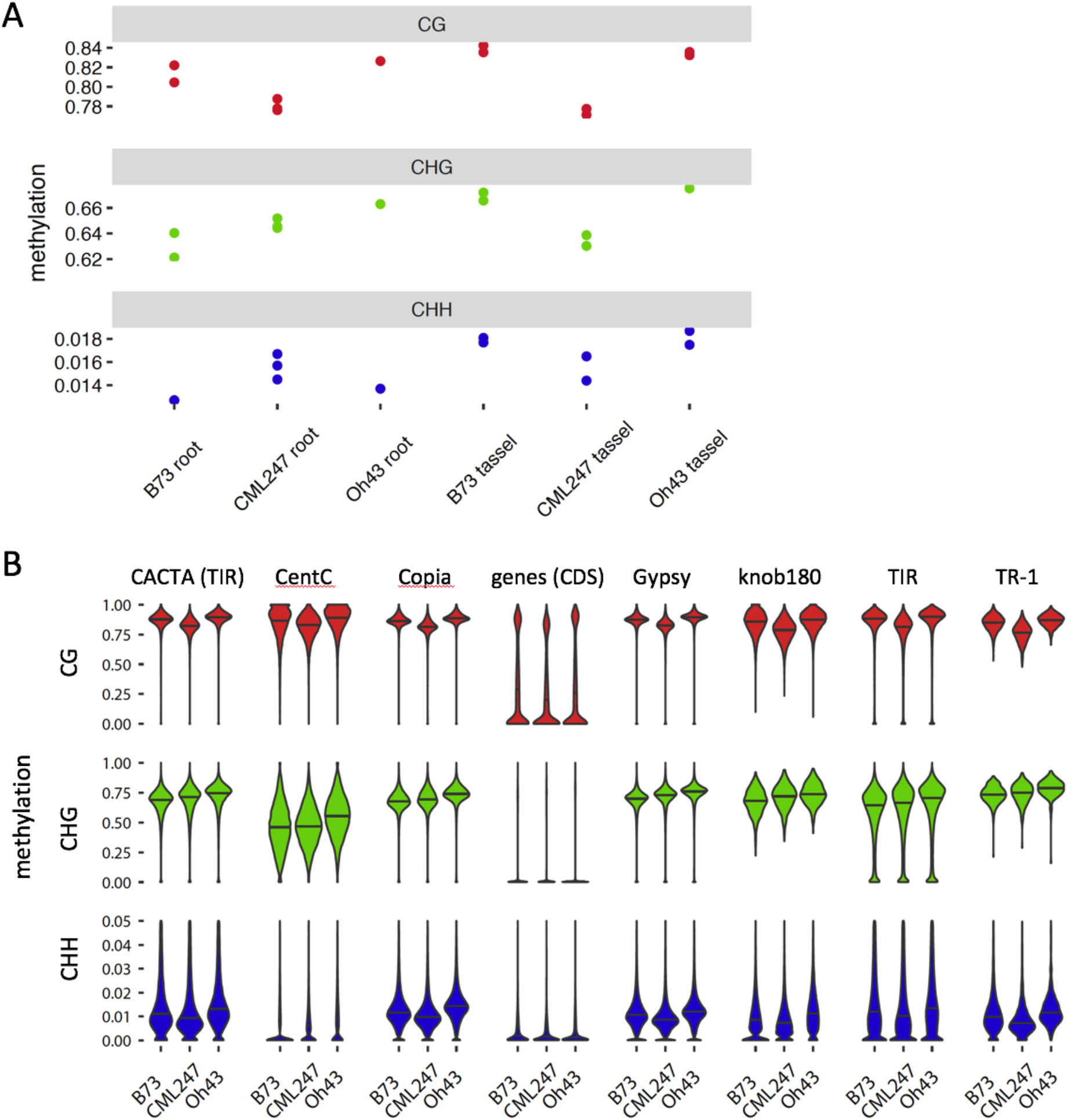
Additional comparisons of B73, CML247, and Oh43 methylation. **A**) Whole-genome methylation levels for individual biological replicates of primary root six days after planting and V18 growth stage meiotic tassel. Methylation is mC/total C for each sequence context. EM-seq reads from each methylome were mapped to their own genomes. **B**) Spread of methylation levels for representative genetic elements in developing second leaves. The same data are shown as in Fig. S18 but with the addition of five more genetic elements. Methylation is mC/total C for each sequence context. Horizontal lines indicate medians. All loci except CentC had to have a minimum of 10 cytosines in the specified context (CG, CHG, or CHH) that were covered by EM-seq reads. CentC was required to have 3 CGs or 5 CHGs. EM-seq reads from each methylome were mapped to their own genomes.

**Figure S21.**
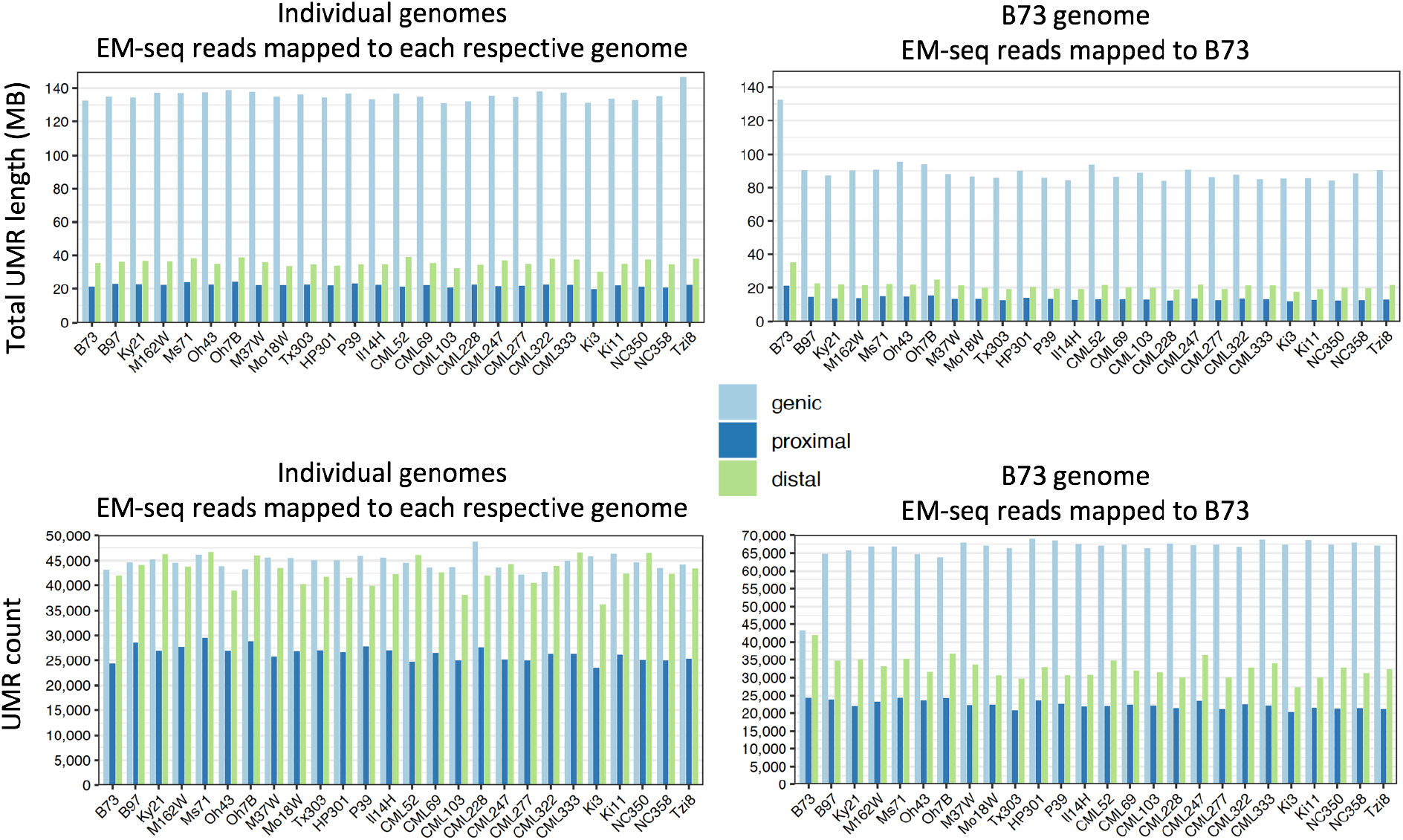
Total length and counts of UMRs. UMRs were defined relative to individual genomes by mapping each set of EM-seq reads to its own genome and defined relative to the B73 genome by mapping to B73. Position categories are as follows: UMRs with any overlap with genes are genic; of the remaining set, those with any overlap with the 5-KB flanks of genes are proximal; and the rest are distal.

**Figure S22.**
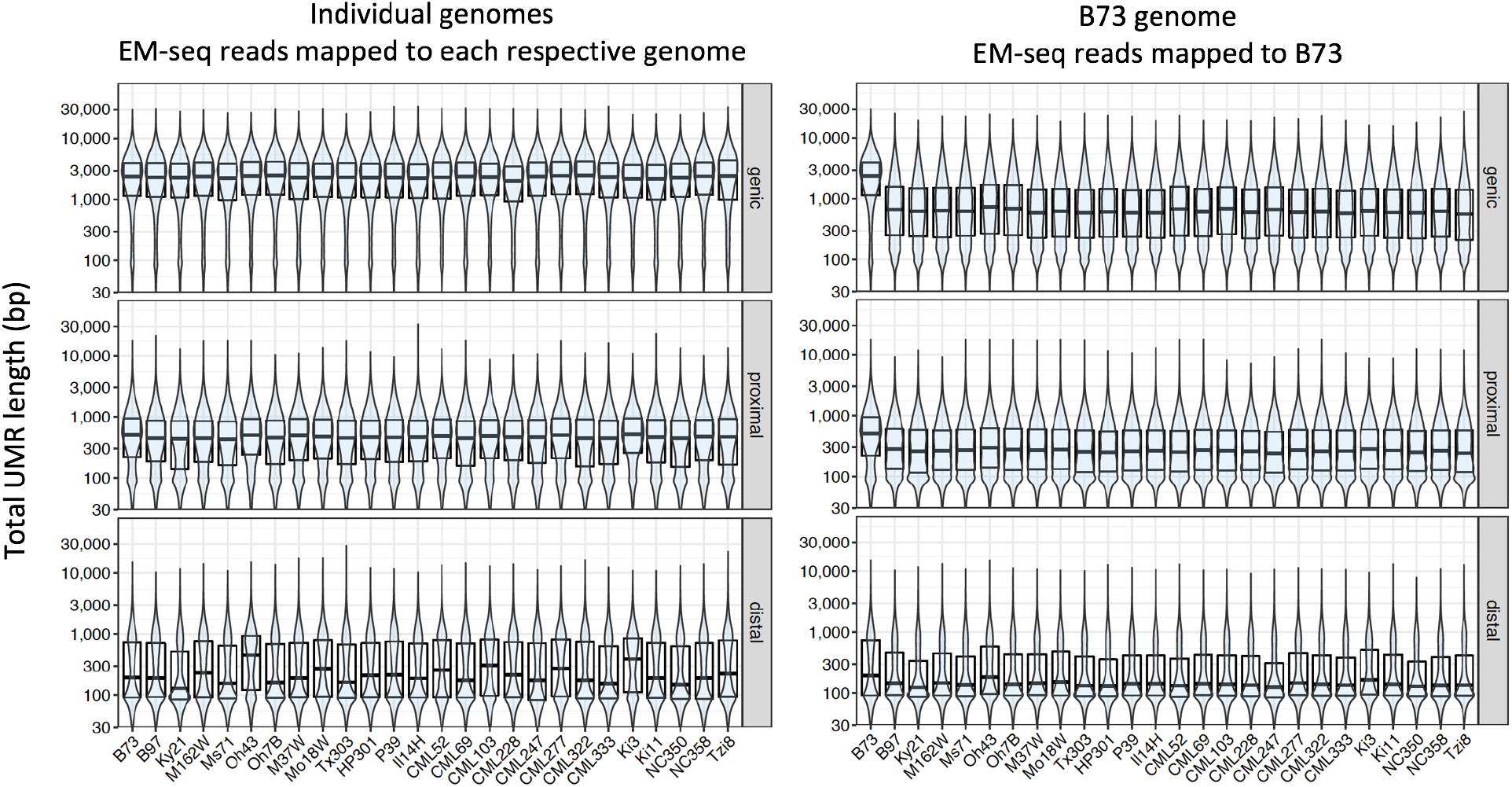
Spread of UMR lengths. UMRs were defined relative to individual genomes by mapping each set of EM-seq reads to its own genome and defined relative to the B73 genome by mapping to B73. Position categories are as follows: UMRs with any overlap with genes are genic; of the remaining set, those with any overlap with the 5-KB flanks of genes are proximal; and the rest are distal. This analysis includes UMRs that are less than 150 bp in length (which were excluded from all other analyses). Y-axes are on a log10 scale. Boxplots denote medians and quartiles.

**Figure S23.**
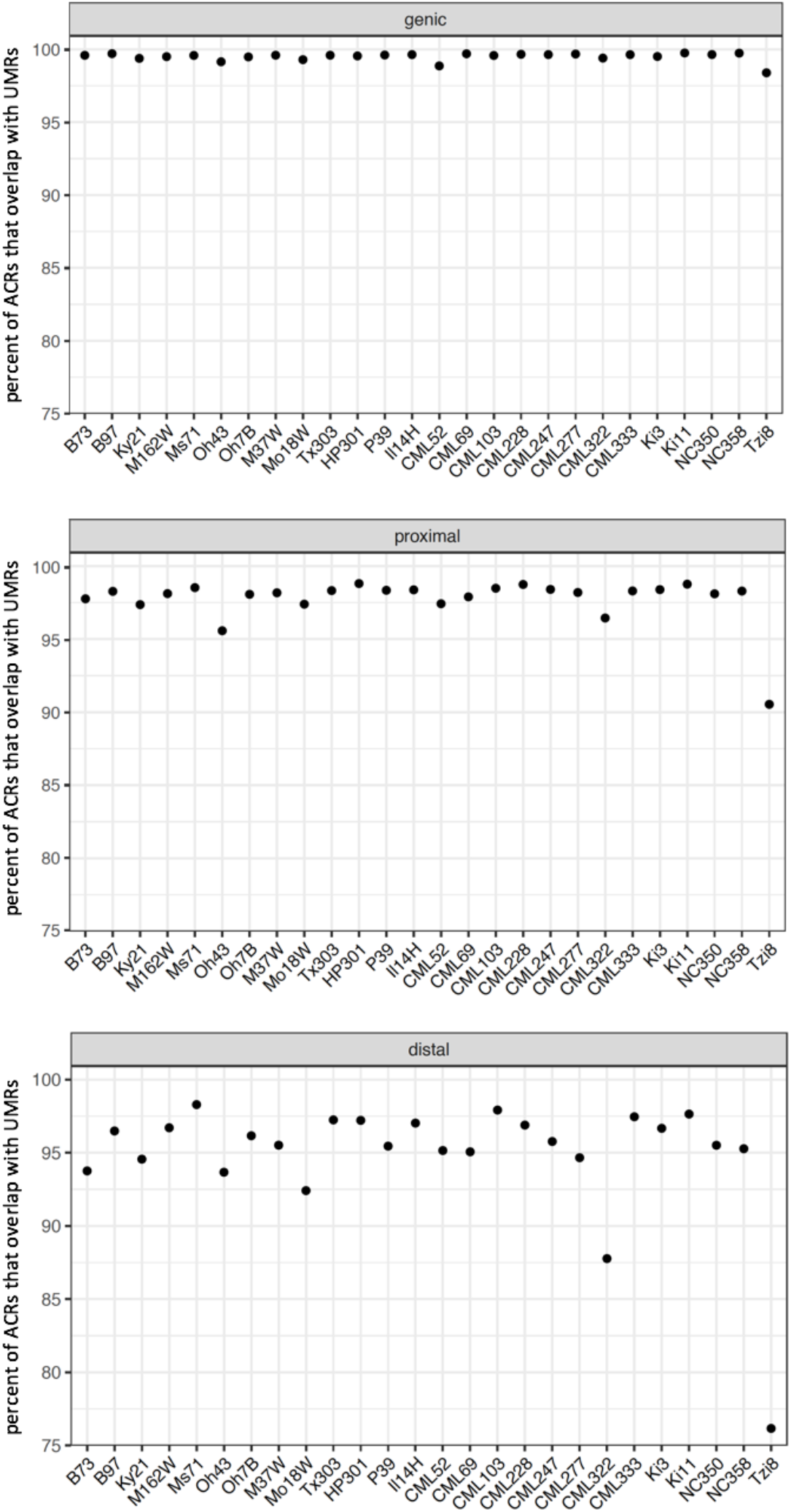
Overlaps of accessible chromatin regions (ACRs) by UMRs. Overlaps are >= 1 bp. UMRs and ACRs were defined relative to individual genomes by mapping each set of EM-seq and ATAC-seq reads to its own genome. Position categories are as follows: ACRs with any overlap with genes are genic; of the remaining set, those with any overlap with the 5-KB flanks of genes are proximal; and the rest are distal.

**Figure S24.**
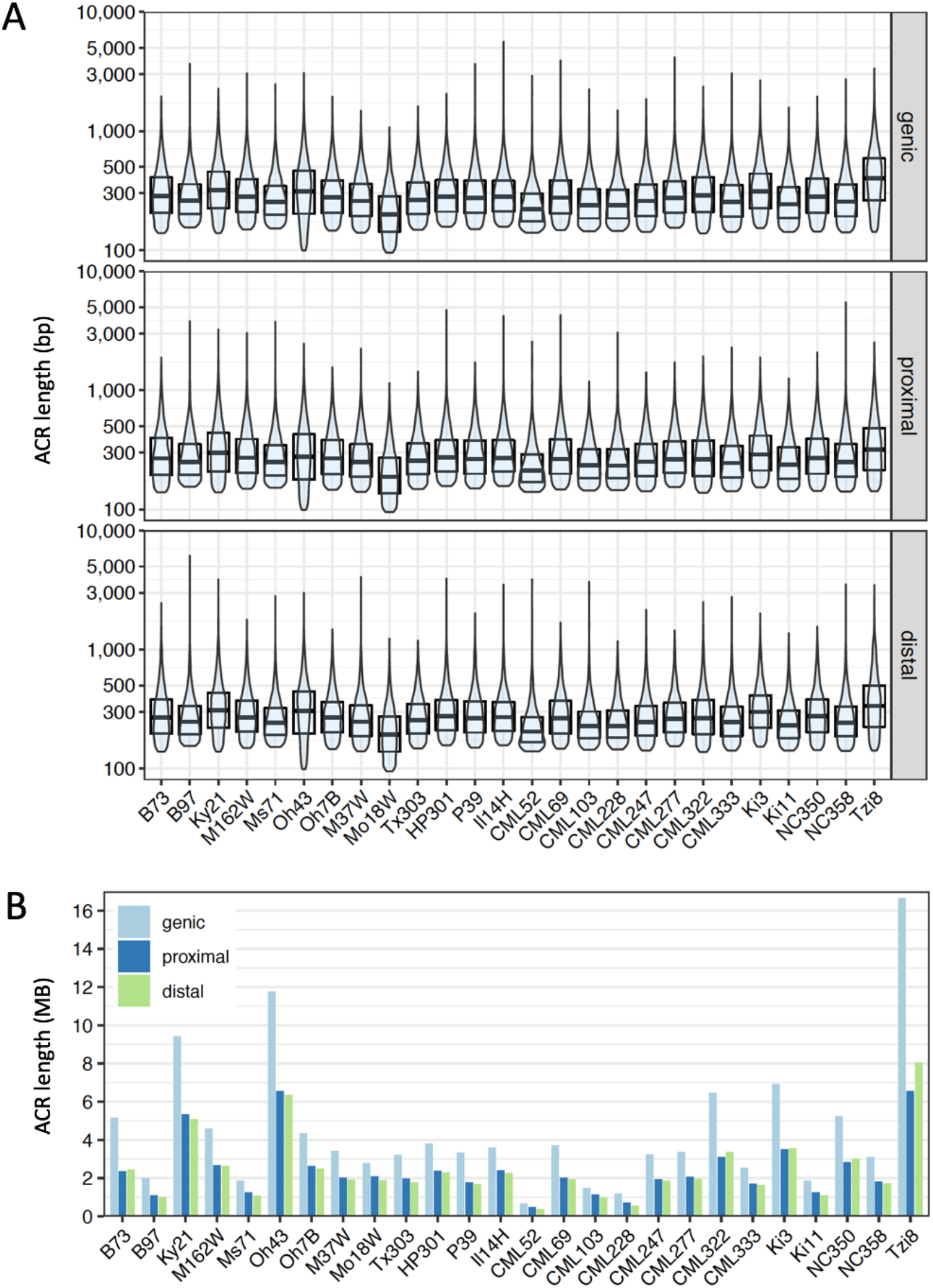
Lengths of Accessible Chromatin Regions (ACRs). **A**) Distributions of lengths of ACRs in each genome. Y-axes are on log10 scale. Position categories are as follows: ACRs/UMRs with any overlap with genes are genic; of the remaining set, those with any overlap with the 5-KB flanks of genes are proximal; and the rest are distal. Horizontal lines indicate medians. **B**) Cumulative length of ACRs in each genome.

**Figure S25.**
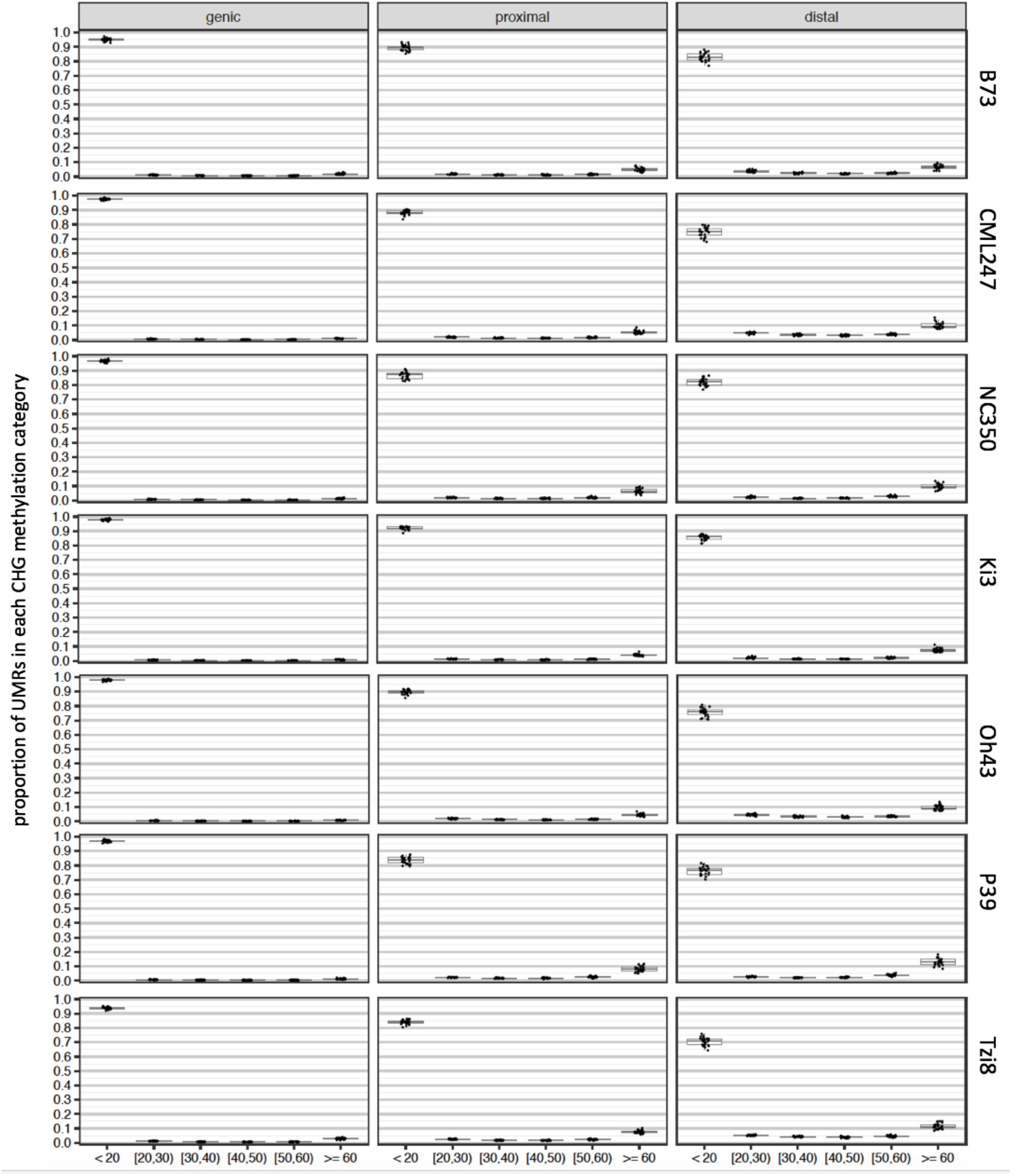
Conserved low CHG methylation in UMRs. UMRs were defined by mapping EM-seq reads from seven inbreds indicated at right. For each of the seven, UMRs were then categorized into one of six methylation bins (percent mCHG relative to total CHG) based on mapping EM-seq reads from the other 25 inbreds. Dots represent the proportion of the UMRs in each category. The “<20“ category is what was used to define UMRs. The data are further categorized based on position relative to genes: UMRs with any overlap with genes are genic; of the remaining set, UMRs with any overlap with the 5-kbp flanks of genes are proximal; and the rest are distal. Boxplots denote medians and quartiles. For these analyses, all EM-seq reads were mapped to the B73 genome.

**Figure S26.**
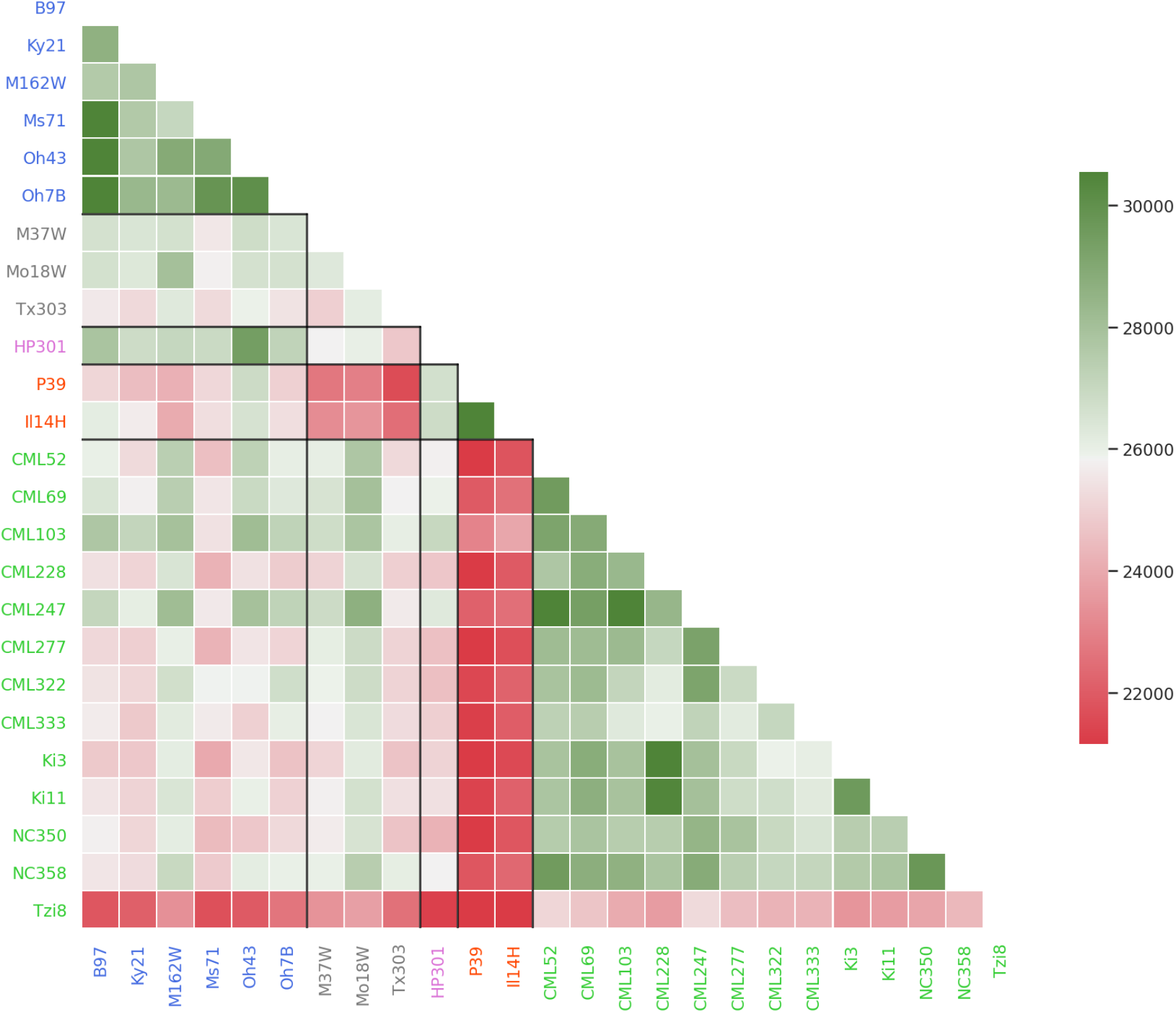
Heat map of the number of shared UMR regions across all pairwise comparisons of NAM lines. Boxed areas represent group by group comparisons.

**Figure S27.**
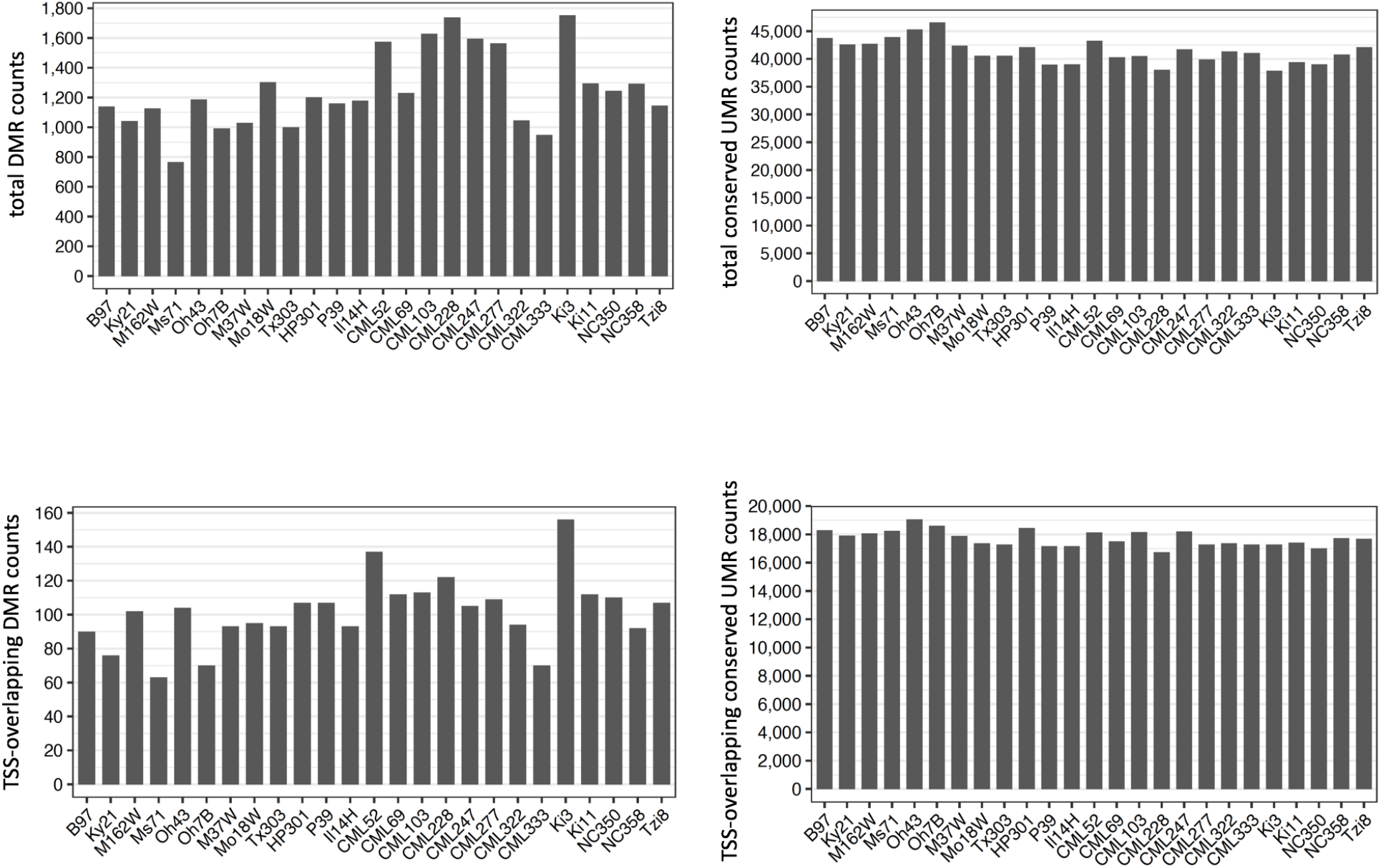
Numbers of differentially methylated regions (DMRs) in each NAM founder methylome. B73 UMRs that had greater than or equal to 60% CHG methylation in another methylome were categorized as DMRs, while B73 UMRs with less than 20% methylation in another methylome were categorized as conserved UMRs. Methylation was measured using the EM-seq reads from each methylome mapped to the B73 genome and was defined as percent mCHG relative to total CHG. A subset of TSS-overlapping pan-genes were selected as those where a region from -10 to +400 bp of the transcription start site was at least 98% overlapped by a DMR or conserved UMR.

## Supplementary Tables

**Table S1:** Accession and DNA isolation information for NAM lines

| Inbred Line | Original Accession | Sequenced Individual Accession | DNA Isolation Method |
| --- | --- | --- | --- |
| B73 | PI 550473 | PI 692136 | CTAB |
| B97 | PI 564682 | PI 692135 | nuclei |
| Ky21 | Ames 27130 | PI 692149 | CTAB |
| M162W | Ames 27134 | PI 692151 | nuclei |
| Ms71 | PI 587137 | PI 692153 | nuclei |
| Oh43 | Ames 19288 | PI 692157 | CTAB |
| Oh7B | Ames 19323 | PI 692156 | CTAB |
| M37W | Ames 27133 | PI 692150 | nuclei |
| Mo18W | PI 550441 | PI 692152 | nuclei |
| Tx303 | Ames 19327 | PI 692159 | CTAB |
| HP301 | PI 587131 | PI 692145 | CTAB |
| P39 | Ames 28186 | PI 692158 | CTAB |
| Il14H | Ames 27118 | PI 692146 | nuclei |
| CML52 | PI 595561 | PI 692137 | CTAB |
| CML69 | Ames 28184 | PI 692138 | CTAB |
| CML103 | Ames 27081 | PI 692139 | nuclei |
| CML228 | Ames 27088 | PI 692140 | CTAB |
| CML247 | PI 595541 | PI 692141 | CTAB |
| CML277 | PI 595550 | PI 692142 | CTAB |
| CML322 | Ames 27096 | PI 692143 | CTAB |
| CML333 | Ames 27101 | PI 692144 | nuclei |
| Ki3 | Ames 27123 | PI 692147 | CTAB |
| Ki11 | Ames 27124 | PI 692148 | CTAB |
| NC350 | Ames 27171 | PI 692154 | CTAB |
| NC358 | Ames 27175 | PI 692155 | CTAB |
| Tzi8 | PI 506246 | PI 692160 | CTAB |

**Table S2:** Quality metrics for the NAM genome assemblies.

|  | Assembly Size (Mb) | Placed Scaffolds Size (Mb) | Total Number of Scaffolds | Placed Scaffolds Number | Largest Scaffold (Mb) | Smallest Scaffold (Mb) | All Contigs N50 (Mb) | All Contigs L50 | All Scaffolds N50 (Mb) | All Scaffolds L50 | Placed Scaffolds N50 (Mb) | Placed Scaffolds L50 | Pseudomolecule %N | Placed Scaffolds %N | Placed Scaffolds BUSCO % | LAI |
| --- | --- | --- | --- | --- | --- | --- | --- | --- | --- | --- | --- | --- | --- | --- | --- | --- |
| **B73** | 2182.08 | 2132.8 | 695 | 20 | 242.93 | 0.0308 | 52.3551 | 13 | 160.85 | 6 | 162.82 | 5 | 0.17 | 0.13 | 95.76 | 27.84 |
| **B97** | 2193.12 | 2135.16 | 817 | 27 | 236.26 | 0.0303 | 49.7671 | 12 | 137.68 | 6 | 137.68 | 6 | 0.15 | 0.16 | 95.69 | 28.06 |
| **Ky21** | 2171.65 | 2121.14 | 664 | 27 | 238.07 | 0.03 | 19.0707 | 31 | 115.38 | 7 | 115.38 | 7 | 0.3 | 0.31 | 96.04 | 28.08 |
| **M162W** | 2184.33 | 2129.28 | 813 | 27 | 237.44 | 0.03 | 27.8072 | 22 | 111.38 | 6 | 111.38 | 6 | 0.17 | 0.18 | 96.04 | 28.09 |
| **Ms71** | 2214.05 | 2128.11 | 1034 | 32 | 202.96 | 0.03 | 34.1022 | 19 | 98.45 | 9 | 101.56 | 8 | 0.22 | 0.21 | 95.97 | 27.91 |
| **Oh43** | 2176.4 | 2116.4 | 777 | 30 | 198.84 | 0.03 | 28.631 | 26 | 105.60 | 8 | 105.60 | 8 | 0.12 | 0.12 | 95.76 | 27.89 |
| **Oh7B** | 2164.7 | 2124 | 634 | 29 | 239.51 | 0.0302 | 13.62 | 41 | 140.10 | 6 | 140.10 | 6 | 0.4 | 0.41 | 95.80 | 28.04 |
| **M37W** | 2192.4 | 2144.22 | 758 | 29 | 234.27 | 0.0301 | 39.6241 | 17 | 105.37 | 7 | 105.37 | 7 | 0.08 | 0.09 | 95.97 | 28.09 |
| **Mo18W** | 2223.23 | 2132.06 | 1097 | 31 | 201.21 | 0.0302 | 24.9769 | 26 | 111.10 | 7 | 111.10 | 7 | 0.33 | 0.33 | 96.60 | 27.81 |
| **Tx303** | 2215.81 | 2124.95 | 1067 | 32 | 240.2 | 0.0302 | 27.971 | 24 | 99.16 | 8 | 99.16 | 8 | 0.31 | 0.26 | 95.83 | 27.71 |
| **HP301** | 2140.95 | 2114.55 | 408 | 23 | 233.97 | 0.0304 | 35.6 | 21 | 135.87 | 6 | 135.87 | 6 | 0.19 | 0.2 | 95.63 | 28.05 |
| **P39** | 2138.71 | 2104.38 | 424 | 21 | 249.96 | 0.0305 | 35.7844 | 21 | 147.88 | 6 | 147.88 | 6 | 0.11 | 0.12 | 95.76 | 27.61 |
| **II14H** | 2124.54 | 2102.61 | 297 | 25 | 238.22 | 0.03 | 19.6425 | 31 | 135.80 | 6 | 135.80 | 6 | 0.15 | 0.16 | 95.63 | 27.83 |
| **CML52** | 2307.69 | 2149.15 | 1708 | 36 | 200.09 | 0.03 | 11.2031 | 58 | 92.05 | 9 | 94.80 | 8 | 0.93 | 0.96 | 95.76 | 27.92 |
| **CML69** | 2224.9 | 2133.04 | 1352 | 31 | 239.36 | 0.0301 | 21.3376 | 31 | 107.57 | 7 | 107.57 | 7 | 0.29 | 0.31 | 95.90 | 28.34 |
| **CML103** | 2162.44 | 2117.65 | 648 | 26 | 234.9 | 0.0301 | 11.3391 | 59 | 129.92 | 6 | 129.92 | 6 | 0.24 | 0.25 | 95.56 | 28.3 |
| **CML228** | 2300.77 | 2145.83 | 1555 | 32 | 240.54 | 0.03 | 9.55255 | 57 | 108.07 | 7 | 140.27 | 6 | 1.2 | 1.08 | 96.25 | 27.93 |
| **CML247** | 2214.75 | 2142.86 | 1091 | 32 | 200.43 | 0.0301 | 11.4266 | 60 | 101.10 | 9 | 101.77 | 8 | 0.45 | 0.47 | 96.32 | 28.44 |
| **CML277** | 2190.8 | 2133.12 | 928 | 32 | 194.76 | 0.0301 | 6.2549 | 100 | 98.85 | 9 | 98.85 | 9 | 0.42 | 0.44 | 96.18 | 27.95 |
| **CML322** | 2219.25 | 2120.08 | 1290 | 32 | 240.12 | 0.0301 | 30.4894 | 24 | 102.20 | 8 | 104.77 | 7 | 0.14 | 0.14 | 95.42 | 28.44 |
| **CML333** | 2231.26 | 2141.31 | 1116 | 34 | 236.82 | 0.03 | 28.8179 | 20 | 99.84 | 8 | 99.84 | 8 | 0.14 | 0.11 | 96.32 | 28.33 |
| **KI3** | 2215.86 | 2139.35 | 1184 | 29 | 244.14 | 0.0301 | 16.1777 | 43 | 107.93 | 7 | 107.93 | 7 | 0.39 | 0.41 | 96.67 | 28.27 |
| **KI11** | 2273.84 | 2151.7 | 1417 | 29 | 240.57 | 0.0301 | 31.4035 | 23 | 110.07 | 6 | 200.72 | 5 | 0.12 | 0.13 | 95.83 | 27.64 |
| **NC350** | 2290.5 | 2163.61 | 1399 | 30 | 200.52 | 0.0303 | 49.0042 | 13 | 100.66 | 8 | 102.34 | 7 | 0.07 | 0.07 | 96.18 | 27.96 |
| **NC358** | 2227.42 | 2126.56 | 1335 | 32 | 237.4 | 0.03 | 25.9366 | 27 | 98.95 | 8 | 99.55 | 7 | 0.17 | 0.16 | 96.18 | 28.22 |
| **Tzi8** | 2271.03 | 2134.06 | 1570 | 32 | 239.1 | 0.03 | 11.6145 | 54 | 100.58 | 8 | 101.61 | 7 | 0.56 | 0.53 | 96.11 | 27.9 |

**Table S3:**
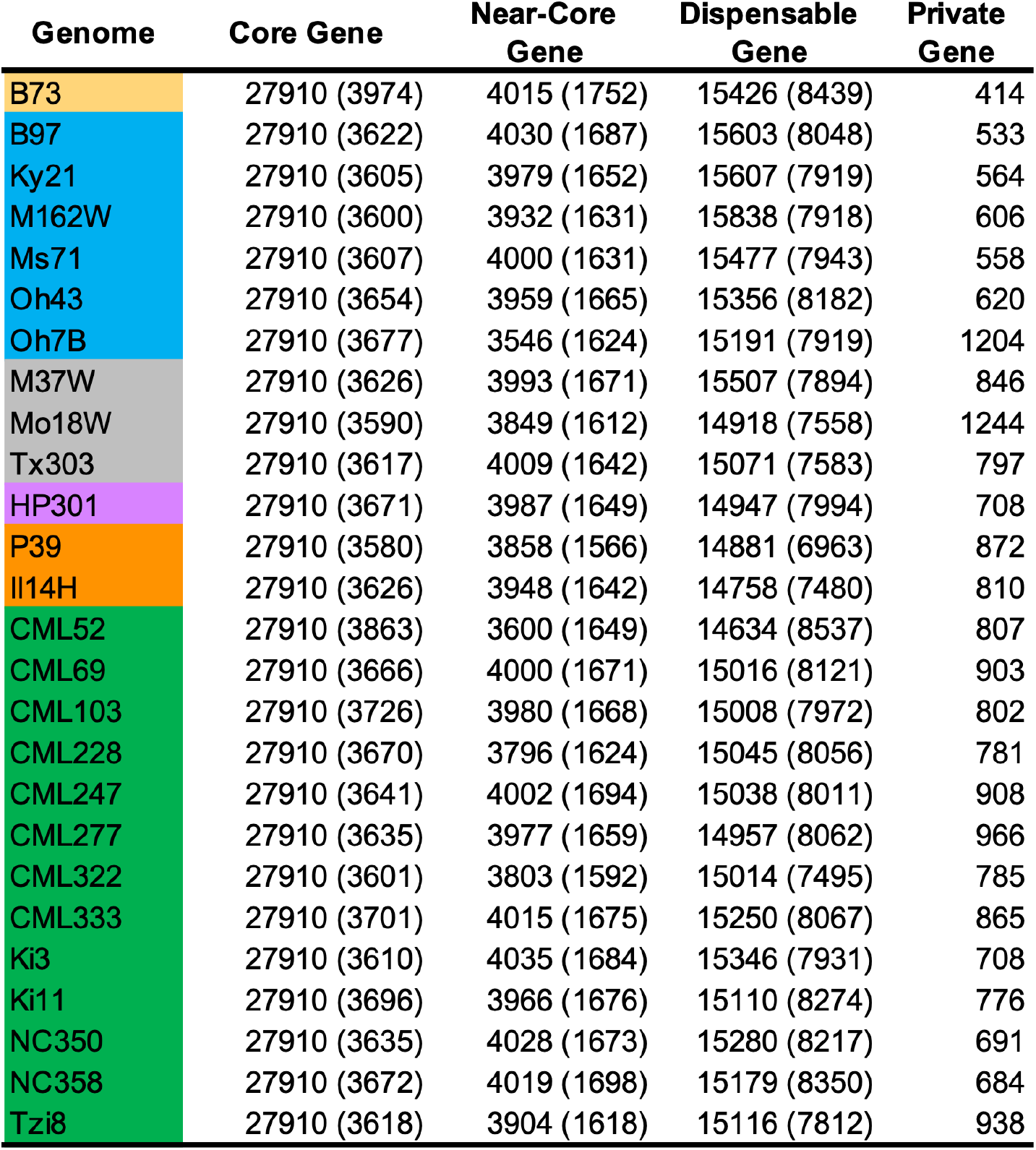
Categorization of pan-genes for the NAM genomes. Numbers in parentheses are identified based on coordinate filling.

**Table S4:**
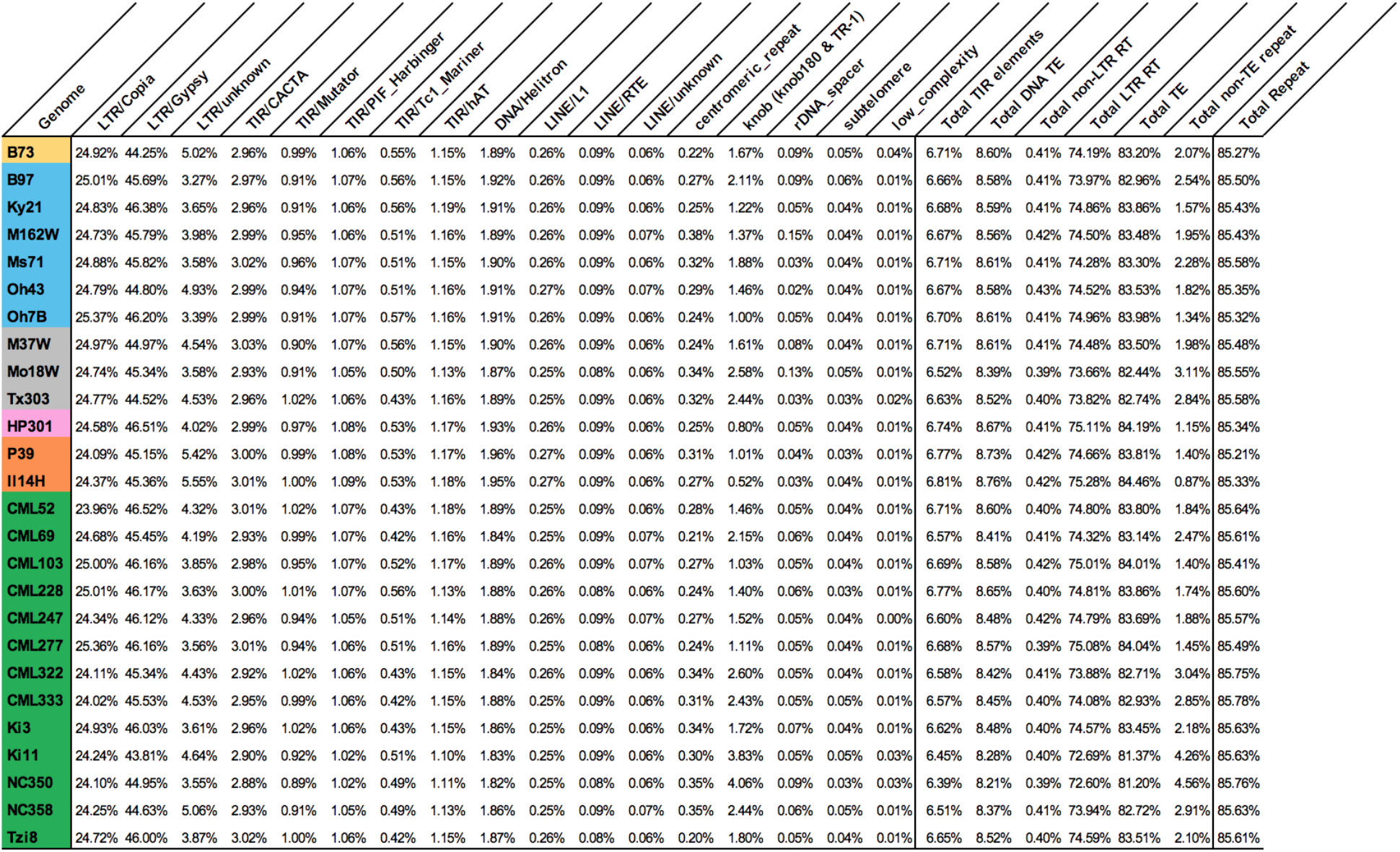
Percentage of repetitive sequences in NAM parent genomes.

**Table S5:**
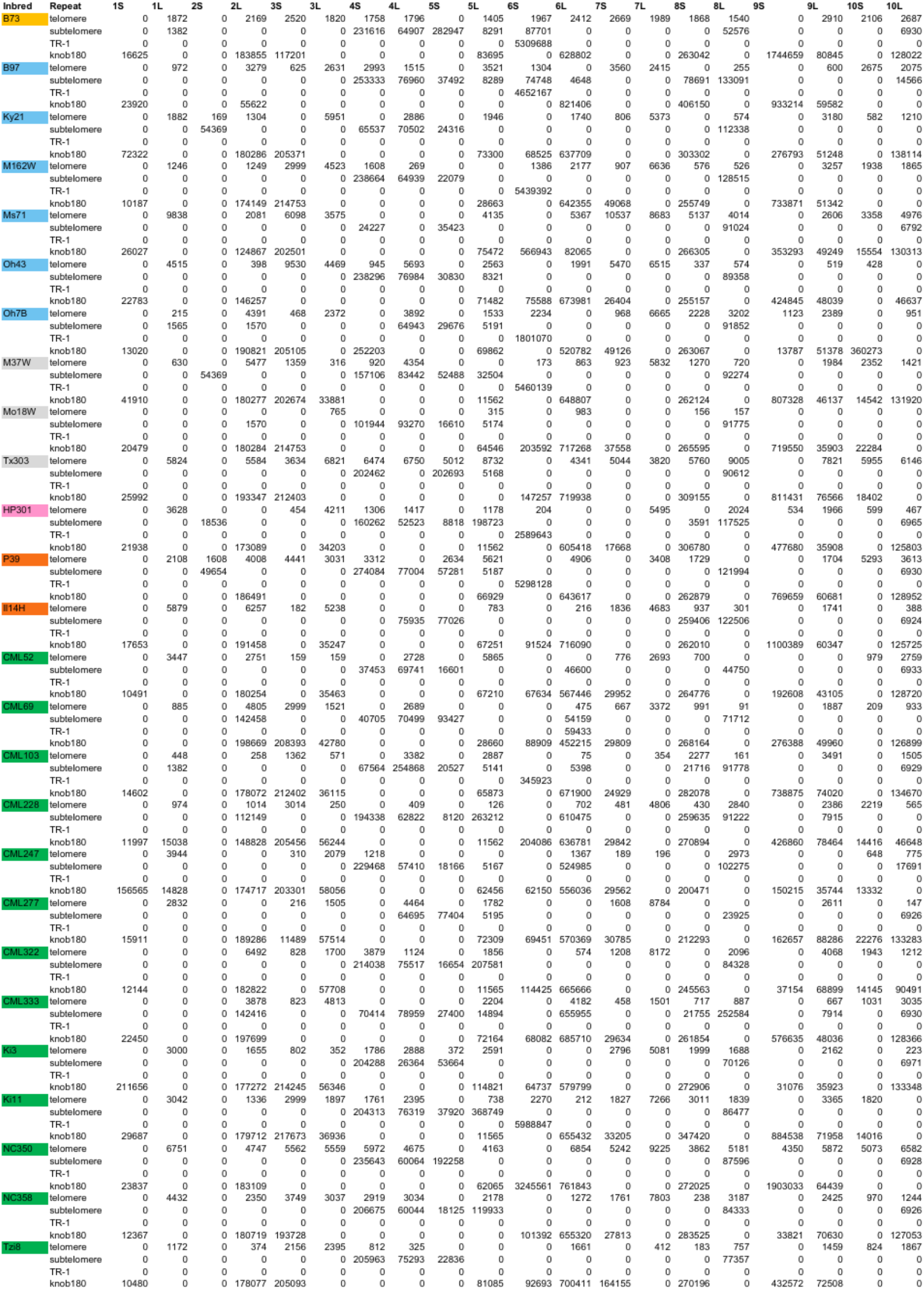
Characterization of assembly content of chromosome ends. Numbers are shown in bp.

**Table S6:** Extent of structural variation relative to B73 across the NAM assemblies including deletions (DEL), insertions (INS), inversions (INV), duplications (DUP), and translocations (TRA).

| nam_line | type | count | size(Mbp) | nam_line | type | count | size(Mbp) | nam_line | type | count | size(Mbp) | nam_line | type | count | size(Mbp) | nam_line | type | count |
| --- | --- | --- | --- | --- | --- | --- | --- | --- | --- | --- | --- | --- | --- | --- | --- | --- | --- | --- |
| B97 | DEL | 23533 | 267.27 | B97 | INS | 4249 | 1.69 | B97 | INV | 84 | 0.61 | B97 | DUP | 10 | 0.0061 | B97 | TRA | 950 |
| Ky21 | DEL | 22851 | 256.46 | Ky21 | INS | 4117 | 1.71 | Ky21 | INV | 69 | 0.97 | Ky21 | DUP | 4 | 0.0015 | Ky21 | TRA | 904 |
| M162W | DEL | 25060 | 278.52 | M162W | INS | 4635 | 1.85 | M162W | INV | 73 | 0.73 | M162W | DUP | 8 | 0.0030 | M162W | TRA | 1029 |
| Ms71 | DEL | 23974 | 270.59 | Ms71 | INS | 4329 | 1.74 | Ms71 | INV | 74 | 0.52 | Ms71 | DUP | 6 | 0.0025 | Ms71 | TRA | 992 |
| Oh43 | DEL | 22755 | 253.28 | Oh43 | INS | 3988 | 1.57 | Oh43 | INV | 82 | 1.09 | Oh43 | DUP | 6 | 0.0033 | Oh43 | TRA | 881 |
| Oh7B | DEL | 22243 | 241.38 | Oh7B | INS | 5923 | 5.72 | Oh7B | INV | 71 | 0.56 | Oh7B | DUP | 8 | 0.0025 | Oh7B | TRA | 909 |
| M37W | DEL | 24652 | 271.33 | M37W | INS | 4497 | 1.76 | M37W | INV | 74 | 0.60 | M37W | DUP | 6 | 0.0032 | M37W | TRA | 968 |
| Mo18W | DEL | 25811 | 286.33 | Mo18W | INS | 4577 | 1.78 | Mo18W | INV | 85 | 1.27 | Mo18W | DUP | 9 | 0.0033 | Mo18W | TRA | 1034 |
| Tx303 | DEL | 25311 | 277.33 | Tx303 | INS | 6612 | 6.87 | Tx303 | INV | 87 | 0.83 | Tx303 | DUP | 10 | 0.0033 | Tx303 | TRA | 999 |
| HP301 | DEL | 25033 | 273.35 | HP301 | INS | 6612 | 6.87 | HP301 | INV | 82 | 1.07 | HP301 | DUP | 8 | 0.0053 | HP301 | TRA | 946 |
| P39 | DEL | 25207 | 284.57 | P39 | INS | 4486 | 1.72 | P39 | INV | 86 | 1.08 | P39 | DUP | 7 | 0.0024 | P39 | TRA | 957 |
| II14H | DEL | 25141 | 286.65 | II14H | INS | 4238 | 1.69 | II14H | INV | 88 | 0.75 | II14H | DUP | 4 | 0.0021 | II14H | TRA | 970 |
| CML52 | DEL | 25534 | 276.54 | CML52 | INS | 6717 | 6.65 | CML52 | INV | 73 | 0.73 | CML52 | DUP | 5 | 0.0022 | CML52 | TRA | 1017 |
| CML69 | DEL | 25754 | 283.95 | CML69 | INS | 6637 | 6.52 | CML69 | INV | 68 | 0.32 | CML69 | DUP | 6 | 0.0021 | CML69 | TRA | 919 |
| CML103 | DEL | 25482 | 283.17 | CML103 | INS | 4468 | 1.73 | CML103 | INV | 87 | 0.82 | CML103 | DUP | 4 | 0.0014 | CML103 | TRA | 1024 |
| CML228 | DEL | 25969 | 281.35 | CML228 | INS | 6674 | 6.59 | CML228 | INV | 77 | 0.56 | CML228 | DUP | 5 | 0.0016 | CML228 | TRA | 991 |
| CML247 | DEL | 26361 | 288.40 | CML247 | INS | 6838 | 6.65 | CML247 | INV | 70 | 0.60 | CML247 | DUP | 13 | 0.0045 | CML247 | TRA | 941 |
| CML277 | DEL | 25663 | 280.11 | CML277 | INS | 6440 | 6.14 | CML277 | INV | 73 | 0.83 | CML277 | DUP | 9 | 0.0027 | CML277 | TRA | 968 |
| CML322 | DEL | 24665 | 267.92 | CML322 | INS | 6362 | 6.15 | CML322 | INV | 67 | 0.68 | CML322 | DUP | 11 | 0.0043 | CML322 | TRA | 974 |
| CML333 | DEL | 25061 | 275.46 | CML333 | INS | 4478 | 2.04 | CML333 | INV | 78 | 0.93 | CML333 | DUP | 13 | 0.0052 | CML333 | TRA | 943 |
| Ki3 | DEL | 25536 | 274.63 | Ki3 | INS | 6495 | 6.53 | Ki3 | INV | 86 | 0.76 | Ki3 | DUP | 8 | 0.0025 | Ki3 | TRA | 953 |
| Ki11 | DEL | 26223 | 287.32 | Ki11 | INS | 6816 | 7.17 | Ki11 | INV | 83 | 0.85 | Ki11 | DUP | 7 | 0.0029 | Ki11 | TRA | 994 |
| NC350 | DEL | 25807 | 284.24 | NC350 | INS | 6941 | 7.40 | NC350 | INV | 75 | 0.46 | NC350 | DUP | 13 | 0.0047 | NC350 | TRA | 1066 |
| NC358 | DEL | 25434 | 278.31 | NC358 | INS | 6613 | 6.71 | NC358 | INV | 77 | 0.90 | NC358 | DUP | 11 | 0.0041 | NC358 | TRA | 965 |
| Tzi8 | DEL | 25456 | 278.79 | Tzi8 | INS | 6461 | 6.23 | Tzi8 | INV | 73 | 0.41 | Tzi8 | DUP | 11 | 0.0036 | Tzi8 | TRA | 944 |

**Table S7:** Extent of structural variation across the major groups of NAM lines relative to B73. Mean/Median total sizes are shown in Mbp and mean/median sizes are shown in bp.

| Group | sv_type | mean_total_size(Mbp) | median_total_size(Mbp) | mean_total_count | median_total_count | mean_size | median_size |
| --- | --- | --- | --- | --- | --- | --- | --- |
| non-stiff-stalk | del | 261.25 | 261.86 | 23,402.67 | 23,192.00 | 11,163.25 | 11,291.11 |
| flints | del | 281.52 | 284.57 | 25,127.00 | 25,141.00 | 11,204.05 | 11,318.84 |
| tropical | del | 280.01 | 280.11 | 25,611.15 | 25,536.00 | 10,933.29 | 10,969.03 |
| mixed | del | 278.33 | 277.33 | 25,258.00 | 25,311.00 | 11,019.47 | 10,957.02 |
| non-stiff-stalk | ins | 2.38 | 1.72 | 4,540.17 | 4,289.00 | 523.97 | 402.10 |
| flints | ins | 3.43 | 1.72 | 5,112.00 | 4,486.00 | 670.21 | 384.17 |
| tropical | ins | 5.89 | 6.53 | 6,303.08 | 6,613.00 | 933.84 | 988.04 |
| mixed | ins | 3.47 | 1.78 | 5,228.67 | 4,577.00 | 663.80 | 389.32 |
| non-stiff-stalk | inv | 0.75 | 0.67 | 75.50 | 73.50 | 9,883.52 | 9,102.57 |
| flints | inv | 0.97 | 1.07 | 85.33 | 86.00 | 11,317.03 | 12,411.92 |
| tropical | inv | 0.68 | 0.73 | 75.92 | 75.00 | 8,967.27 | 9,768.79 |
| mixed | inv | 0.90 | 0.83 | 82.00 | 85.00 | 11,010.98 | 9,764.09 |
| non-stiff-stalk | dup | 0.0031 | 0.0027 | 7.00 | 7.00 | 446.86 | 390.93 |
| flints | dup | 0.0033 | 0.0024 | 6.33 | 7.00 | 515.32 | 344.14 |
| tropical | dup | 0.0032 | 0.0029 | 8.92 | 9.00 | 359.06 | 319.11 |
| mixed | dup | 0.0032 | 0.0033 | 8.33 | 9.00 | 389.52 | 361.22 |
| non-stiff-stalk | tra | . | . | 944.17 | 929.50 | . | . |
| flints | tra | . | . | 957.67 | 957.00 | . | . |
| tropical | tra | . | . | 976.85 | 968.00 | . | . |
| mixed | tra | . | . | 1,000.33 | 999.00 | . | . |

**Table S8:** Coordinates and copy number of the *rp1* tandem array on chromosome 10S. Coordinates are referenced to each of the individual genome assemblies.

| genome | rp1 alignment<br>locus start | rp1 alignment<br>locus stop | size | copy<br>number | # gaps in<br>rp1 locus |
| --- | --- | --- | --- | --- | --- |
| B73 | 2823128 | 3532004 | 708876 | 14 | 10 |
| B97 | 3454017 | 3917737 | 463720 | 14 | 0 |
| Ky21 | 4184592 | 4460903 | 276311 | 10 | 0 |
| M162W | 3214938 | 3531971 | 317033 | 11 | 1 |
| Ms71 | 3007141 | 3073115 | 65974 | 8 | 0 |
| Oh43 | 3847294 | 4311005 | 463711 | 12 | 0 |
| Oh7B | 3485246 | 3551229 | 65983 | 5 | 5 |
| M37W | 3089289 | 4252158 | 1162869 | 30 | 29 |
| Mo18W | 2870789 | 3095369 | 224580 | 7 | 0 |
| Tx303 | 2607068 | 3056758 | 449690 | 13 | 0 |
| HP301 | 3172164 | 3238132 | 65968 | 7 | 0 |
| P39 | 2746503 | 2880037 | 133534 | 4 | 0 |
| II14H | 3585827 | 4088413 | 502586 | 15 | 0 |
| CML52 | 2596259 | 2756085 | 159826 | 5 | 0 |
| CML69 | 3439600 | 3736711 | 297111 | 8 | 0 |
| CML103 | 3211757 | 4325999 | 1114242 | 23 | 36 |
| CML228 | 3114513 | 4067585 | 953072 | 20 | 20 |
| CML247 | 2762984 | 2858688 | 95704 | 7 | 0 |
| CML277 | 2707651 | 3004761 | 297110 | 7 | 0 |
| CML322 | 2950611 | 3216435 | 265824 | 10 | 0 |
| CML333 | 2819939 | 3359941 | 540002 | 20 | 0 |
| Ki3 | 2861923 | 3119062 | 257139 | 7 | 0 |
| Ki11 | 3192016 | 3738606 | 546590 | 17 | 2 |
| NC350 | 3699401 | 4027572 | 328171 | 15 | 0 |
| NC358 | 3225450 | 3761930 | 536480 | 19 | 0 |
| Tzi8 | 3730377 | 4252564 | 522187 | 17 | 4 |

**Table S9:** Number of significant GWAS SNPs (p ≤ 0.05 after FDR correction) for each trait within and outside of UMR intervals.

| Traits | Trait ID | Non-Genic UMRs | Genic UMRs | Non-Genic, Not in UMRs |
| --- | --- | --- | --- | --- |
| Days_To_Silk | T2 | 158,365 | 129,278 | 585,696 |
| ASI | T3 | 161,772 | 129,806 | 536,267 |
| Leaf_Length | T4 | 160,487 | 125,367 | 592,425 |
| Leaf_Width | T5 | 165,028 | 128,798 | 548,964 |
| Leaf_Angle | T7 | 167,842 | 131,599 | 548,246 |
| NLB_Index | T8 | 163,889 | 128,753 | 581,637 |
| Ear_Mass | T10 | 161,378 | 123,497 | 617,330 |
| Kernels_Per_Row | T12 | 158,128 | 127,793 | 590,508 |
| Twenty_Kernel_Weight | T15 | 164,603 | 128,236 | 556,810 |
| Tassel_Length | T16 | 164,883 | 130,534 | 580,184 |
| Spike_Length | T17 | 164,876 | 135,222 | 545,094 |
| Branch_Zone | T19 | 157,589 | 133,458 | 493,808 |
| Cob_Length | T20 | 155,803 | 126,585 | 527,054 |
| Cob_Diameter | T21 | 162,893 | 131,872 | 546,333 |
| Ear_Row_Number | T22 | 167,956 | 132,579 | 554,642 |
| GDD_Anth_Long | T23 | 150,234 | 130,778 | 591,526 |
| GDD_Anth_Short | T24 | 164,127 | 122,255 | 640,282 |
| GDD_Anth_Photo_Resp | T25 | 159,969 | 120,768 | 691,542 |
| GDD_Silk_Long | T26 | 163,678 | 130,114 | 580,472 |
| GDD_Silk_Short | T27 | 162,665 | 130,710 | 624,963 |
| GDD_Silk_Photo_Resp | T28 | 154,231 | 115,941 | 673,853 |
| SLB | T29 | 164,783 | 129,207 | 610,486 |
| Days_To_Anthesis | T30 | 159,054 | 130,507 | 577,097 |
| ChlorophyllA | T31 | 156,771 | 130,643 | 541,779 |
| ChlorophyllB | T32 | 172,509 | 119,504 | 701,721 |
| Malate | T33 | 164,180 | 133,006 | 540,370 |
| Fumarate | T34 | 165,037 | 126,789 | 651,454 |
| Fum2 | T35 | 173,892 | 124,777 | 657,173 |
| Glutamate | T36 | 157,213 | 138,957 | 580,812 |
| Amino_Acids | T37 | 167,341 | 132,337 | 572,995 |
| Protein | T38 | 162,263 | 128,924 | 687,760 |
| Nitrate | T39 | 158,667 | 140,264 | 629,072 |
| Starch | T40 | 160,254 | 133,203 | 613,835 |
| Sucrose | T41 | 158,926 | 129,010 | 609,554 |
| Glucose | T42 | 164,502 | 124,720 | 615,003 |
| Fructose | T43 | 162,445 | 125,856 | 637,147 |

## Supplementary Dataset

**Dataset S1**. Spreadsheet with data used for fractionation analysis. Data show the exon count matrix, genomic coordinates of regions syntenic with sorghum, and loci used for the GO analysis.

## Notes

https://github.com/HuffordLab/NAM-genomes

